# The functions of SARS-CoV-2 neutralizing and infection-enhancing antibodies in vitro and in mice and nonhuman primates

**DOI:** 10.1101/2020.12.31.424729

**Authors:** Dapeng Li, Robert J Edwards, Kartik Manne, David R. Martinez, Alexandra Schäfer, S. Munir Alam, Kevin Wiehe, Xiaozhi Lu, Robert Parks, Laura L. Sutherland, Thomas H. Oguin, Charlene McDanal, Lautaro G. Perez, Katayoun Mansouri, Sophie M. C. Gobeil, Katarzyna Janowska, Victoria Stalls, Megan Kopp, Fangping Cai, Esther Lee, Andrew Foulger, Giovanna E. Hernandez, Aja Sanzone, Kedamawit Tilahun, Chuancang Jiang, Longping V. Tse, Kevin W. Bock, Mahnaz Minai, Bianca M. Nagata, Kenneth Cronin, Victoria Gee-Lai, Margaret Deyton, Maggie Barr, Tarra Von Holle, Andrew N. Macintyre, Erica Stover, Jared Feldman, Blake M. Hauser, Timothy M. Caradonna, Trevor D. Scobey, Wes Rountree, Yunfei Wang, M. Anthony Moody, Derek W. Cain, C. Todd DeMarco, ThomasN. Denny, Christopher W. Woods, Elizabeth W. Petzold, Aaron G. Schmidt, I-Ting Teng, Tongqing Zhou, Peter D. Kwong, John R. Mascola, Barney S. Graham, Ian N. Moore, Robert Seder, Hanne Andersen, Mark G. Lewis, David C. Montefiori, Gregory D. Sempowski, Ralph S. Baric, Priyamvada Acharya, Barton F. Haynes, Kevin O. Saunders

## Abstract

SARS-CoV-2 neutralizing antibodies (NAbs) protect against COVID-19. A concern regarding SARS-CoV-2 antibodies is whether they mediate disease enhancement. Here, we isolated NAbs against the receptor-binding domain (RBD) and the N-terminal domain (NTD) of SARS-CoV-2 spike from individuals with acute or convalescent SARS-CoV-2 or a history of SARS-CoV-1 infection. Cryo-electron microscopy of RBD and NTD antibodies demonstrated function-specific modes of binding. Select RBD NAbs also demonstrated Fc receptor-γ (FcγR)-mediated enhancement of virus infection *in vitro*, while five non-neutralizing NTD antibodies mediated FcγR-independent *in vitro* infection enhancement. However, both types of infection-enhancing antibodies protected from SARS-CoV-2 replication in monkeys and mice. Nonetheless, three of 31 monkeys infused with enhancing antibodies had higher lung inflammation scores compared to controls. One monkey had alveolar edema and elevated bronchoalveolar lavage inflammatory cytokines. Thus, while *in vitro* antibody-enhanced infection does not necessarily herald enhanced infection *in vivo*, increased lung inflammation can occur in SARS-CoV-2 antibody-infused macaques.

## Introduction

The severe acute respiratory syndrome coronavirus 2 (SARS-CoV-2) has caused a global pandemic with over 96 million cases and 2.1 million deaths (https://coronavirus.jhu.edu). Development of combinations of neutralizing antibodies for prevention or treatment of infection may help to control the pandemic, while the ultimate solution to control the COVID-19 pandemic is a safe and effective vaccine (Graham, 2020; Sempowski et al., 2020).

Neutralizing antibodies (NAbs) that can block viral entry are crucial for controlling virus infections (Battles and McLellan, 2019; Corti and Lanzavecchia, 2013; Dashti et al., 2019). Previously reported SARS-CoV and MERS-CoV NAbs function by targeting the receptor-binding domain (RBD) or the N-terminal domain (NTD) of spike (S) protein to block receptor binding, or by binding to the S2 region of S protein to interfere with S2-mediated membrane fusion (Du and Jiang, 2010; Jiang et al., 2020; Xu et al., 2019). Importantly, prophylactic or therapeutic use of SARS-CoV-2 NAbs in non-human primates (Shi et al., 2020) or rodent models (Hassan et al., 2020; Rogers et al., 2020; Wu et al., 2020b) showed protection from SARS-CoV-2-induced lung inflammation and/or reduction in viral load. SARS-CoV-2 NAbs reported to date predominantly target the RBD region (Baum et al., 2020; Brouwer et al., 2020b; Cao et al., 2020; Hansen et al., 2020; Ju et al., 2020; Liu et al., 2020a; Pinto et al., 2020; Robbiani et al., 2020; Rogers et al., 2020; Shi et al., 2020; Wrapp et al., 2020a; Wu et al., 2020b). In contrast, NTD antibodies that neutralize SARS-CoV-2 mediate more modest neutralization potency (Brouwer et al., 2020a; Chi et al., 2020; Wec et al., 2020; Zost et al., 2020a; Zost et al., 2020b).

Antibody-dependent enhancement (ADE) of infection *in vitro* has been reported with vaccination for respiratory syncytial virus (RSV), with vaccination for dengue virus, or with dengue virus infection (Arvin et al., 2020). ADE is often mediated by Fc receptors for IgG (FcγRs), complement receptors (CRs) or both, and is most commonly observed in cells of monocyte/macrophage and B cell lineages (Iwasaki and Yang, 2020; Ubol and Halstead, 2010). *In vitro* studies have demonstrated FcγR-mediated ADE of SARS-CoV-1 infection of ACE2-negative cells (Jaume et al., 2011; Kam et al., 2007; Wan et al., 2020; Wang et al., 2014; Yilla et al., 2005; Yip et al., 2016; Yip et al., 2014). One group demonstrated FcγR-independent infection enhancement of SARS-CoV-1 in Vero cells, and isolated a monoclonal antibody that was suggested to induce enhanced lung viral load and pathology *in vivo* (Wang et al., 2016). The ability of antibodies that bind the SARS-CoV-2 S protein to mediate infection enhancement *in vivo* is unknown but is a theoretical concern for COVID-19 vaccine development (Arvin et al., 2020; Bournazos et al., 2020; Haynes et al., 2020; Iwasaki and Yang, 2020).

Here, we identified potent *in vitro* neutralizing RBD and NTD antibodies as well as *in vitro* infection-enhancing RBD and NTD antibodies from individuals infected with SARS-CoV-1 or SARS-CoV-2. Negative stain electron microscopy (NSEM) and cryo-electron microscopy (cryo-EM) revealed distinct binding patterns and the precise epitopes of infection-enhancing and neutralizing antibodies. *In vitro* studies using human FcγR-expressing or ACE2-expressing cell lines demonstrated that the RBD antibodies mediated FcγR-dependent infection enhancement, whereas the NTD antibodies induced FcγR-independent infection enhancement. However, using monkey and mouse models of SARS-CoV-2 infection, none of the *in vitro* infection-enhancing antibodies enhanced SARS-CoV-2 viral replication or infectious virus in the lung *in vivo*. Rather, three of 31 monkeys had lung pathology or bronchoalveolar lavage (BAL) cytokine levels suggestive of enhanced lung disease. Thus, *in vitro* infection enhancing activity of SARS-CoV-2 RBD and NTD antibodies controlled virus *in vivo* but was rarely associated with enhanced lung pathology.

## RESULTS

### Isolation of neutralizing and infection-enhancing SARS-CoV-2 antibodies

SARS-CoV-2-reactive human monoclonal antibodies from plasmablasts or SARS-CoV-2-reactive memory B cells were isolated by flow cytometric sorting (Liao et al., 2009; Liao et al., 2013) from a SARS-CoV-2 infected individual 11, 15 and 36 days post-onset of symptoms. Additional memory B cells were isolated from an individual infected with SARS-CoV-1 ∼17 years prior to sample collection (***Figures 1A-B, S1 and S2***). We isolated and characterized 1,737 antibodies that bound to SARS-CoV-2 S or nucleocapsid (NP) proteins (***Figure 1C; Table S1***). We selected 187 antibodies for further characterization, and examined neutralization of SARS-CoV-2 pseudovirus and replication-competent SARS-CoV-2. Forty-four of 81 recombinant RBD antibodies exhibited neutralization when assayed in 293T/ACE2 cell pseudovirus, SARS-CoV-2 microneutralization, or SARS-CoV-2 plaque reduction assays (***Figures S3A-F; Tables S2-S3***).

**Figure 1.**
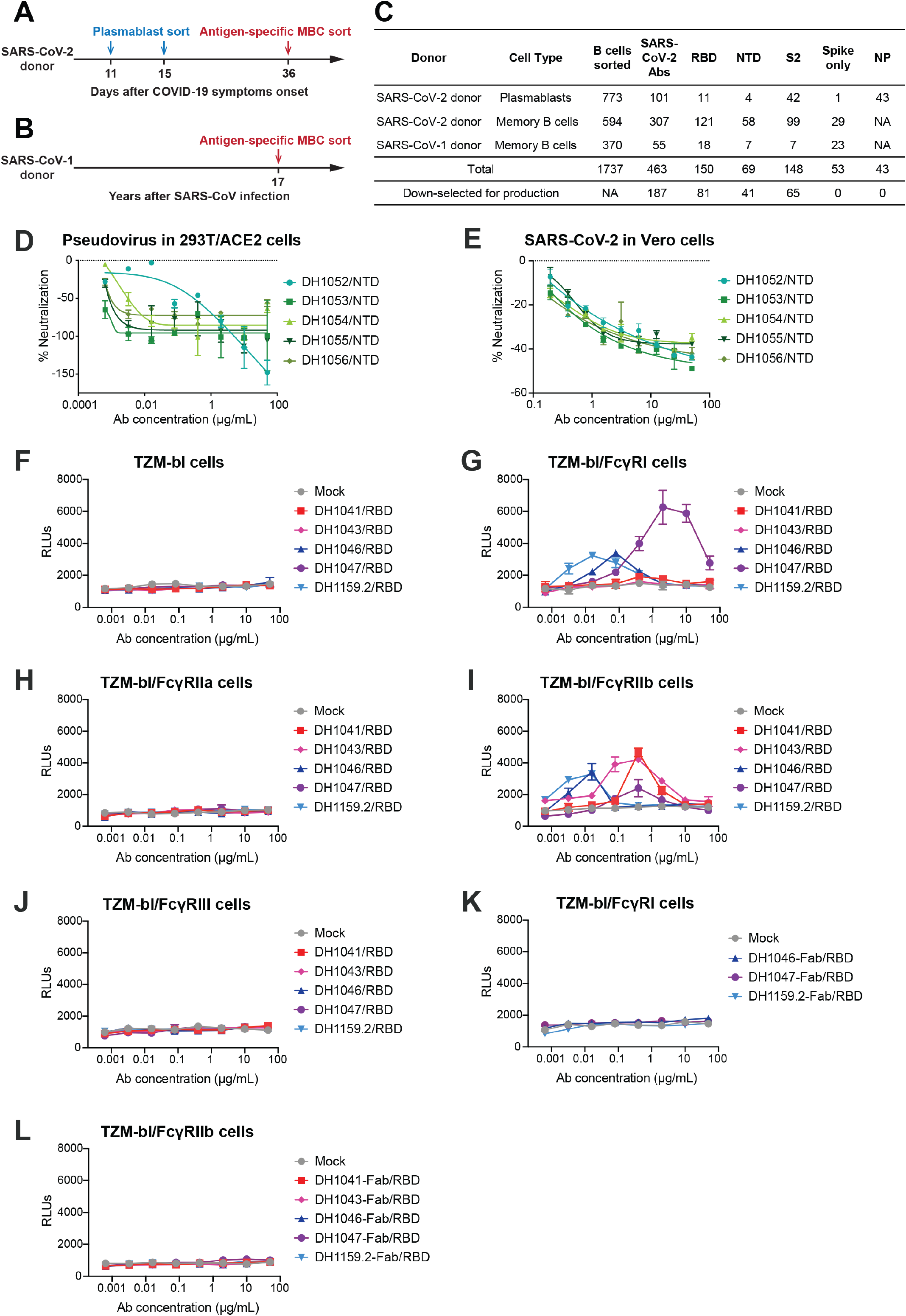
SARS-CoV-2 receptor-binding domain (RBD) and N-terminal domain (NTD) antibodies mediate FcγR-dependent and FcγR-independent enhancement of SARS-CoV-2 infection respectively. **(A-B)** Timeline of blood sampling and antibody isolation from convalescent SARS-CoV-2 and SARS-CoV-1 donors. Plasmablasts and/or antigen-specific memory B cells (MBC) were sorted from a (A) SARS-CoV-2 infected individual (SARS-CoV-2 donor) and a (B) 2003 SARS survivor (SARS-CoV-1 donor). **(C)** Summary of number and specificity of antibodies isolated from each donor. **(D-E)** FcγR-independent SARS-CoV-2 infection-enhancement mediated by non-neutralizing NTD antibodies. *In vitro* neutralization curves for NTD infection-enhancing antibodies against (D) pseudotyped SARS-CoV-2 D614G in 293T-hACE2 cells, and I replication-competent nano-luciferase (nLuc) SARS-CoV-2 in Vero cells. **(F-J)** FcγR-dependent SARS-CoV-2 infection-enhancement in ACE2-negative cells mediated by neutralizing RBD antibodies. Pseudotyped SARS-CoV-2 incubated with RBD antibodies or mock medium control were inoculated on (F) parental TZM-bl cells, and TZM-bl cells stably expressing human FcγR receptors (G) FcγRI, (H) FcγRIIa, (I) FcγRIIb or (J) FcγRIII. (**K-L**)The effect of RBD antibody fragment antigen-binding regions (Fabs) on pseudotyped SARS-CoV-2 D614G infection were tested in (K) FcγRI-expressing TZM-bl cells and (L) FcγRIIb-expressing TZM-bl cells. Relative luminescence units (RLUs) were measured in cell lysate at 68-72 hours post-infection. Upward deflection of RLUs in the presence of antibody indicates FcγR-mediated infection. Three or four independent experiments were performed and representative data are shown.

Ten of forty-one NTD antibodies neutralized in the 293T/ACE2 pseudovirus and plaque reduction assays, with the most potent antibody neutralizing pseudovirus with an IC_50_ of 39 ng/mL (***Figures S3G-I; Tables S4-S5***). In addition, 5 non-neutralizing NTD antibodies enhanced SARS-CoV-2 pseudovirus infection in 293T/ACE2 cells by 56% to 148% (***Figure 1D***). Infection of replication-competent SARS-CoV-2 nano-luciferase virus (Hou et al., 2020) also increased in the presence of each of the 5 non-neutralizing NTD antibodies (***Figure 1E***). Analysis of *in vitro* enhancement of NTD antibodies in ACE2-negative TZM-bl cells demonstrated no infection enhancement, demonstrating enhancement-dependence on spike-ACE2 engagement. Both ACE2-expressing 293T cells used for pseudovirus assays and Vero cells lack FcγR expression (Takada et al., 2007). Thus, NTD enhancement of SARS-CoV-2 infection was FcγR-independent.

Previous studies have demonstrated FcγR-mediated ADE of SARS-CoV-1 infection in ACE2-negative cells (Jaume et al., 2011; Kam et al., 2007; Wan et al., 2020; Wang et al., 2014; Yilla et al., 2005; Yip et al., 2016; Yip et al., 2014). Here, FcγR-dependent infection enhancement was determined by generating stable TZM-bl cell lines that expressed individual human FcγRs (FcγRI, FcγRIIa, FcγRIIb or FcγRIII). TZM-bl cells lack ACE2 and TMPRSS2 receptors, thus SARS-CoV-2 was unable to infect FcγR-negative TZM-bl cells (***Figure 1F***). One hundred S-reactive IgG1 antibodies selected from ***Tables S2-S7*** were tested for their ability to facilitate SARS-CoV-2 infection of TZM-bl cells. Three of the antibodies enabled SARS-CoV-2 infection of TZM-bl cells expressing FcγRI, and five antibodies mediated infection of FcγRIIb-expressing TZM-bl cells (***Figures 1F-J***). The antigen-binding fragments (Fabs) of these antibodies did not mediate infection enhancement of TZM-bl cells expressing FcγRI or FcγRIIb, demonstrating Fc-dependence of enhancement (***Figures 1K-L***). FcγRI and FcγRIIb-dependent infection-enhancing antibodies were specific for the RBD of S protein, consistent with recent results using COVID-19 patient sera or a recombinant antibody (Wu et al., 2020a). Thus, RBD antibodies can be either neutralizing in the 293T/ACE2 cell line, infection-enhancing in the TZM-bl-FcγR-expressing cell lines, or both (***Figure 2A***). NTD antibodies can either be neutralizing or infection-enhancing in the 293T/ACE2 cell line or Vero E6 cells (***Figure 2A***).

**Figure 2.**
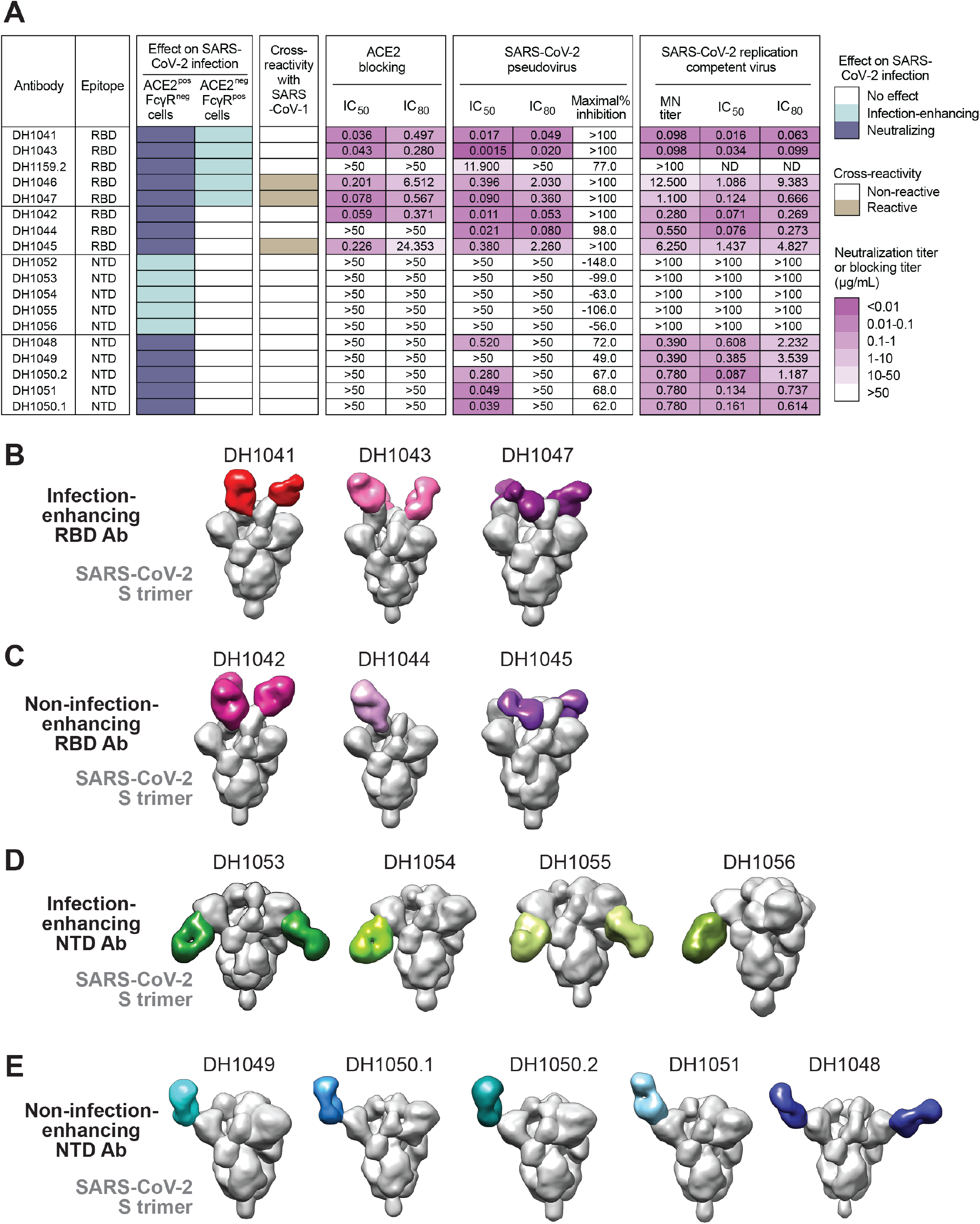
Structural and phenotypic characterization of infection-enhancing and non-infection-enhancing RBD and NTD antibodies. **(A)** Phenotypic summary of antibodies selected for in-depth characterization. Antibody functions are color-coded based on the key shown at the right. The heatmap denotes for each antibody the epitope location, neutralizing or infection-enhancing activity in ACE2-positive/FcγR-negative cells or ACE2-negative/FcγR-positive cells. Additionally, the heatmap indicates the ability of each antibody to bind to SARS-CoV-1 S protein by ELISA, the ability of each antibody to block ACE2 binding to SARS-CoV-2 S protein, and neutralization titers against SARS-CoV-2 pseudovirus and replication-competent virus. MN titer, micro-neutralization titer; ND, not determined. **(B-E)** 3D reconstruction of negative stain electron microscopy images of SARS-CoV-2 Spike ectodomain trimers stabilized with 2 proline mutations (S-2P; gray) bound to (B) infection-enhancing RBD antibody Fabs, (C) non-infection-enhancing RBD antibody Fabs, (D) infection-enhancing NTD antibody Fabs, (E) non-infection-enhancing NTD antibody Fabs. Fabs are pseudo-colored according to the phenotypic category to which they belong.

### Characterization of infection-enhancing Spike recombinant antibodies

We compared the phenotypes and binding modes to S protein for five infection-enhancing RBD antibodies and three RBD antibodies that lacked infection enhancement to elucidate differences between these two types of antibodies. The selected RBD antibodies neutralized SARS-CoV-2 pseudovirus and/or replication-competent virus in ACE2-expressing cells (***Figures 2A and S5***), despite five of these antibodies mediating infection enhancement in ACE2-negative, FcγR-positive TZM-bl cells (***Figures 1F-L, 2A, and S5***). Both types of selected RBD antibodies blocked ACE2 binding to S protein. Fabs of four of the infection-enhancing RBD antibodies and two of the non-infection-enhancing RBD antibodies bound to S with high affinities ranging from 0.1 to 9 nM (***Figures S6-8***). Thus, the infection-enhancing or non-enhancing RBD antibodies showed similarities in ACE2 blocking, affinity, and neutralization of ACE2-dependent SARS-CoV-2 infection (***Figure 2A***).

For six representative RBD antibodies, we obtained NSEM reconstructions of Fabs in complex with stabilized S ectodomain trimer. Infection-enhancing RBD antibodies DH1041 and DH1043 bound with a vertical approach (***Figure 2B***), parallel to the central axis of the S trimer, similar to non-infection-enhancing antibodies DH1042 and DH1044 (***Figure 2C***). The epitopes of antibodies DH1041, DH1042, and DH1043 overlapped with that of the ACE-2 receptor (Wang et al., 2020), consistent with their ability to block ACE-2 binding to S protein (***Figures 2A and S10***). Their epitopes were similar to those of three previously described antibodies, P2B-2F6 (Ju et al., 2020), H11-H4, and H11-D4 (***Figure S10***) (Huo et al., 2020; Zhou et al., 2020). The epitope of another non-infection-enhancing RBD antibody DH1044 was only slightly shifted relative to DH1041, DH1042 and DH1043 (***Figure 2C***), but resulted in DH1044 not blocking ACE2 binding (***Figures 2A and S10***). The remaining two RBD antibodies, DH1045 and DH1047, cross-reacted with both SARS-CoV-1 and SARS-CoV-2 S (***Figures 2A, S4, S31***). DH1047 also reacted with bat and pangolin CoV spike proteins (***Figures 2A, S4, S31***). Although DH1047 mediated FcγR-dependent infection of TZM-bl cells and DH1045 did not, both antibodies bound to RBD-up S conformations with a more horizontal angle of approach (***Figures 2B-C and S10***) (Pak et al., 2009). Thus, epitopes and binding angles of RBD antibodies determined by NSEM did not discriminate between antibodies that mediated FcγR-dependent infection enhancement and those that did not.

We performed NSEM analyses of five FcγR-independent, infection-enhancing NTD antibodies, and five non-infection-enhancing NTD antibodies (DH1048, DH1049, DH1051, DH1050.1 and DH1050.2) that neutralized SARS-CoV-2 (pseudovirus IC_50_ titers 39 - 520 ng/mL; SARS-CoV-2 plaque reduction IC_50_ titers 390 - 780 ng/mL) (***Figures 2A and S5C-D***). The Fabs of neutralizing antibodies DH1050.1 and DH1051 bound to stabilized S ectodomain with affinities of 16 and 19 nM respectively, whereas the infection-enhancing antibody DH1052 bound with 294 nM affinity (***Figures S6-8***). NSEM reconstructions obtained for nine of the ten NTD antibodies showed that the FcγR-independent, infection-enhancing NTD antibodies (DH1053-DH1056) bound to S with their Fab constant domains directed downward toward the virus membrane (***Figure 2D)***, whereas the five neutralizing NTD-directed Abs (DH1048-DH1051) bound to S with the constant domain of the Fab directed upward away from the virus membrane (***Figure 2E)***. Thus, S protein antibody epitopes and binding modes were associated with FcγR-independent, infection-enhancing activity of NTD antibodies. The five neutralizing antibodies bound the same epitope as antibody 4A8 (Wrapp et al., 2020a), with three of the five having the same angle of approach as 4A8 (***Figure S11***). Interestingly, the NTD antibodies with the same angle of approach as 4A8, were also genetically similar to 4A8, being derived from the same V_H_1-24 gene segment (***Table S8***), although their light chains were different (Wrapp et al., 2020a). These NTD antibodies may constitute a neutralizing antibody class that can be elicited in multiple individuals.

### Competition between infection-enhancing and non-infection enhancing antibodies

To determine whether infection-enhancing antibodies could compete with non-infection-enhancing antibodies for binding to S ectodomain, we performed surface plasmon resonance (SPR) competitive binding assays. RBD antibodies segregated into two clusters, where antibodies within a cluster blocked each other and antibodies in different clusters did not block each other (***Figures 3A and S12***). One cluster included antibodies DH1041, DH1043 and DH1044, and the other cluster included antibodies DH1046 and DH1047. NSEM reconstructions showed DH1041 and DH1047 Fabs bound simultaneously to different epitopes of the stabilized S trimer (***Figure 3B***). Similarly, DH1043 and DH1047 Fabs also bound simultaneously to different epitopes on the stabilized S protein (***Figure 3B***).

**Figure 3.**
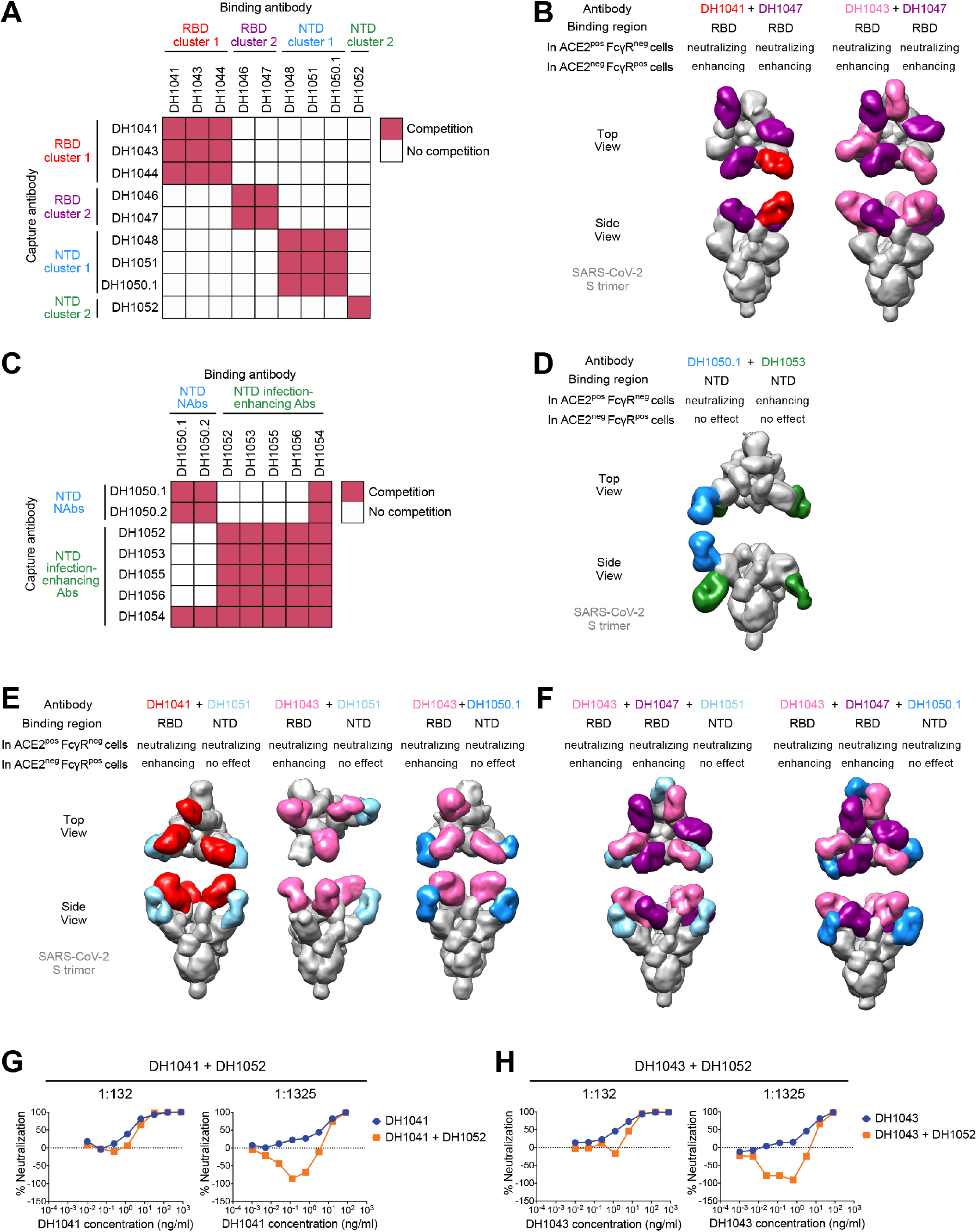
Biophysical and structural determination that infection-enhancing and non-infection enhancing antibodies can simultaneously bind to the same S protein. **(A)** Cross-blocking activity of RBD and NTD neutralizing antibodies tested by surface plasmon resonance (SPR). Soluble, stabilized SARS-CoV-2 S trimer (S-2P) was captured by the antibody on the Y-axis followed by binding by the antibody on the X-axis. Antibody binding was considered competitive (red squares) if the binding antibody did not react with the captured S protein. **(B)** 3D reconstruction of simultaneous recognition of SARS-CoV-2 S-2P trimer by two RBD antibodies DH1041+DH1047, or DH1043+DH1047. All three antibodies are SARS-CoV-2 infection-enhancing in ACE2-negative/FcγR-positive cells, but neutralizing in ACE2-positive/FcγR-negative cells. **(C)** Cross-blocking activity of neutralizing antibodies and infection-enhancing NTD antibodies tested by SPR. SARS-CoV-2 S-2P trimer was captured by the antibody on the Y-axis followed by binding by the antibody on the X-axis. Antibody binding was considered competitive (red squares) if the binding antibody did not react with the captured S protein. **(D)** 3D reconstruction of by NTD antibodies DH1053 and DH1050.1 simultaneously bound to SARS-CoV-2 S trimer protein. In ACE2-positive/FcγR-negative cells, DH1050.1 is neutralizing while DH1053 enhances SARS-CoV-2 infection. Both antibodies have no effect in ACE2-negative/FcγR-positive cells. **(E)** 3D reconstruction of SARS-CoV-2 S protein simultaneously bound to a RBD infection-enhancing antibody and a NTD non-infection-enhancing antibody. All of these antibodies neutralize SARS-CoV-2 infection of ACE2-positive/FcγR-negative cells. **(F)** 3D reconstruction of SARS-CoV-2 S protein bound to triple-antibody combinations of RBD antibody DH1043, RBD antibody DH1047, and either NTD antibody DH1051 (left) or DH1050.1 (right). Both RBD antibodies enhance SARS-CoV-2 infection of ACE2-negative/FcγR-positive cells, but neutralize infection of ACE2-positive/FcγR-negative cells. NTD antibodies DH1051 and DH1050.1 neutralize SARS-CoV-2 infection of ACE2-negative/FcγR-positive cells, but have no effect on infection of ACE2-negative/FcγR-positive cells. **(G-H)** RBD antibody neutralization of SARS-CoV-2 pseudovirus infection of ACE2-expressing cells in the presence of infection-enhancing NTD antibody DH1052. The infection-enhancing NTD antibody DH1052 was mixed with RBD antibodies DH1041 **(G)** or DH1043 **(H)** in 1:132 ratio or 1:1,325 ratio, respectively. Serial dilutions of the NTD:RBD antibody mixtures (orange), as well as RBD antibody alone (blue) were examined for neutralization of SARS-CoV-2 D614G pseudovirus infection of 293T/ACE2 cells.

NTD antibodies also segregated into two clusters based on their ability to block each other (***Figure 3A***). Neutralizing NTD antibodies blocked each other and formed one cluster, while infection-enhancing/non-neutralizing NTD antibodies blocked each other and formed a second cluster (***Figures 3A, 3C, S12 and S13***). NSEM reconstruction of SARS-CoV-2 S trimer bound with Fabs of neutralizing NTD antibody DH1050.1 and infection-enhancing NTD antibody DH1052 confirmed that the two antibodies could simultaneously bind to distinct epitopes on a single SARS-CoV-2 S trimer (***Figure 3D***). DH1054 was unique as it was able to block both infection-enhancing and neutralizing NTD antibodies (***Figures 3C and S13***).

NTD antibodies did not compete with RBD antibodies for binding to S trimer (***Figure 3A***). This result gave rise to the notion that in a polyclonal mixture of antibodies, the SARS-CoV-2 S trimer could bind both RBD and NTD antibodies. To determine the potential for this complex to form, we liganded SARS-CoV-2 S trimer with Fabs of each type of antibody and visualized the complex using NSEM. NSEM showed that neutralizing RBD antibodies could also bind to the same S protomer as neutralizing NTD antibodies DH1050.1 or DH1051 (***Figure 3E***). Moreover, we found that a single S protomer could be simultaneously occupied by two RBD antibodies (DH1043 and DH1047) and an NTD antibody (DH1050.1) (***Figure 3F***). Thus, the S trimer could simultaneously bind to multiple RBD and NTD neutralizing antibody Fabs.

### FcγR-independent infection-enhancement in the presence of neutralizing antibodies

The NSEM determination of antibody binding modes demonstrated that certain infection-enhancing antibodies and non-infection enhancing antibodies bound to distinct epitopes on the same S protomer (***Figures 3A-F***). Thus, we determined the functional outcome of infection-enhancing antibodies binding to S in the presence of neutralizing antibodies. We examined whether a FcγR-independent, infection-enhancing NTD antibody could inhibit RBD antibody neutralization of ACE-2-mediated SARS-CoV-2 pseudovirus infection of 293T/ACE2 cells *in vitro*. We hypothesized that the outcome would be dependent on which antibody was present at the highest concentration. We examined RBD antibody DH1041 neutralization in the presence of 132 and 1,325-fold excess of infection-enhancing NTD antibody DH1052. DH1041 neutralization activity was minimally decreased in the presence of 132-fold excess of DH1052. When DH1041 neutralization was assessed in the presence of 1,325-fold excess of antibody DH1052, infection enhancement was observed when DH1041 concentration was below 10 ng/mL (***Figures 3G and S14***). A nearly identical result was obtained when we examined neutralization by DH1043 (***Figures 3H and S14***). Thus, a ∼1000-fold excess of infection-enhancing NTD antibody was required to out-compete the effect of a potent RBD neutralizing antibody *in vitro*.

### Cryo-EM structural determination of RBD and NTD-directed antibody epitopes

To visualize atomic level details of their interactions with the S protein, we selected representatives from the panels of RBD and NTD-directed antibodies for structural determination by cryo-EM. Of the RBD-directed antibodies, we selected two (DH1041 and DH1043) that most potently neutralized SARS-CoV-2 virus in the 293T/ACE2 pseudovirus assay and also enhanced infection in TZM-bl-FcγRI or -FcγRIIb cells. We also selected the RBD-directed antibody DH1047, which shared the infection enhancing and ACE-2 blocking properties of DH1041 and DH1043, but unlike DH1041 and DH1043, DH1047 also showed reactivity with the SARS-CoV-1, bat SHC014-CoV, RaTG13-CoV and pangolin GX-P4L-CoV S proteins (***Figure S4A and S29***). From the panel of NTD-directed antibodies, we selected one infection-enhancing NTD antibody, DH1052, and one neutralizing NTD antibody, DH1050.1, for higher resolution structural determination by cryo-EM. The stabilized SARS-CoV-2 S ectodomain “2P” (S-2P) (Wrapp et al., 2020b) was used for preparing complexes for structural determination.

For all three RBD-directed antibodies, the cryo-EM datasets revealed heterogeneous populations of S protein with at least one RBD in the “up” position (***Figures 4, S15 and S16***). We did not find any unliganded S or any 3-RBD-down S population, even though the unliganded S-2P consistently shows a 1:1 ratio of 1-RBD-up and 3-RBD-down populations (Henderson et al., 2020; Walls et al., 2020), suggesting that binding of the RBD-directed antibodies to S protein alters RBD dynamics. All S-2P trimers were stoichiometrically bound to three Fabs, with antibodies bound to both up and down RBDs in an S-2P trimer. We observed considerable disorder in the bound antibodies, primarily due to RBD motion, which was partially resolved by 3D classification, following which we identified the least disordered Fab for each complex by visual examination of the cryo-EM reconstructions for model building. Although the resolutions in regions of interest around the RBD-Fab binding interface were poorer than the overall reported resolutions, the cryo-EM reconstructions in these regions were resolved enough to obtain unambiguous definition of the binding sites.

**Figure 4.**
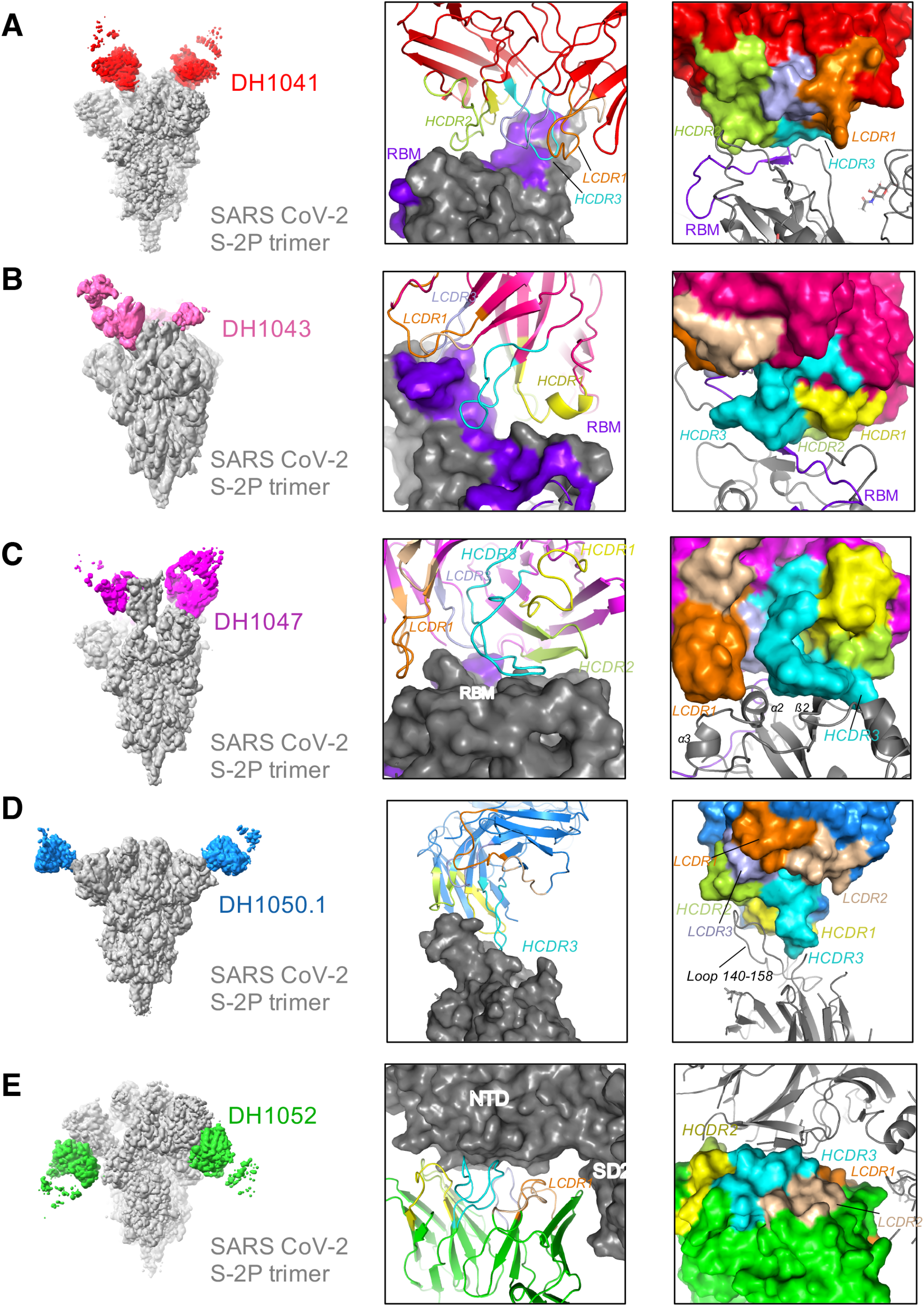
Cryo-electron microscopy of neutralizing and non-neutralizing antibodies in complex with SARS-CoV-2 Spike ectodomain. Structures of SARS-CoV-2 S protein in complex with RBD antibodies (**A)** DH1041 (red), (**B)** DH1043 (pink), (**C)** DH1047 (magenta), (**D)** neutralizing NTD antibody DH1050.1 (blue), and (**E)** infection-enhancing NTD antibody DH1052 (green). Each antibody is bound to Spike ectodomain stabilized with 2 proline mutations (S-2P) shown in gray with its Receptor Binding Motif (RBM) colored purple blue. (Right) Zoomed-in views of the antibody interactions with S-2P trimers. The antibody complementarity determining (CDR) loops are colored: HCDR1 yellow, HCDR2 limon, HCDR3 cyan, LCDR1 orange, LCDR2 wheat and LCDR3 light blue.

We observed that the primary epitopes of DH1041 and DH1043 were centered on the Receptor Binding Motif (RBM; residues 483-506) of the RBD (***Figures 4A-B, S17 and S18***), providing structural basis for the ACE-2 blocking phenotype of these antibodies. While DH1041 utilized its heavy chain complementarity determining regions (CDRs) to contact the RBM, the DH1043 paratope included both its heavy and light chains.

Unlike the RBM-focused epitope of DH1041 and DH1043, the epitope of antibody DH1047 was focused around the α2 and α3 helices and β2 strand that are located outside the N-terminus of the RBM (***Figures 4C and S19***) (Ju et al., 2020). Contact is also made by RBD residues 500-506 that are also outside the RBM but at its C-terminal end, and stack against the N-terminal end of the α3 helix providing a continuous interactive surface for DH1047 binding. The DH1047 paratope included heavy chain complementarity determining regions HCDR2, HCDR3 and light chain LCDR1 and LCDR3. The HCDR3 stacks against and interacts with the residues in the ß2 strand. Interactions with the ß2 strand are also mediated by HCDR2. Similar to DH1041 and DH1043, the DH1047 interacted with an “up” RBD conformation from an adjacent protomer although these interactions were not well-characterized due to disorder in that region.

We next determined cryo-EM structures of the NTD-directed neutralizing antibodies, DH1050.1 (***Figure 4D***) and NTD-directed infection-enhancing antibody, DH1052 (***Figure 4E***), at 3.4 Å and 3.0 Å resolutions, respectively. Unlike the RBD-directed antibodies DH1041, DH1043 and DH1047, where we only observed spikes with at least one RBD in the up position, the cryo-EM datasets of DH1050.1- and DH1052-bound complexes showed antibody bound to both 3-RBD-down and 1-RBD-up S-2P spikes (***Figures S20-S21***). Consistent with the NSEM reconstructions, the neutralizing antibody DH1050.1 and the non-neutralizing, infection-enhancing antibody DH1052 bound opposite faces of the NTD, with the epitope for the neutralizing antibody DH1050.1 facing the host cell membrane and the epitope for the non-neutralizing, infection-enhancing antibody DH1052 facing the viral membrane. The dominant contribution to the DH1050.1 epitope came from NTD loop region 140-158 that stacks against the antibody HCDR3 and extends farther into a cleft formed at the interface of the DH1050.1 HCDR1, HCDR2 and HCDR3 loops. The previously described NTD antibody 4A8 interacts with the same epitope in a similar manner, with its elongated HCDR3 dominating interactions and making similar contacts as were observed in the DH1050.1 complex, although DH1050.1 and 4A8 (Robbiani et al., 2020) show a rotation relative to each other about the stacked HCDR3 and NTD 140-158 loops, suggesting focused recognition of the elongated NTD loop by a class of antibodies sharing the same VH origin. The light chains of DH1050.1 and 4A8 do not contact the S protein, which is consistent with their diverse light chain gene origins (***Figures 4E and S21)***.

The infection enhancing NTD-directed antibody DH1052 bound the NTD at an epitope facing the viral membrane and comprised of residues spanning 27-32, 59-62 and 211-218, with all the CDR loops of both heavy and light chains involved in contacts with the NTD. We also observed contact of the antibody with the glycan at position 603, as well as the conformationally invariant SD2 region.

Thus, we found that the RBD-directed antibodies isolated in this study influenced RBD dynamics and bound only to spike with at least one RBD in the up conformations, and in some cases, also induced the 2-RBD-up and 3-RBD-up spike conformations. In contrast, the NTD-directed antibodies bound to both the 3-RBD-down and 1-RBD-up spikes that are present in the unliganded S-2P. Our results provide a structural explanation for the ACE-2 blocking activity of RBD-directed antibodies as well as for the cross-reactivity of the DH1047 antibody. We observed two distinct orientations for the NTD-directed antibodies that are either neutralizing or non-neutralizing, with the former binding the NTD surface that faces away from the viral membrane and the latter binding the surface that faces the viral membrane.

### Effect of NTD enhancing antibody DH1052 in mouse and macaque models of SARS-CoV-2 infection

Next, we assessed the effect of NTD infection enhancing Ab DH1052 in a COVID-19 disease mouse model of aged BALB/c mice challenged with the mouse-adapted SARS-CoV-2 MA10 strain (Leist et al., 2020a). The FcγR-independent, *in vitro* infection-enhancing antibody DH1052 or a control influenza antibody CH65 were given 12 hours prior to SARS-CoV-2 MA10 infection (***Figure 5A***). Throughout the four days of infection, DH1052-infused mice exhibited similar levels of body weight loss and higher survival than mice given negative control IgG (2/9 control mice died while 0/10 DH1052-treated mice died) (***Figures 5B-C***). In addition, DH1052-treated mice exhibited lower lung hemorrhagic scores, lower lung viral plaque-forming unit (PFU) titers and lower lung tissue subgenomic RNA levels compared to control IgG-infused mice (***Figures 5D-F***). Therefore DH1052 treatment resulted in less severe disease and reduced viral replication rather than infection enhancement. Thus, NTD antibodies that enhanced infection *in vitro* did not enhance infection *in vivo* in the SARS-CoV-2 MA10 virus infection mouse model. To determine whether DH1052 could have mediated protection by non-neutralizing, FcR-mediated functions, we determined the ability of DH1052 to interact with mouse FcγRI, IIb, III and IV and found DH1052 reacted primarily with FcγRI and FcγRIV (***Figure S22***). Thus, it is plausible that non-neutralizing antibody DH1052 may have protected mice with FcγR-dependent anti-SARS**-**CoV-2 mechanisms.

**Figure 5.**
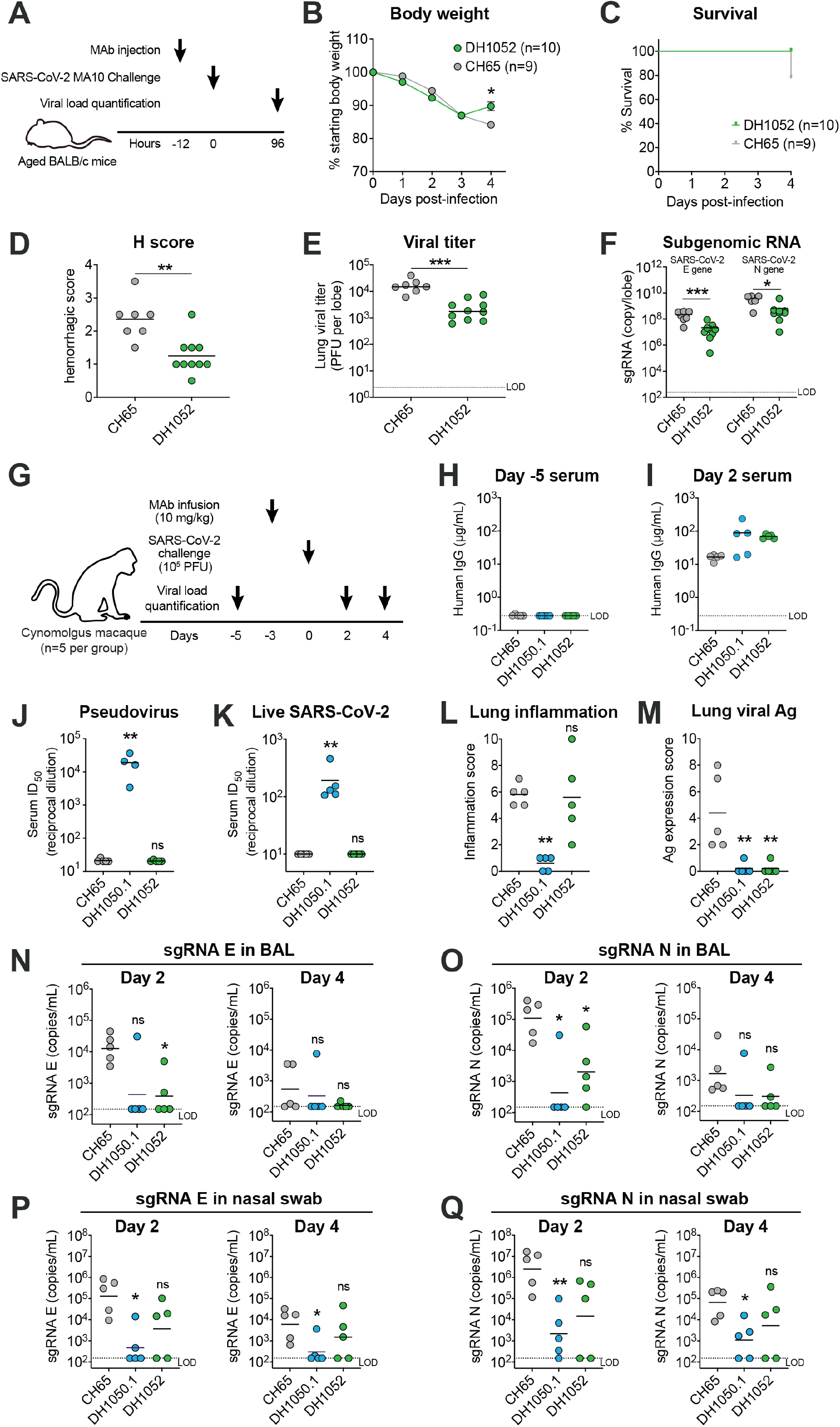
NTD antibody DH1052 enhances SARS-CoV-2 infection *in vitro*, but does not always enhance SARS-CoV-2 replication or disease *in vivo*. **(A-F)** DH1052 passive immunization and murine SARS-CoV-2 challenge study design and outcome. (**A**) Diagram of the study design showing 52 week old female BALB/c mice were i.p. injected with DH1052 (200 µg/mouse, n=10, green symbols) or CH65 control antibody (gray, 200 µg/mouse, n=9, gray symbols). After 12 hours, mice were challenged with 1×10^4 PFU of mouse-adapted SARS-CoV-2 MA10 virus. Mice were euthanized and tissues were harvested 96 hours post-infection. (**B**) Body weight and (**C**) survival were monitored daily. (**D**) Hemorrhagic scores, (**E**) lung viral titers, as well as (**F**) viral subgenomic RNA (sgRNA) for both SARS-CoV-2 envelope (E) and nucleocapsid (N) gene were measured 96 hours post-infection. **(G-Q)** Reduction of SARS-CoV-2 replication and disease in cynomolgus macaques by prophylactic administration of an NTD neutralizing antibody DH1050.1 or an in vitro infection-enhancing NTD Ab DH1052. **(G)** Diagram of the macaque study design showing cynomolgus macaques (n=5 per group) were infused with DH1052, DH1050.1 or an irrelevant control CH65 antibody 3 days before 10^5^ PFU of SARS-CoV-2 challenge via intranasal and intratracheal routes. Viral load including viral RNA and subgenomic RNA (sgRNA) were measured at the indicated pre-challenge and post-challenge timepoints. Lungs were harvested on Day 4 post-challenge for histopathology analysis. **(H-I)** Serum human IgG concentrations at (**H**) Day -5 and (**I**) Day 2. **(J-K)** Day 2 serum neutralization titers shown as the reciprocal serum dilution that inhibits 50% (ID_50_) of (**J**) pseudotyped SARS-CoV-2 replication in 293T/ACE2 cells or (**K**) SARS-CoV-2 replication in Vero cells. **(L-M)** Lung histopathology. Sections of the left caudal (Lc), right middle (Rm), and right caudal (Rc) lung were evaluated and scored for the presence of (**L**) inflammation by hematoxylin and eosin (H&E) staining, and for the presence of (**M**) SARS-CoV-2 nucleocapsid by immunohistochemistry (IHC) staining. Symbols indicate the sums of Lc, Rm, and Rc scores in each animal. **(N-O)** SARS-CoV-2 (**N**) E gene sgRNA and (**O**) N gene sgRNA in bronchoaveolar lavage (BAL) on Day 2 and Day 4 post challenge. **(P-Q)** SARS-CoV-2 (**P**) E gene sgRNA and (**Q**) N gene sgRNA in nasal swab on Day 2 and Day 4 post challenge. LOD, limit of detection. Statistical significance in all the panels were determined using Wilcoxon rank sum exact test. Asterisks show the statistical significance between indicated group and CH65 control group: ns, not significant, *P<0.05, **P<0.01, ***P<0.001.

We next examined the effect of infusion of NTD enhancing antibody DH1052, NTD neutralizing antibody DH1050.1 and control antibody CH65 on SARS-CoV-2 infection in monkeys (Leist et al., 2020b; Rockx et al., 2020). Cynomolgus macaques were infused with 10 mg of antibody per kg body weight and then challenged intranasally and intratracheally with 10^5^ plaque forming units of SARS-CoV-2 three days later (***Figure 5G***) (54). Antibody infusion resulted in human antibody concentrations ranging from 11 to 238 μg/mL in serum at day 2 post-challenge (***Figures 5H-I and S23A-D***). Sera with DH1050.1 neutralized SARS-CoV-2 pseudovirus at a mean ID_50_ titer of 19 (***Figure 5J***), and neutralized SARS-CoV-2 replication-competent virus at a mean ID_50_ titer of 192 (***Figure 5K***). In contrast, the presence of DH1052 or control antibody CH65 in serum did not neutralize SARS-CoV-2 (***Figures 5J-K***). Four of 5 macaques that received DH1052 had comparable lung inflammation to control CH65-infused macaques four days after infection (***Figures 5L, S24 and S25A***). However, one macaque (BB536A) administered DH1052 showed increased perivascular mononuclear inflammation, perivascular and alveolar edema (***Figure S25B***), and multiple upregulated bronchoalveolar fluid (BAL) cytokines (***Figures S26-S27***) compared to either control antibody-infused animals or the four other monkeys in the DH1052-treated group. Immunohistologic analysis with markers of macrophage subsets demonstrated alveolar and perivascular infiltration of M2-type macrophages in both monkey BB536A with histologic appearance of alveolar edema and in control antibody-treated monkey BB785E (***Figure S28***).

In contrast, macaques administered DH1050.1, a neutralizing NTD antibody, had lower lung inflammation than CH65-infused macaques (***Figures 5L, S24 and S25A***) and fewer infiltrating macrophages (***Figure S28***). Infusion of either neutralizing NTD DH1050.1 or *in vitro* infection-enhancing antibody DH1052 reduced viral nucleocapsid antigen in the lung (***Figures 5M, S24 and S25A)***. Envelope (E) gene subgenomic RNA (sgRNA) and nucleocapsid (N) gene sgRNA in the BAL were also reduced in macaques that were administered DH1050.1 or DH1052 compared to macaques treated with negative control antibody (***Figures 5N-O and S23G***). In nasal swab fluid, macaques showed reduced E and N gene sgRNA when neutralizing antibody DH1050.1 was infused (*p* < 0.05, nonparametric exact Wilcoxon test) (***Figures 5P-Q, S23E-F and S23H-I***). With DH1052 infusion, there was a trend to viral control but not significant.

Since DH1052-mediated infection enhancement *in vitro* increased as the antibody concentration increased (***Figures 1D-E***), we performed an additional challenge study in 6 additional cynomolgus macaques with either 30 mg/kg of DH1052 or CH65 control antibody (***Figure S29A***). After challenge, the infection-enhancing, non-neutralizing NTD antibody DH1052 again showed a trend to suppress BAL viral load (***Figures S29B-D***), and significantly reduced viral replication in nasal swab samples (*p* < 0.05, nonparametric exact Wilcoxon test) (***Figures S29E-G***) compared to the negative control antibody CH65. Moreover, DH1052-treated macaques showed equivalent lung inflammation (***Figures S29H-I***), antigen expression (***Figures S29J-K***), and cytokine expression (***Figure S30***) compared to CH65 control-treated macaques. Thus, with high dose (30mg/kg) of DH1052 antibody, there was no enhanced lung pathology or elevated BAL cytokine levels post-SARS-CoV-2 challenge. These results raised the hypothesis that the lung pathology seen in monkey BB536A was rare and may not have been caused by antibody infusion.

### FcγR-dependent, *in vitro* infection-enhancing RBD antibodies do not enhance SARS-CoV-2 infection in mice

We next tested RBD neutralizing antibodies that also mediated infection enhancement in TZM-bl cells expressing FcγRI or FcγRII in a SARS-CoV-2 acquisition mouse model (***Figures 6A-B***). Aged (12 months old) BALB/c mice were injected intraperitoneally with 300 μg of antibody, and challenged with a SARS-CoV-2 mouse-adapted (MA) isolate twelve hours later (Dinnon et al., 2020). Mice received either FcγR-dependent, *in vitro* infection-enhancing antibody DH1041, non-infection enhancing antibody DH1050.1, or a combination of both antibodies. Administration of RBD NAb DH1041 alone or in combination with DH1050.1 protected all mice from detectable infectious virus in the lungs 48h after challenge (***Figure 6A***). In the setting of therapeutic treatment, administration of DH1041 alone (300 μg) or in combination with DH1050.1 (150 μg of each) twelve hours after SARS-CoV-2 challenge significantly reduced lung infectious virus titers, with half of the mice having undetectable infectious virus in the lung 48h after challenge (***Figure 6B***). Thus, while RBD antibody DH1041 could mediate FcγR-dependent, *in vitro* infection enhancement, it protected mice from SARS-CoV-2 infection when administered prophylactically or therapeutically.

**Figure 6.**
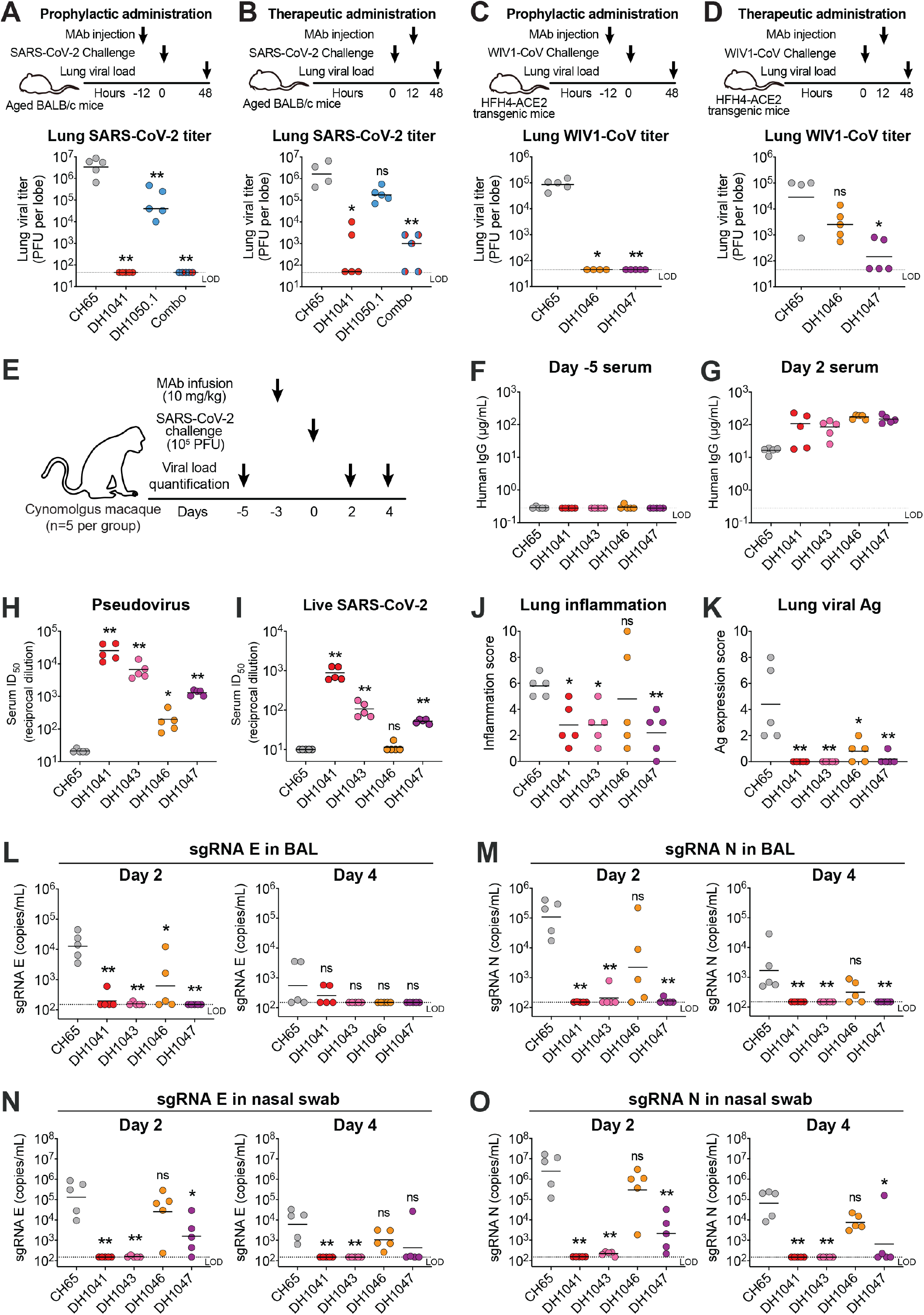
RBD antibodies that mediate FcγR-dependent infection enhancement *in vitro*, protect mice and non-human primates from SARS-CoV-2 challenge. **(A-B)** Protection in BALB/c mice against mouse-adapted SARS-CoV-2. Mice (n=5 per group) were intraperitoneally (i.p.) injected with 300 μg of a single antibody or 150 μg of two antibodies in combination (A) prophylactically at 12h pre-infection or (B) therapeutically at 12h post-infection. Infection was performed with mouse-adapted SARS-CoV-2 2AA MA virus via intranasal (i.n.) route. Titers of infectious virus in the lung were examined 48 post-infection. An irrelevant human antibody CH65 was used as a negative control. **(C-D)** Protection in HFH4-hACE2-transgenic mice against SARS-related bat WIV1-CoV challenge. Mice (n=5 per group) were intraperitoneally (i.p.) injected with 300 μg of indicated antibody or CH65 control antibody (C) prophylactically at 12h pre-infection or (D) therapeutically at 12h post-infection. Infection was performed with WIV1-CoV via i.n. route. Lung viral titers were examined at 48 post-infection. **(E-O)** RBD NAbs and infection-enhancing Abs protected SARS-CoV-2 infection in non-human primates. **(E)** Study design. Cynomolgus macaques (n=5 per group) were infused with DH1041, DH1043, DH1046, DH1047 or an irrelevant CH65 antibody 3 days before 10^5^ PFU of SARS-CoV-2 challenge via intranasal route and intratracheal route. Viral load including viral RNA and subgenomic RNA (sgRNA) were measured on the indicated pre-challenge and post-challenge timepoints. Lungs were harvested on Day 4 post-challenge for histopathology study. **(F-G)** Serum human IgG concentrations at Day -5 (H) and Day 2 (I). **(H-I)** Day 2 serum neutralization titers shown as the reciprocal serum dilution that inhibits 50% (ID_50_) of (**H**) pseudotyped SARS-CoV-2 replication in 293T/ACE2 cells or (**J**) SARS-CoV-2 replication in Vero cells. **(J-K)** Lung histopathology. Sections of the left caudal (Lc), right middle (Rm), and right caudal (Rc) lung were evaluated and scored for (**J**) the presence of inflammation by hematoxylin and eosin (H&E) staining, and (**K**) for the presence of SARS-CoV-2 nucleocapsid by immunohistochemistry (IHC) staining. Symbols indicate the sums of Lc, Rm, and Rc scores for each animal. While two monkeys in the DH1046 RBD antibody infusion group had higher overall lung pathology scores than controls, lung histology did not show alveolar or perivascular edema nor was BAL inflammatory cytokine levels elevated. Thus, these two animals had more lung involved with inflammatory macrophage infiltration but did not have pathological evidence of vascular leakage. **(L-M)** SARS-CoV-2 (**L**) E gene sgRNA and (**M**) N gene sgRNA in bronchoaveolar lavage (BAL) on Day 2 and Day 4 post challenge. **(N-O)** SARS-CoV-2 (**N**) E gene sgRNA and (**O**) N gene sgRNA in nasal swab on Day 2 and Day 4 post challenge. Statistical significance in all the panels were determined using Wilcoxon rank sum exact test. Asterisks show the statistical significance between indicated group and CH65 control group: ns, not significant, *P<0.05, **P<0.01.

DH1046 and DH1047 are RBD cross-reactive antibodies that neutralize SARS-CoV, SARS-CoV-2 and bat WIV1-CoV (***Figures 2A, S4A-B, S5E-H and S31***). Both of these RBD antibodies mediated FcγR-dependent, *in vitro* SARS-CoV-2 infection enhancement of TZM-bl cells that lacked ACE2 expression (***Figures 1F-L***). To determine whether SARS-CoV-2 *in vitro* infection enhancement predicted *in vivo* infection enhancement by SARS-related bat coronaviruses, we assessed the ability of either DH1046 or DH1047 to enhance or protect against bat WIV1-CoV infection in HFH4-ACE2-transgenic mice. Mice were challenged either 12 hours before or 12 hours after intraperitoneal injection of antibody (***Figures 6C-D***). Mice administered DH1046 or DH1047 before challenge had no detectable infectious virus in the lung, whereas control IgG administered mice had a mean titer of 84,896 plaque forming units per lung lobe (***Figure 6C***). Administration of DH1047 after challenge eliminated detectable infectious virus in the lung in 3 of 5 mice (***Figure 6D***). Therapeutic administration of DH1046 reduced infectious virus titers 10-fold compared to negative control IgG (***Figure 6D***). Thus, FcγR-dependent *in vitro* infection-enhancing RBD antibodies DH1046 and DH1047 did not enhance infection *in vivo*, but rather protected mice from SARS-related bat coronavirus infection.

### *In vitro* infection-enhancing RBD antibodies in SARS-CoV-2-challenged nonhuman primates

Finally, we assessed the *in vivo* relevance of RBD antibody infection enhancement in the cynomolgus macaque SARS-CoV-2 intranasal/intratracheal challenge model. We examined *in vivo* infection enhancement by RBD antibodies DH1041, DH1043, DH1046, and DH1047 that neutralized SARS-CoV-2 pseudovirus and replication-competent virus, but enhanced infection *in vitro* in FcγRI or FcγRIIb-expressing TZM-bl cells (***Figures 1 and 6E***). After antibody infusion at 10 mg of antibody per kg of macaque body weight, serum human IgG concentrations reached 11-228 μg/mL at day 2 post-challenge (***Figures 6F-G and S23A-D***). The same macaque serum containing the RBD antibodies exhibited a wide range of neutralization potency (ID_50_ titers) against SARS-CoV-2 pseudovirus or replication-competent virus, commensurate with the neutralization potency of each antibody (***Figures 6H and 6I***). Infusion of RBD antibody DH1041, DH1043, or DH1047 resulted *in vivo* protection from SARS-CoV-2 infection. In macaques administered DH1041, DH1043, or DH1047, lung inflammation was reduced and lung viral antigen was undetectable compared to control (***Figures 6J-K, S24 and S25A***). E gene sgRNA and N gene sgRNA were significantly reduced in the upper and lower respiratory tract based on analyses of bronchoalveolar lavage fluid, nasal swabs, and nasal wash samples (***Figures 6L-O and S23E-I***).

RBD antibody DH1046, a weaker neutralizing Ab compared to DH1041, DH1043 or DH1047 ***(Figure 2A***), did not enhance sgRNA E or N in BAL or nasal swab samples (***Figures 6L-O and S23E-I***), but protected only a subset of infused monkeys. Three monkeys treated with RBD antibody DH1046 exhibited the same or lower levels of lung inflammation compared to monkeys that received control IgG (***Figure 6J***). Two DH1046-infused monkeys had increased lung inflammation scores of 8 and 10 due to increased total areas of inflammation compared to control antibody monkeys (***Figures 6J, S24 and S25***), but had no evidence of perivascular or alveolar edema nor evidence of abnormal BAL cytokines (***Figures S26 and S27***). Thus, these two animals had more lung involved with inflammatory macrophage infiltration but did not have pathological evidence of vascular leakage. Comparing the DH1046 group to the control IgG group, viral nucleocapsid antigen in the lung was reduced (***Figures 6K, S24 and S25A***).

Thus, overall, all 31 spike enhancing antibody-infused monkeys did not show enhanced virus infectivity *in vivo*. In contrast, 3 of 31 antibody-treated monkeys exhibited enhancement of lung pathology, with 1 of 31 antibody-treated monkeys had alveolar and perivascular edema and with elevated BAL inflammatory cytokines.

## DISCUSSION

Here, we demonstrated SARS-CoV-2 RBD and NTD antibodies can increase infection *in vitro*, but either protect or do not increase coronavirus replication in mouse and monkey models *in vivo*. However, increased lung inflammation was observed infrequently in SARS-CoV-2 antibody-treated macaques despite having low or undetectable lung viral antigen. In three of 31 monkeys administered RBD or NTD antibodies that mediated infection enhancement *in vitro*, lung inflammation scores were greater than in control IgG-infused monkeys. One of these three macaques had alveolar edema and elevated BAL cytokine levels compatible with acute respiratory syndrome, while the other two monkeys with high lung inflammation scores did not show edema nor BAL inflammatory cytokines. Thus, only one antibody-infused monkey had severe lung inflammation.

Notably, we observed two different types of *in vitro* infection enhancement. First, RBD antibodies mediated classical antibody-dependent enhancement that required FcγRs and antibody Fc for virus uptake (Lee et al., 2020). Previous studies have demonstrated that uptake of MERS-CoV or SARS-CoV has mostly been mediated by FcγRIIa on the surface of macrophages (Bournazos et al., 2020; Wan et al., 2020; Yip et al., 2016). In contrast to SARS-CoV and MERS-CoV infection enhancing antibodies, we identified SARS-CoV-2 RBD antibodies utilized FcγRIIb or FcγRI. Thus, different FcγRs can mediate SARS-CoV and MERS-CoV *in vitro* infection enhancement compared to SARS-CoV-2 *in vitro* infection enhancement. Second, non-neutralizing NTD antibodies mediated FcγR-independent infection enhancement in two different FcγR-negative, ACE2-expressing cell types. The mechanism of FcγR-negative *in vitro* enhancement remains unclear, but one hypothesis is the possibility of antibody modulation of S protein conformation because of the requirement for ACE2 expression on target cells. In NSEM studies, NTD antibodies preferentially bound to S in a 3-RBD-down conformation. One study has reported that binding of select NTD antibodies to S enhances S binding to ACE2 (Liu et al., 2020b). Whether S protein has different entry kinetics or higher affinity for ACE-2 when liganded to infection-enhancing antibody DH1052 will be a focus of future studies.

Previous studies with vaccine-induced antibodies against SARS-CoV-1 have also shown *in vitro* FcγRII-dependent infection enhancement, but no *in vivo* infection enhancement in hamsters (Kam et al., 2007). Macrophages and other phagocytes are the target cells that take up MERS-CoV leading to infection enhancement (Hui et al., 2020; Wan et al., 2020; Zhou et al., 2014). In contrast, neither SARS-CoV-1 nor SARS-CoV-2 productively infect macrophages (Bournazos et al., 2020; Hui et al., 2020; Yip et al., 2016). Importantly, a recent study demonstrated that alveolar macrophages harboring SARS-CoV-2 RNA produce T cell chemoattractants leading to T cell IFN-γ production that in turn, stimulates inflammatory cytokine release from alveolar macrophages (Grant et al., 2021). Why severe lung pathology and inflammatory cytokine production occurred in only 1 of 31 monkeys is unknown, but may relate to host-specific differences regulating inflammatory cytokine production (Bastard et al., 2020; Zhang et al., 2020). It is important to note that the one monkey that developed alveolar and perivascular edema and elevated BAL inflammatory cytokines could have been caused by antibody enhancement of disease, or could have been due to unknown factors that caused more severe disease in animal BB536A that were unrelated to DH1052 administration. That none of 6 animals infused with a higher dose (30mg/kg) of DH1052 did not have enhanced pulmonary disease supports the hypothesis that the lung pathology may have been a severe case of COVID-19 lung disease unrelated to antibody infusion.

*In vitro* enhancing antibodies may have the ability to suppress SARS-CoV-2 replication *in vivo* through non-neutralizing FcR-mediated antibody effector functions (Bournazos et al., 2020; Schafer et al., 2021). While circulating *in vivo*, antibodies can opsonize infected cells or virions and recruit effector immune cells to kill virus-infected cells through Fc-mediated mechanisms (Bournazos et al., 2020). A recent study in a SARS-CoV-2 mouse model of acquisition suggested that Fc effector functions contribute to the protective activity of SARS-CoV-2 neutralizing antibodies C104, C002, and C110 (Schafer et al., 2021). Thus, antibody effector functions may contribute to the outcome *in vivo*, but not be accounted for in SARS-CoV-2 enhancement or neutralization assays *in vitro*. In support of this hypothesis, infection-enhancing, non-neutralizing NTD antibody DH1052 reduced infectious virus titers in the lungs of challenged mice and monkeys compared to the negative control antibodies. Consistent with previous findings for human IgG (Dekkers et al., 2017), we have demonstrated that DH1052 antibody can bind to select murine FcγRs., raising the hypothesis that DH1052 Fc-mediated effector functions may be responsible for the reduction of virus replication. Future studies will investigate Fc-mediated effector functions of DH1052 to discern their role in reducing infectious SARS-CoV-2 virus titers in mice and monkeys.

Finally, administration of COVID-19 convalescent sera to over 35,000 COVID-19 patients have demonstrated the treatment to be safe and is not associated with enhanced disease (Joyner et al., 2020). Of greater importance is that both the Pfizer/BioNTech and Moderna mRNA-lipid nanoparticle (LNP) vaccine efficacy trials have completed and showed ∼95% vaccine efficacy (Jackson et al., 2020; Polack et al., 2020). That the Moderna mRNA-LNP COVID-19 vaccine efficacy trial had 30 severe cases of COVID-19 occur—all in the placebo group (Baden et al., 2020), demonstrated that if antibody-based enhancement of infection or lung pathology will occur in humans with vaccination, it will be rare. Finally, a recent study demonstrated that suboptimal neutralizing antibody level is a significant predictor of severity for SARS-CoV-2 (Garcia-Beltran et al., 2020). Thus, in spite of the rarity of severe lung pathology associated with presence of anti-spike antibody in animal model studies reported here, it will be important to continue to monitor on-going COVID-19 vaccine efficacy trials for the possibility of vaccine associated enhanced disease when suboptimal neutralizing antibody titers are induced (Haynes et al., 2020).

## METHODS

Detailed methods are provided in the supplemental online material.

## ACKNOWLEDGMENTS

We thank the COVID-19 donors enrolled in the Molecular and Epidemiological Study of Suspected Infection protocol (MESSI), the MESSI clinical support team, the DHVI Clinical Accessioning Unit Core and the DHVI Immunology and Virology Quality Assessment Core for sample procurement, processing and biobanking. We thank Joseph Gilmore, Steve Slater, Kwan-Ki Hwang, Ahmad Yousef Abuahmad, and the Duke Human Vaccine Institute Flow Cytometry Facility for flow cytometric sorting; Tyler Evangelous for purification of PCR product; Caroline Jones for production of fluorophore-labeled reagents; Kara Anasti and the Duke BMI CORE facility for SPR measurements of antibody affinities; Madison Berry and Sravani Venkatayogi for bioinformatics assistance; Paige Rawls, Lena Smith, Jemma Hwang, Beth Bryan, Jingjing Li, Haiyan Chen for protein and DNA production assistance; Nicole De Naeyer for RNA extraction assistance. We thank Drs. Matthew Gagne and Daniel C. Douek at VRC/NIH for the helpful discussions regarding the subgenomic RNA assays. We thank Kelly Soderberg, Elizabeth Donahue, Amelia Karlsson, Amanda Newman and Whitney Edwards for program management. COVID-19 donor sample processing, flow cytometric sorting, and replication-competent virus neutralization assays were performed in the Duke Regional Biocontainment Laboratory, which received partial support for construction from the National Institutes of Health, National Institute of Allergy and Infectious Diseases (UC6-AI058607). This work was supported by a contract from the State of NC funded by the Coronavirus Aid, Relief, and Economic Security Act (CARES Act); an administrative supplement to NIH R01 AI145687 for coronavirus research (P.A.); NIH grants R01AI157155 and U54CA260543 (R.S.B.); NIH NIAID U19AI142596 grant (B.F.H.); NIH R01 AI146779 and a Massachusetts Consortium on Pathogenesis Readiness (MassCPR) grant (A.G.S.); training grants: NIGMS T32 GM007753 (B.M.H. and T.M.C); T32 AI007245 (J.F.); and a cooperative agreement with DOD/DARPA (HR0011-17-2-0069; G.D.S). Cryo-EM data were collected at the National Center for Cryo-EM Access and Training (NCCAT) and the Simons Electron Microscopy Center located at the New York Structural Biology Center, supported by the NIH Common Fund Transformative High Resolution Cryo-Electron Microscopy program (U24 GM129539,) and by grants from the Simons Foundation (SF349247) and NY State. We thank Ed Eng, Carolina Hernandez and Daija Bobe for microscope alignments and assistance with cryo-EM data collection. This study utilized the computational resources offered by Duke Research Computing (http://rc.duke.edu; NIH 1S10OD018164-01) at Duke University. David R. Martinez is funded by an NIH F32 AI152296, a Burroughs Wellcome Fund Postdoctoral Enrichment Program Award, and was previously supported by an NIH NIAID T32 AI007151. We thank M. DeLong, C. Kneifel, M. Newton, V. Orlikowski, T. Milledge, and D. Lane from the Duke Office of Information Technology and Research Computing for assisting with setting up and maintaining the computing environment. We thank Laurent Pessaint, Anthony Cook, Alan Dodson, Katelyn Steingrebe and Bridget Bart at BIOQUAL for administrative and technical assistance with non-human primate studies.

## AUTHOR CONTRIBUTIONS

D.L. designed and performed experiments, analyzed data, and wrote the manuscript. R.J.E., K.Mansouri, K.Manne, S.G., K.J., M.K., V.S. and P.A. performed structure experiments and analysis. R.J.E. and P.A. supervised structural studies and wrote the manuscript. S.M.A. and K.C. performed and analyzed the SPR work. A.S. and D.R.M. performed the mouse challenge work. D.R.M., L.V.T., T.D.S performed SARS-CoV-1 and bat CoV neutralization assays. K.W. performed bioinformatic analysis of antibody sequences. X.L. performed antibody isolation. R.P., V.G., and M.D. performed ELISA assay. L.L.S. oversaw the NHP study. C.T.D. and T.N.D oversaw the viral RNA assay. T.H.O. and G.D.S. performed SARS-CoV-2 virus neutralization assays. E.L., A.F., F.C., G.E.H., A.S., K.T. and C.J. prepared DNA and produced antibodies. L.G.P., C.M. and D.C.M. performed pseudovirus neutralization assays. M.B., T.V.H. and B.F.H. performed autoreactivity assays. W.R. and Y.W. provided statistical analyses. C.W.W and E.P. oversaw setting up the MESSI donor cohort. A.M., E.S., and R.P. performed COVID-19 donor serology. M.A.M. oversaw the protein fluorescent labeling and flow cytometry sorting. D.W.C. designed and performed flow cytometry sorting. A.G.S., J.F., B.M.H, T.M.C., I.T., T.Z., P.D.K., J.M. and B.G. provided key reagents for this study. K.W.B., M.M., B.M.N., I.N.M., and R.S. oversaw or performed pathology experiments. H.A., and M.G.L. led and performed the non-human primate studies. R.S.B. supervised the SARS-CoV-1 and bat CoV neutralization assays and the mouse challenge studies. K.O.S. and B.F.H. conceived, designed and supervised the study, reviewed all data, and wrote the paper. All authors reviewed and approved the manuscript.

