## Supplemental Figures and Methods for "The functions of SARS-CoV-2 neutralizing and infection-enhancing antibodies in vitro and in mice and nonhuman primates"

**This PDF file includes:**

Materials and Methods

References

Supplementary Figs. S1 to S27

Supplementary Tables. S2 to S7

**Other Supplementary Material for this manuscript includes the following:**

Supplementary Table S1

Supplementary Table S8

#### MATERIALS AND METHODS

##### Human Subjects

Nasopharyngeal swabs and peripheral blood samples were collected from a convalescent COVID-19 donor (MESSI ID #450905) on designated days after reported onset of COVID symptoms. The SARS-CoV-1 donor PBMC were provided by NIH/VRC. Human subject specimens were collected and used with the informed consent of study participants and in compliance with the Duke University Medical Center Institutional Review Board (DUHS IRB Pro00100241).

##### Symptom data collections

Participant self-reported symptoms were recorded at each time-point for 39 symptom categories (nasal discharge, nasal congestion, sneezing, coughing, shortness of breath, malaise, throat discomfort, fever, headache, shaking chills, loss of smell, loss of taste, excessive sweating, dizziness, pain behind the eyes, itchy/watery eyes, visual blurring, hearing problems, ear pain, confusion, stiff neck, swollen glands, palpitations, chest pain, pain in joints, muscle soreness, fatigue, loss of appetite, abdominal pain, nausea/vomiting, diarrhea, swelling, itchy skin, rash, skin lesions, unusual bleeding, red fingers or toes, red eyes, other: specify). Each symptom was scored on a scale of 0–4, with 0 indicating not present, 1 mild, 2 moderate, 3 severe, and 4 very severe symptoms. Daily symptom count (number of non-zero symptom categories) and symptom severity (sum of all symptom scores) were determined for each survey timepoint. At enrollment, date of symptom onset was determined, and an initial "historical" symptom survey recorded maximum score for each symptom category between symptom onset and study enrollment.

##### Expression of Recombinant Viral Proteins

The SARS-CoV-2 ectodomain constructs were produce and purified as describe previously (Wrapp, D.et al. 2020). Plasmids encoding Spike-2P and HexaPro (Hsieh et al., 2020) were transiently transfected in FreeStyle 293 cells (Thermo Fisher) using Turbo293 (SpeedBiosystems). The cultures were collected on Day 6 post transfection. The cells were separated from the medium by centrifugation. Protein were purified from filtered cell supernatants by StrepTactin resin (IBA) and additionally by size exclusive chromatography using Superose 6 10/300 increase column (GE Healthcare) in 2mM Tris pH 8, 200mMnNaCl, 0.02% $NaN_3$. SARS-CoV-2 NTD was produced as previously described (Zhou et al., 2020). SARS-CoV-1 RBD and MERS-CoV Spike RBD were cloned into pVRC vector for mammalian expression (FreeStyle 293F or Expi293F suspension cells). The construct contains an HRV 3C-cleavable C-terminal SBP-8xHis tag. Supernatants were harvested 5 days post-transfection and passaged directly over Cobalt-TALON resin (Takara) followed by size exclusion chromatography on Superdex 200 Increase (GE Healthcare) in 1x PBS. Typical yields from FreeStyle 293F cells are approximately 50 mg/liter culture. Affinity tags can be removed using HRV 3C protease (ThermoScientific) and the protein repurified using Cobalt-TALON resin to remove the protease, tag and non-cleaved protein.

**Antigen-Specific Single B Cell Sorting**

Plasmablasts were sorted by flow cytometry from the SARS-CoV-2 donor on Day 11 and Day 15 post symptom onset. PBMCs were stained with optimal concentrations of the following fluorochrome-antibody conjugates: IgD PE (Clone# IA6-2, BD Biosciences, Catalog# 555779), CD3 PE-Cy5 (Clone# HIT3a, BD Biosciences, Catalog# 555341), CD10 PE-CF594 (Clone# HI10A, BD Biosciences, Catalog# 562396), CD27 PE-Cy7 (Clone# O323, eBioscience, Catalog# 25-0279), CD38 APC-Alexa Fluor (AF) 700 (Clone# LS198-4-2, Beckman Coulter, Catalog# B23489), CD19 APC-Cy7 (Clone# LJ25C1, BD Biosciences, Catalog# 561743), CD16 BV570 (Clone# 3G8, Biolegend, Catalog# 302035), CD14 BV605 (Clone# M5E2, Biolegend, Catalog# 301834), and CD20 BV650 (Clone# 2H7, BD, Catalog# 563780). The cells were then labeled with Fixable Aqua Live/Dead Cell Stain Kit (Invitrogen, Catalog# L34957). On a BD FACSAria II flow cytometer (BD Biosciences), plasmablasts were identified as viable $CD14^{-}$/$CD16^{-}$/$CD19^{+}$/$CD20^{low}$/$IgD^{-}$/$CD27^{high}$/$CD38^{high}$ cells and sorted as single cells into 96-well plates containing lysis buffer. Sorted plates were frozen at -80°C in the DHVI Flow Facility under BSL3 precautions in the Duke Regional Biocontainment Laboratory (Durham, NC) until processing.

Antigen-specific memory B cells (MBCs) were isolated by flow cytometric sorting from the SARS-CoV-2 donor on Day 36 post symptom onset, and a donor with SARS-CoV-1 history. PBMCs were stained with IgD FITC (Clone# IA6-2, BD Biosciences, Catalog# 555778), IgM PerCp-Cy5.5 (Clone# G20-127, BD Biosciences, Catalog# 561285), CD10 PE-CF594 (Clone# HI10A, BD Biosciences, Catalog# 562396), CD3 PE-Cy5 (Clone# HIT3a, BD Biosciences, Catalog# 555341), CD235a PE-Cy5 (Clone# GA-R2, BD Biosciences, Catalog# 559944), CD27 PE-Cy7 (Clone# O323, eBioscience, Catalog# 25-0279), CD38 APC-AF700 (Clone# LS198-4-2, Beckman Coulter, Catalog# B23489), CD19 APC-Cy7 (Clone# LJ25C1, BD Biosciences, Catalog# 561743), CD14 BV605 (Clone# M5E2, Biolegend, Catalog# 301834), CD16 BV570 (Clone# 3G8, Biolegend, Catalog# 302035), and fluorescent-labeled SARS-CoV-2 Spike probes (AF647-conjugated Spike-2P, PE-conjugated Spike-2P, AF647-conjugated NTD, AF647-conjugated RBD, VioBright 515-conjugated RBD). The cells were then labeled with Fixable Aqua Live/Dead Cell Stain Kit (Invitrogen, Catalog# L34957). On a BD FACSAria II flow cytometer (BD Biosciences), antigen-specific MBCs were identified as viable $CD3^-$/$CD14^-$/$CD16^-$/$CD235a^-$/$CD19^+$/$IgD^-$/$probe^+$ cells and were sorted as single cells into 96-well plates containing lysis buffer. Collection plates were immediately frozen in a dry ice/ethanol bath, and stored at -80 °C in the DHVI Flow Facility under BSL3 precautions in the Duke Regional Biocontiamet Laboratory until processing. Flow cytometric data were analyzed using FlowJo version 10.

**PCR Amplification of Human Antibody Genes**

Antibody genes were amplified by RT-PCR from flow cytometry-sorted single B cells using the methods as described previously (Liao et al., 2009; Wrammert et al., 2008) with modification. The PCR-amplified genes were then purified and sequenced with 10 μM forward and reverse primers. Sequences were analyzed by using the human library in Clonalyst for the VDJ arrangements of the immunoglobulin IGHV, IGKV, and IGLV sequences and mutation frequencies (Kepler et al., 2014). Clonal relatedness of $V_HD_HJ_H$ and $V_LJ_L$ sequences was determined as previously described (Liao et al., 2013).

**Expression of Antibody Viable Region Genes as Full-Length IgG Recombinant mAbs**

Transient transfection of recombinant antibodies was performed as previously described (Liao et al., 2009). Briefly, purified PCR products were used for overlapping PCR to generate linear human IgG expression cassettes. The expression cassettes were transfected into 293i cells using ExpiFectamine (Thermo Fisher Scientific, Catalog# A14525). The supernatant samples containing recombinant IgGs were used for IgG quantification and preliminary ELISA binding screening.

The down-selected human antibody genes were then synthesized and cloned (GenScript) in a human IgG1 backbone with 4A mutations to enhance antibody-dependent cell-mediated cytotoxicity (ADCC) or a human IgG1 backbone with a LS mutation to extent antibody half-life (Saunders, 2019). Recombinant IgG antibodies were then produced in HEK293i suspension cells by transfection with ExpiFectamine and purified using Protein A resin. The purified IgG antibodies were run in SDS-PAGE for Coomassie blue staining and western blot for quality control and then used for the downstream experiments.

**Antibody Binding ELISA**

For ELISA binding assays of Coronavirus Spike antibodies, the antigen panel included SARS-CoV-2 Spike S1+S2 ectodomain (ECD) (SINO, Catalog # 40589-V08B1), SARS-CoV-2 Spike-2P (Wrapp et al., 2020), SARS-CoV-2 Spike S2 ECD (SINO, Catalog # 40590-V08B), SARS-CoV-2 Spike RBD from insect cell sf9 (SINO, Catalog # 40592-V08B), SARS-CoV-2 Spike RBD from mammalian cell 293 (SINO, Catalog # 40592- V08H), SARS-CoV-2 Spike NTD-Biotin, SARS-CoV Spike Protein DeltaTM (BEI, Catalog # NR-722), SARS-CoV WH20 Spike RBD (SINO, Catalog # 40150-V08B2), SARS-CoV WH20 Spike S1 (SINO, Catalog #40150-V08B1), SARS-CoV-1 RBD, MERS-CoV Spike S1+S2 (SINO, Catalog # 40069-V08B), MERS-CoV Spike S1 (SINO, Catalog #40069-V08B1), MERS-CoV Spike S2 (SINO, Catalog #40070-V08B), MERS-CoV Spike RBD (SINO, Catalog #40071-V08B1), MERS-CoV Spike RBD. In preliminary ELISA screening of the transient transfection supernatants, we also screened the antibodies against SARS-CoV CL Protease protein (BEI, Catalog # 30105) and SARS-CoV Membrane (M) protein (BEI, Catalog # 110705).

For binding ELISA, 384-well ELISA plates were coated with 2 μg/mL of antigens in 0.1 M sodium bicarbonate overnight at 4°C. Plates were washed with PBS + 0.05% Tween 20 and blocked with blocked with assay diluent (PBS containing 4% (w/v) whey protein, 15% Normal Goat Serum, 0.5% Tween-20, and 0.05% Sodium Azide) at room temperature for 1 hour. Purified MAb samples in 3-fold serial dilutions in assay diluent starting at 100 μg/mL, or un-diluted transfection supernatant were added and incubated for 1 hour, followed by washing with PBS-0.1% Tween 20. HRP-conjugated goat anti-human IgG secondary Ab (SouthernBiotech, catalog# 2040-05) was diluted to 1:10,000 and incubated at room temperature for 1 hour. These plates were washed four times and developed with tetramethylbenzidine substrate (SureBlue Reserve- KPL). The reaction was stopped with 1 M HCl, and optical density at 450 nm ($OD_{450}$) was determined.

**Affinity Measurements**

SPR measurements of SARS-CoV-2 antibody Fab binding to Spike-2P or Spike-Hexaro proteins were performed using a Biacore S200 instrument (Cytiva, formerly GE Healthcare, DHVI BIA Core Facility, Durham, NC) in HBS-EP+ 1x running buffer. The Spike proteins were first captured onto a Series S Streptavidin chip to a level of 300-400 RU for Spike-2P and 350-450 resonance units (RU) for Spike-HexaPro. The antibody Fabs were injected at 0.5 to 500 nM over the captured S proteins using the single cycle kinetics injection mode at a flow rate of 50uL/min. Association phase was maintained with either 120 or 240 second injections of each Fab at increasing concentrations followed by a dissociation of 600 seconds after the final injection. After dissociation, the S proteins were regenerated from the streptavidin surface using a 30 second pulse of Glycine pH1.5. Results were analyzed using the Biacore S200 Evaluation software (Cytiva). A blank streptavidin surface along with blank buffer binding were used for double reference subtraction to account for non-specific protein binding and signal drift. Subsequent curve fitting analyses were performed using a 1:1 Langmuir model with a local Rmax for the

Fabs with the exception of DH1050.1 Fab which was fit using the heterogeneous ligand model with local Rmax. The reported binding curves are representative of two data sets.

**Surface Plasmon Resonance Antibody Blocking Assay**

RBD and NTD Abs binding to S protein was measured by surface plasmon resonance (BIAcore 3000; Cytiva, formerly GE Healthcare, DHVI BIA Core Facility, Durham, NC) analysis. Antibody binding competition and blocking were measured by SPR following immobilization by amine coupling of monoclonal antibodies to CM5 sensor chips (BIAcore/Cytiva). Antibody competition experiments were performed by mixing S protein and mAb (30 minutes incubation) followed by injection for 5 minutes at 50 μL/min. In separate assays and from analysis of binding to an identical epitope binding ligand, it was determined that S protein at 20 μm and antibody at 200 μm bind to complete saturation. Antibody blocking assays were performed by co-injecting S protein (20 μM) over mAb immobilized surfaces for 3 minutes at 30 μL/min and a test Ab (200 μM) for 3 minutes at 30 μL/min. The dissociation of the antibody sandwich complex with the spike protein was monitored for 10 minutes with buffer flow and then a 24 second injection of Glycine pH2.0 for regeneration. Blank buffer binding was used for subtraction to account for signal drift. Data analyses were performed with BIA-evaluation 4.1 software (BIAcore/Cytiva).

**ACE2-blocking assay**

For ACE-2 blocking assays, plates were coated as stated above with 2 ug/mL recombinant ACE-2 protein, then washed and blocked with 3% BSA in 1X PBS. While assay plates blocked, purified antibodies were diluted as stated above, only in 1% BSA with 0.05% Tween-20. In a separate dilution plate Spike-2P protein was mixed with the antibodies at a final concentration equal to the EC50 at which spike binds to ACE-2 protein. The mixture was allowed to incubate at room temperature for 1 hour. Blocked assay plates were then washed and the antibody-spike mixture was added to the assay plates for a period of 1 hour at room temperature. Plates were washed and a polyclonal rabbit serum against the same spike protein (nCoV-1 nCoV-2P.293F) was added for 1 hour, washed and detected with goat anti rabbit-HRP (Abcam cat# ab97080) followed by TMB substrate. The extent to which antibodies were able to block the binding spike protein to ACE-2 was determined by comparing the OD of antibody samples at 450 nm to the OD of samples containing spike protein only with no antibody. The following formula was used to calculate percent blocking: blocking% = (100 - (OD sample/OD of spike only)*100).

**Negative-stain electron microscopy**

For each Fab-spike complex, an aliquot of spike protein at ~1-5 mg/ml concentration that had been flash frozen and stored at -80 °C was thawed in an aluminum block at 37 °C for 5 minutes; then 1-4 µl of spike was mixed with sufficient Fab to give a 9:1 molar ratio of Fab to spike and incubated for 1 hour at 37 °C. The complex was then cross-linked by diluting to a final spike concentration of 0.1 mg/ml into room-temperature buffer containing 150 mM NaCl, 20 mM HEPES pH 7.4, 5% glycerol, and 7.5 mM glutaraldehyde. After 5 minutes cross-linking, excess glutaraldehyde was quenched by adding sufficient 1 M Tris pH 7.4 stock to give a final concentration of 75 mM Tris and incubated for 5 minutes. For negative stain, carbon-coated grids (EMS, CF300-cu-UL) were glow-discharged for 20s at 15 mA, after which a 5-µl drop of quenched sample was incubated on the grid for 10-15 s, blotted, and then stained with 2% uranyl formate. After air drying grids were imaged with a Philips EM420 electron microscope operated at 120 kV, at 82,000x magnification and images captured with a 2k x 2k CCD camera at a pixel size of 4.02 Å.

**Image processing of negative stain images**

The RELION 3.0 program was used for all negative stain image processing. Images were imported, CTF-corrected with CTFFIND, and particles were picked using a spike template from previous 2D class averages of spike alone. Extracted particle stacks were subjected to 2-3 rounds of 2D class averaging and selection to discard junk particles and background picks. Cleaned particle stacks were then subjected to 3D classification using a starting model created from a bare spike model, PDB 6vsb, low-pass filtered to

30 Å. Classes that showed clearly-defined Fabs were selected for final refinements followed by automatic filtering and B-factor sharpening with the default Relion post-processing parameters.

**Cryo-EM sample preparation, data collection and processing**

To prepare antibody-bound complexes of the SARS-CoV-2 2P spike, the spike at a final concentration of 1–2 mg/mL, in a buffer containing 2 mM Tris pH 8.0, 200 mM NaCl and 0.02% NaN3, was incubated with 5-6 fold molar excess of the antibody Fab fragments for 30–60 min. 2.5 µL of protein was deposited on a Quantifoil-1.2/1.3 holey carbon grid that had been glow discharged for 15s in a PELCO easiGlow™ Glow Discharge Cleaning System. After a 30 s incubation in >95% humidity, excess protein was blotted away for 2.5 seconds before being plunge frozen into liquid ethane using a Leica EM GP2 plunge freezer (Leica Microsystems). Cryo-EM data were collected on a Titan Krios (Thermo Fisher) equipped with a K3 detector (Gatan). Data were acquired using the Leginon system (Suloway et al., 2005). All the datasets were energy filtered through a either a 20eV or 30eV slit. The dose was fractionated over 50 raw frames and collected at 50ms framerate. Individual frames were aligned and dose-weighted (Zheng et al., 2017). CTF estimation, particle picking, 2D classifications, *ab initio* model generation, heterogeneous refinements, homogeneous 3D refinements and local resolution calculations were carried out in cryoSPARC (Punjani et al., 2017).

**Cryo-EM structure fitting and analysis**

Previously published SARS-CoV-2 ectodomain structures of the all 'down' state (PDB ID 6VXX) and single RBD 'up' state (PDB ID 6VYB), and models of 2-RBD-up and 3-RBD-up states derived from these, were used to fit the cryo-EM maps in Chimera (Pettersen et al., 2004a). Models of Fabs were generated in SWIS-MODEL and docked into the cryo-EM reconstructions using Chimera. Mutations were made in Coot (Emsley and Cowtan, 2004). Coordinates were fit to the maps first using ISOLDE (Croll, 2018) followed by iterative refinement using Phenix (Afonine et al., 2018) real space refinement and subsequent manual coordinate fitting in Coot as needed. Structure and map analysis were performed using PyMol (Schrodinger, 2015), Chimera (Pettersen et al., 2004b) and ChimeraX (Goddard et al., 2018).

**Live SARS-CoV-2 neutralization assays**

The SARS-CoV-2 virus (Isolate USA-WA1/2020, NR-52281) was deposited by the Centers for Disease Control and Prevention and obtained through BEI Resources, NIAID, NIH. SARS-CoV-2 Micro-neutralization (MN) assays were adapted from a previous study (Berry et al., 2004). In short, sera or purified antibodies are diluted two-fold and incubated with 100 TCID50 virus for 1 hour. These dilutions are used as the input material for a TCID50. Each batch of MN includes a known neutralizing control antibody (Clone D001; SINO, CAT# 40150-D001). Data are reported as the last concentration at which a test sample protects Vero E6 cells.

SARS-CoV-2 Plaque Reduction Neutralization Test (PRNT) were performed in the Duke Regional Biocontaiment Laboratory BSL3 (Durham, NC) as previously described with virus-specific modifications (Berry et al., 2004). Briefly, two-fold dilutions of a test sample (e.g. serum, plasma, purified Ab) were incubated with 50 PFU SARS-CoV-2 virus (Isolate USA-WA1/2020, NR-52281) for 1 hour. The antibody/virus mixture is used to inoculate Vero E6 cells in a standard plaque assay (Coleman and Frieman, 2015; Kint et al., 2015). Briefly, infected cultures are incubated at 37°C, 5% CO2 for 1 hour. At the end of the incubation, 1 mL of a viscous overlay (1:1 2X DMEM and 1.2% methylcellulose) is added to each well. Plates are incubated for 4 days. After fixation, staining and washing, plates are dried and plaques from each dilution of each sample are counted. Data are reported as the concentration at which 50% of input virus is neutralized.  A known neutralizing control antibody is included in each batch run (Clone D001; SINO, CAT# 40150-D001). GraphPad Prism was used to determine $IC/EC_{50}$ values.

SARS-CoV-2 nano-luciferase (nanoLuc), SARS-CoV-1 nanoLuc and WIV1-CoV nanoLuc replication-competent virus neutralization assay were described previously (Hou et al., 2020; Menachery et al., 2016; Sheahan et al., 2017).

**Pseudo-typed SARS-CoV-2 neutralization assay and infection-enhancing assays**

Neutralization of SARS-CoV-2 Spike-pseudotyped virus was performed by adopting an infection assay described previously (Korber et al., 2020) with lentiviral vectors and infection in either 293T/ACE2.MF (the cell line was kindly provided by Drs. Mike Farzan and Huihui Mu at Scripps). Cells were maintained in DMEM containing 10% FBS and 50 µg/ml gentamicin. An expression plasmid encoding codon-optimized full-length spike of the Wuhan-1 strain (VRC7480), was provided by Drs. Barney Graham and Kizzmekia Corbett at the Vaccine Research Center, National Institutes of Health (USA). The D614G mutation was introduced into VRC7480 by site-directed mutagenesis using the QuikChange Lightning Site-Directed Mutagenesis Kit from Agilent Technologies (Catalog # 210518). The mutation was confirmed by full-length spike gene sequencing. Pseudovirions were produced in HEK 293T/17 cells (ATCC cat. no. CRL-11268) by transfection using Fugene 6 (Promega, Catalog #E2692). Pseudovirions for 293T/ACE2 infection were produced by co-transfection with a lentiviral backbone (pCMV ΔR8.2) and firefly luciferase reporter gene (pHR' CMV Luc) (Naldini et al., 1996). Culture supernatants from transfections were clarified of cells by low-speed centrifugation and filtration (0.45 µm filter) and stored in 1 ml aliquots at -80 °C.

For 293T/ACE2 neutralization assays, a pre-titrated dose of virus was incubated with 8 serial 3-fold or 5-fold dilutions of mAbs in duplicate in a total volume of 150 µl for 1 hr at 37 °C in 96-well flat-bottom poly-L-lysine-coated culture plates (Corning Biocoat). Cells were suspended using TrypLE express enzyme solution (Thermo Fisher Scientific) and immediately added to all wells (10,000 cells in 100 µL of growth medium per well). One set of 8 control wells received cells + virus (virus control) and another set of 8 wells received cells only (background control). After 66-72 hrs of incubation, medium was removed by gentle aspiration and 30 µL of Promega 1x lysis buffer was added to all wells. After a 10-minute incubation at room temperature, 100 µl of Bright-Glo luciferase reagent was added to all wells. After 1-2 minutes, 110 µl of the cell lysate was transferred to a black/white plate (Perkin-Elmer). Luminescence was measured using a PerkinElmer Life Sciences, Model Victor2 luminometer.

Neutralization titers are the mAb concentration (IC50/IC80) at which relative luminescence units (RLU) were reduced by 50% and 80% compared to virus control wells after subtraction of background RLUs. Negative neutralization values are indicative of infection-enhancement. Maximum percent inhibition (MPI) is the reduction in RLU at the highest mAb concentration tested.

For the TZM-bl neutralization assays, a pre-titrated dose of virus was incubated with serial 3-fold dilutions of test sample in duplicate in a total volume of 150 ul for 1 hr at 37 °C in 96-well flat-bottom culture plates. Freshly trypsinized cells (10,000 cells in 100 μl of growth medium containing 75 μg/ml DEAE dextran) were added to each well. One set of control wells received cells + virus (virus control) and another set received cells only (background control). After 68-72 hours of incubation, 150 μl of cultured medium was removed from each well, and 100ul of Britelite Luminescence Reporter Gene Assay System (PerkinElmer Life Sciences) were added and plates incubated for 2 min at room temperature. After this period 150 μl of the lysate was transferred to black solid plates (Costar) for measurements of luminescence in a Perking Elmer instrument. Neutralization titers are the serum dilution at which relative luminescence units (RLU) were reduced by 50% and 80% compared to virus control wells after subtraction of background RLUs. MPI is the reduction in RLU at the highest mAb concentration tested. Infection-enhancing assays were performed with the same format but using TZM-bl cell lines stably expressing each of the four human FcγR receptors (Perez et al., 2009). In this assay an increase in RLUs over the virus control signal represents FcR-mediated entry.

**Non-human primate protection study**

Groups of five cynomolgus macaques (4-8 kg) were given intravenous infusion with antibodies at 10 mg/kg body weight on Day -3, relative to infectious virus challenge. For each animal, $10^5$ PFU (~$10^6$ TCID50) SARS-CoV-2 virus (Isolate USA-WA1/2020) were diluted in 4 mL, and were given by 1 mL intranasally and 3 mL intratracheally on Day 0. Plasma and serum samples were collected on Day -5, 0, 2, and 4. Nasal swabs, nasal washes, and bronchoalveolar lavage (BAL) were collected on Day -5, 2, and 4.

**Histopathology**

Lung specimen from nonhuman primates were fixed in 10% neutral buffered formalin, processed, and blocked in paraffin for histological analysis. All samples were sectioned at 5 µm and stained with hematoxylin-eosin (H&E) for routine histopathology. Samples were evaluated by a board-certified veterinary pathologist in a blinded manner. Sections were examined under light microscopy using an Olympus BX51 microscope and photographs were taken using an Olympus DP73 camera.

**Immunohistochemistry (IHC)**

Staining for SARS-CoV-2 antigen was achieved on the Bond RX automated system with the Polymer Define Detection System (Leica) used per manufacturer's protocol. Tissue sections were dewaxed with Bond Dewaxing Solution (Leica) at 72oC for 30 min then subsequently rehydrated with graded alcohol washes and 1x Immuno Wash (StatLab). Heat-induced epitope retrieval (HIER) was performed using Epitope Retrieval Solution 1 (Leica), heated to 100oC for 20 minutes. A peroxide block (Leica) was applied for 5 min to quench endogenous peroxidase activity prior to applying the SARS-CoV-2 antibody (1:2000, GeneTex, GTX135357). Antibodies were diluted in Background Reducing Antibody Diluent (Agilent). The tissue was subsequently incubated with an anti-rabbit HRP polymer (Leica) and colorized with 3,3'-Diaminobenzidine (DAB) chromogen for 10 min. Slides were counterstained with hematoxylin.

**Luminex assay**

For cytokine profiling, 7-fold concentrated cynomolgus macaques BAL samples were measured using a 25-analyte multiplex bead array (Millipore, catalog # PRCYT2MAG40K) including sCD137, Eotaxin, sFasL, FGF-2, Fractalkine, Granzyme A, Granzyme B, IL-1α, IL-2, IL-4, IL-6, IL-16, IL-17A, IL-17E/IL-25, IL-21, IL-22, IL-23, IL-28A, IL-31, IL-33, IP-10, MIP-3α, Perforin, RANTES, TNFβ. Samples were prepared according to the manufacturer's recommended protocol and read using a Flexmap

3D suspension array reader (Luminex Corp.). Data were analyzed using Bio-Plex manager software v6.2 (Bio-Rad).

For human antibody quantification, SARS-CoV-2 Spike-2P protein, A/Solomon Islands/3/2006 hemagglutinin (HA) protein or bovine serum albumen (Sigma) was carbodiimide coupled to MagPlex-C beads (Luminex Corp) according to the bead manufacturer's protocol. Briefly, beads were washed in $H_2O$ then activated by incubation with 5 mg/mL sulfo-N-hydroxysulfosuccinimide and 5 mg/mL 1-ethyl-3-(3-dimethylaminopropyl) carbodiimide hydrochloride (ThermoFisher) for 20 minutes. Activated beads were washed twice in PBS (ThermoFisher) and then vortexed at 1,500 RPM for two hours at room temperature with 25 µg protein per 5.0 x $10^6$ beads. Labelled beads were washed in PBS (ThermoFisher), 1% BSA, 0.02% Tween-20, 0.05% Sodium Azide (all Sigma), counted using a hemacytometer and stored at -80°C. NHP sera were diluted 1:200 in assay buffer (PBS, 1% BSA, pH 7.4, Gibco), then 50 µL of diluted sera or monoclonal antibody 3-fold serially diluted in assay buffer (1000-0.45ng/mL) was added to a 96-well plate and mixed with 50 µL assay buffer containing 2500 BSA-conjugated beads (negative control) plus 2500 HA or Spike-conjugated beads. The plate was shaken at 800RPM for 60 minutes at room temperature, washed twice in assay buffer and then 100 µL 4 µg/mL biotin-conjugated mouse anti-human IgG Fc clone H2 (Southern Biotech) in assay buffer was added to every well. The plate was shaken at 800 RPM for 30 minutes at room temperature, washed two times in assay buffer and then 50 µL 4 µg/mL streptavidin-r-phycoerythrin (Invitrogen) in assay buffer was added to every well. The plate was shaken at 800 RPM for 30 minutes at room temperature and washed twice in assay buffer. Beads were resuspended in 150 µL/well assay buffer, shaken at 800 RPM for 15 minutes at room temperature and then analyzed on a BioPlex 200 bead reader (Bio-Rad). Sera antigen-specific antibody concentrations were calculated using Bio-Plex Manager software (Bio-Rad) by extrapolating from the results of the serially-diluted monoclonal antibody. Sera with antibodies above the upper limit of quantitation were re-assayed at 1:1000 or 1:5000. The limit of detection (LOD) for this assay is 0.278 µg/mL.

**Mouse protection study**

Eleven to twelve-month old female immunocompetent BALB/c mice purchased from Envigo (BALB/c AnNHsd, stock# 047) were used for SARS-CoV-2 *in vivo* protection experiments as described previously (Dinnon et al., 2020). Ten-week-old *HFH4-hACE2* transgenic mice were bred and maintained at the University of North Carolina at Chapel Hill and used for WIV-1 *in vivo* protection experiments. Mice were housed in groups of five animals per cage and fed standard chow diet. Virus inoculations were performed under anesthesia (Ketamine and Xylazine) and effort was taken to minimize animal suffering. For evaluating the prophylactic efficacy of mAbs, mice were intraperitoneally treated with 300 μg of each mAb or 150 μg of each mAb in combination 12 hours prior to infection. Mice were infected intranasally with $1X10^5$ PFU of mouse-adapted SARS-CoV-2 2AA MA (Dinnon et al., 2020) or WIV-1. For evaluating the therapeutic efficacy of mAbs, mice were intraperitoneally treated with 300 μg of each mAb or 150 μg of each mAb in combination 12 hours following infection. Forty-eight hours post infection, mice were sacrificed, and lungs were harvested for viral titer as measured by plaque assays and RNA analysis. In another study, fifty-two weeks old female BALB/c mice were i.p. injected with DH1052 (200ug/mice, n=10) or CH65 control antibody (200ug/mice, n=9). After 12 hours, mice were challenged with 1X10^4 PFU of mouse-adapted SARS-CoV-2 MA10 virus (Leist et al., 2020). Mice were sacrificed at day 4 post infection, and lungs were harvested for viral titer as measured by plaque assays and RNA analysis. The study was carried out in accordance with the recommendations for care and use of animals by the Office of Laboratory Animal Welfare (OLAW), National Institutes of Health and the Institutional Animal Care. All mouse studies were performed at the University of North Carolina (Animal Welfare Assurance #A3410-01) using protocols (19-168) approved by the UNC Institutional Animal Care and Use Committee (IACUC) and all mouse studies were performed in a BSL3 facility at UNC.

**Viral RNA Extraction and Quantification**

The assay for SARS-CoV-2 quantitative Polymerase Chain Reaction (qPCR) detects total RNA using the WHO primer/probe set E_Sarbeco (Charité/Berlin). A QIAsymphony SP (Qiagen, Hilden, Germany) automated sample preparation platform along with a virus/pathogen DSP midi kit and the *complex800* protocol were used to extract viral RNA from 800 µL of pooled samples. A reverse primer specific to the envelope gene of SARS-CoV-2 (5'-ATA TTG CAG CAG TAC GCA CAC A-3') was annealed to the extracted RNA and then reverse transcribed into cDNA using SuperScript™ III Reverse Transcriptase (Thermo Fisher Scientific, Waltham, MA ) along with RNAse Out (Thermo Fisher Scientific, Waltham, MA ). The resulting cDNA was treated with RNase H (Thermo Fisher Scientific, Waltham, MA ) and then added to a custom 4x TaqMan™ Gene Expression Master Mix (Thermo Fisher Scientific, Waltham, MA ) containing primers and a fluorescently labeled hydrolysis probe specific for the envelope gene of SARS-CoV-2 (forward primer 5'-ACA GGT ACG TTA ATA GTT AAT AGC GT-3', reverse primer 5'-ATA TTG CAG CAG TAC GCA CAC A-3', probe 5'-6FAM/AC ACT AGC C/ZEN/A TCC TTA CTG CGC TTC G/IABkFQ-3'). The qPCR was carried out on a QuantStudio 3 Real-Time PCR System (Thermo Fisher Scientific, Waltham, MA) using the following thermal cycler parameters: heat to 50°C, hold for 2 min, heat to 95°C, hold for 10 min, then the following parameters are repeated for 50 cycles: heat to 95°C, hold for 15 seconds, cool to 60°C and hold for 1 minute. SARS-CoV-2 RNA copies per reaction were interpolated using quantification cycle data and a serial dilution of a highly characterized custom DNA plasmid containing the SARS-CoV-2 envelope gene sequence. Mean RNA copies per milliliter were then calculated by applying the assay dilution factor (DF=11.7). The limit of detection (LOD) for this assay is approximately 62 RNA copies per milliliter of sample.

**Subgenomic mRNA assay**

SARS-CoV-2 E gene and N gene subgenomic mRNA (sgmRNA) was measured by a one-step RT-qPCR adapted from previously described methods (Wolfel et al., 2020; Yu et al., 2020). To generate standard curves, a SARS-CoV-2 E gene sgmRNA sequence, including the 5'UTR leader sequence, transcriptional regulatory sequence (TRS), and the first 228 bp of E gene, was cloned into a pcDNA3.1

plasmid. For generating SARS-CoV-2 N gene sgmRNA, the E gene was replaced with the first 227 bp of N gene. The recombinant pcDNA3.1 plasmids were linearized, transcribed using MEGAscript T7 Transcription Kit (ThermoFisher, Catalog # AM1334), and purified with MEGAclear Transcription Clean-Up Kit (ThermoFisher, Catalog # AM1908). The purified RNA products were quantified on Nanodrop, serial diluted, and aliquoted as E sgmRNA or N sgmRNA standards.

RNA extracted from animal samples or standards were then measured in Taqman custom gene expression assays (ThermoFisher Scientific). For these assays we used TaqMan Fast Virus 1-Step Master Mix (ThermoFisher, catalog # 4444432) and custom primers/probes targeting the E gene sgmRNA (forward primer: 5’ CGATCTCTTGTAGATCTGTTCTCE 3’; reverse primer: 5’ ATATTGCAGCAGT ACGCACACA 3’; probe: 5’ FAM-ACACTAGCCATCCTTACTGCGCTTCG-BHQ1 3’) or the N gene sgmRNA (forward primer: 5’ CGATCTCTTGTAGATCTGTTCTC 3’; reverse primer: 5’ GGTGAA CCAAGACGCAGTAT 3’; probe: 5’ FAM-TAACCAGAATGGAGAACGCAGTG GG-BHQ1 3’). RT-Qpcr reactions were carried out on a QuantStudio 3 Real-Time PCR System (Applied Biosystems) or a StepOnePlus Real-Time PCR System (Applied Biosystems) using a program below: reverse transcription at 50°C for 5 minutes, initial denaturation at 95°C for 20 seconds, then 40 cycles of denaturation-annealing-extension at 95°C for 15 seconds and 60°C for 30 seconds. Standard curves were used to calculate E or N sgmRNA in copies per ml; the limit of detections (LOD) for both E and N sgRNA assays were 12.5 copies per reaction or 150 copies per mL of BAL/nasal swab/nasal wash.

**Statistics Analysis**

Data were plotted using Prism GraphPad 8.0. Wilcoxon rank sum exact test was performed to compare differences between groups with p-value < 0.05 considered significant using SAS 9.4 (SAS Institute, Cary, NC).

### Figure S1

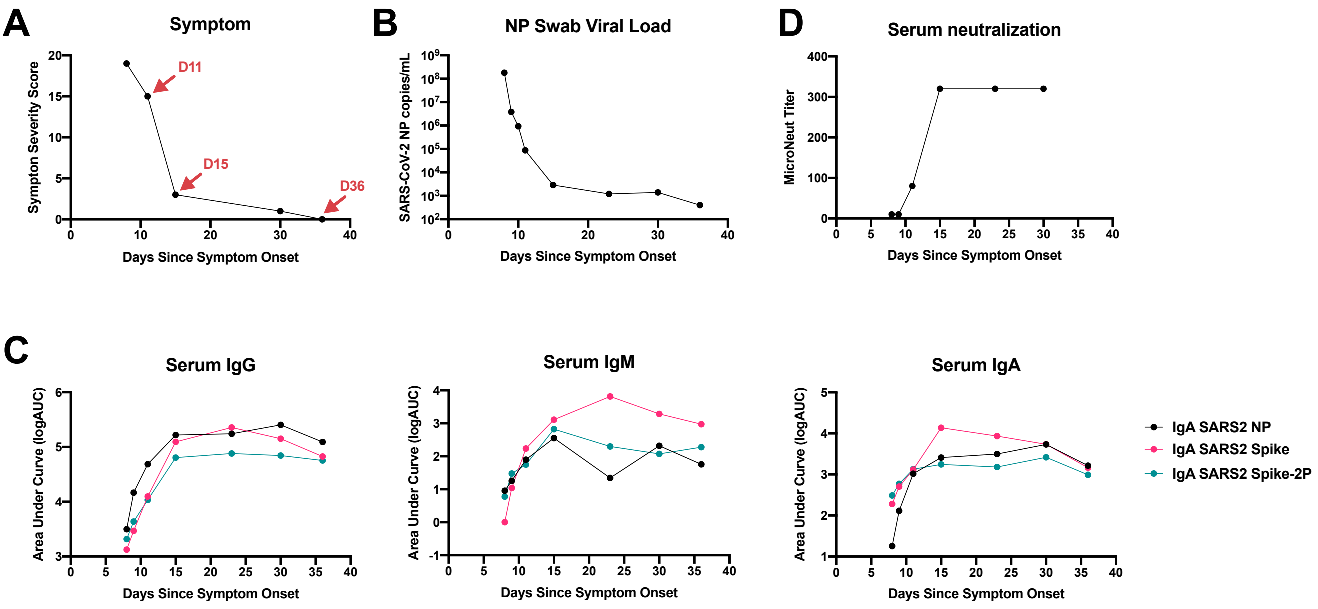

**Figure S1. Humoral responses and viral load of the COVID-19 convalescent donor.**
**(A)** Symptom severity scores of the COVID-19 convalescent donor. The method to determine severity score is in supplementary online material. Red arrows indicate the blood sampling time points that we used to isolate antibodies.
**(B)** Viral load from nasopharyngeal (NP) swabs.
**(C)** Serum IgG, IgM, IgA antibody binding**.**
**(D)** Serum micro-neutralization titer. Micro-Neutralization titers were defined as the highest serum dilution that neutralize all the virus, or 99% inhibitory concentration ($IC_{99}$).

### Figure S2

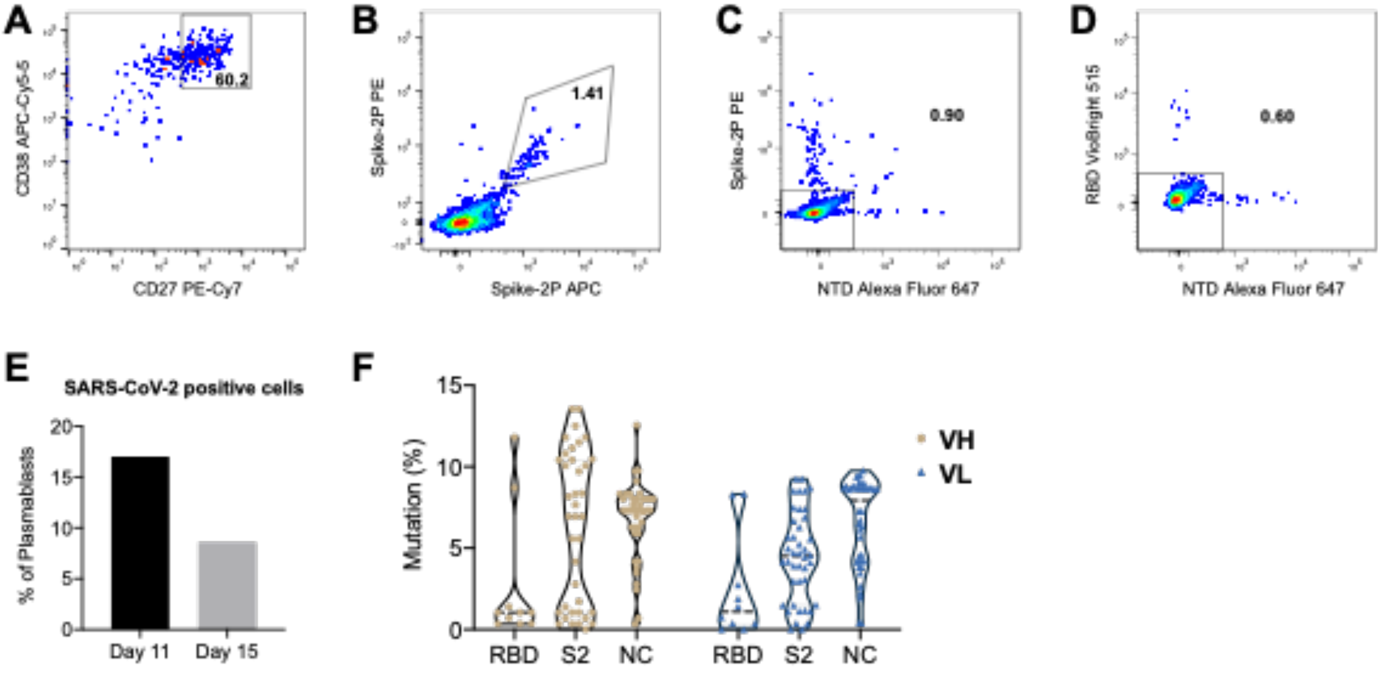

**Figure S2. Isolation of SARS-CoV-2-reactive antibodies from single cell-sorted plasmablasts and memory B cells.**

**(A)** Flow cytometry gating strategy for unbiased plasmablasts sorting. At day 11 and day 15 post onset of COVID-19 symptom, plasmablasts ($CD14^-/CD16^-/CD3^-/CD235a^-/CD19^+/CD20^{low}/IgD^-/CD27^{high}/CD38^{high}$) from a SARS-CoV-2 donor.

**(B-D)** Flow cytometry gating strategy for antigen specifc-memory B cells sorting. Antigen specific B cells from SARS-CoV-1 and SARS-CoV-2 donors were sorted with different combinations of the SARS-CoV-2 S-2P, RBD, NTD probes. Representative data for sorting Spike double positive (**B**), $Spike^+$ or $NTD^+$ (**C**), as well as $RBD^+$ or $NTD^+$ (**D**) subsets were shown.

**(E)** Percentages of SARS-CoV-2-reactive B cells in sorted plasmablasts from donor #1 Day 11 and Day 15 samples. Single-cells sorted plasmablasts that yielded productive $V_HD_HJ_H$ and $V_LJ_L$ gene pairs were considered as SARS-CoV-2-reactive plasmablasts.

**(F)** Mutation frequencies of $V_H$ and $V_L$ genes of Nucleocapsid (NC), RBD and S2 antibodies isolated from plasmablasts. Mutation frequency was analyzed and quality-controlled (expected error < 10) by Clonanalyst. Mean value and the mutation frequencies of each gene were shown in the violin plots.

### Figure S3

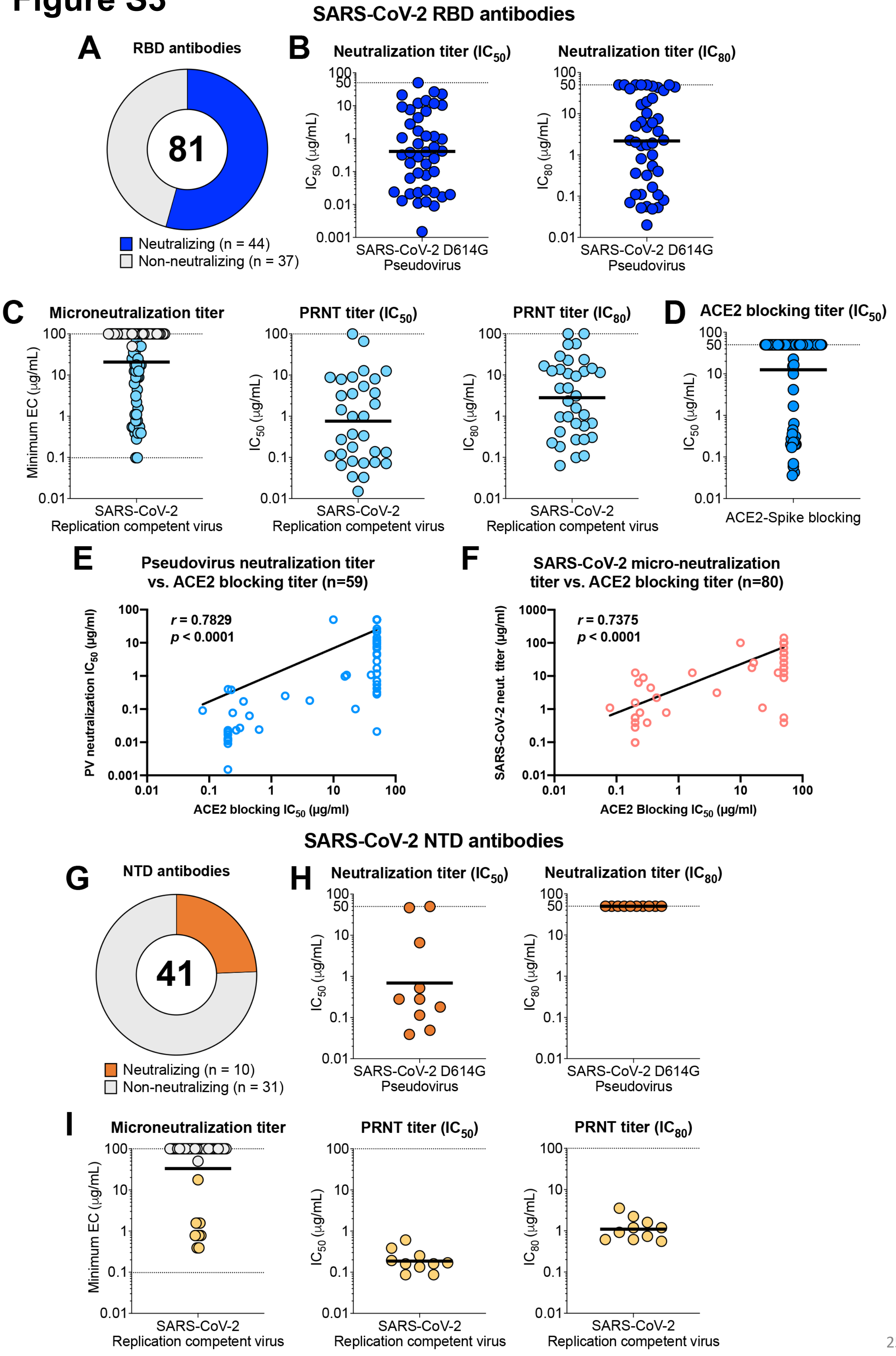
SARS-CoV-2 RBD antibodies
A
RBD antibodies
81
Neutralizing (n = 44)
Non-neutralizing (n = 37)
B
Neutralization titer (IC50)
IC50 (μg/mL)
SARS-CoV-2 D614G Pseudovirus
Neutralization titer (IC80)
IC80 (μg/mL)
SARS-CoV-2 D614G Pseudovirus
C
Microneutralization titer
Minimum EC (μg/mL)
SARS-CoV-2 Replication competent virus
PRNT titer (IC50)
IC50 (μg/mL)
SARS-CoV-2 Replication competent virus
PRNT titer (IC80)
IC80 (μg/mL)
SARS-CoV-2 Replication competent virus
D
ACE2 blocking titer (IC50)
IC50 (μg/mL)
ACE2-Spike blocking
E
Pseudovirus neutralization titer vs. ACE2 blocking titer (n=59)
r = 0.7829
p < 0.0001
PV neutralization IC50 (μg/ml)
ACE2 blocking IC50 (μg/ml)
F
SARS-CoV-2 micro-neutralization titer vs. ACE2 blocking titer (n=80)
r = 0.7375
p < 0.0001
SARS-CoV-2 neut. titer (μg/ml)
ACE2 Blocking IC50 (μg/ml)
SARS-CoV-2 NTD antibodies
G
NTD antibodies
41
Neutralizing (n = 10)
Non-neutralizing (n = 31)
H
Neutralization titer (IC50)
IC50 (μg/mL)
SARS-CoV-2 D614G Pseudovirus
Neutralization titer (IC80)
IC80 (μg/mL)
SARS-CoV-2 D614G Pseudovirus
I
Microneutralization titer
Minimum EC (μg/mL)
SARS-CoV-2 Replication competent virus
PRNT titer (IC50)
IC50 (μg/mL)
SARS-CoV-2 Replication competent virus
PRNT titer (IC80)
IC80 (μg/mL)
SARS-CoV-2 Replication competent virus

**Figure S3. Neutralization activities of the RBD and NTD antibodies.**
**(A-D)** Neutralization activity of RBD antibodies. **(A)** Proportion of SARS-CoV-2 RBD antibodies (n=81) that exhibited detectable neutralization in the microneutralization assay. **(B)** Neutralization $IC_{50}$ and $IC_{80}$ of RBD neutralizing antibodies (NAbs) against pseudotyped SARS-CoV-2. **(C)** Microneutralization titer, plaque reduction neutralization test (PRNT) $IC_{50}$ and $IC_{80}$ of RBD NAbs against replication-competent SARS-CoV-2. Microneutralization titer was defined as the lowest antibody concentration that neutralize all the virus, or 99% inhibitory concentration ($IC_{99}$). Antibodies with undetectable microneutralization titers are shown as gray symbols and nAbs are represented by blue symbols. **(D)** RBD NAbs blocking of ACE2 binding to SARS-CoV-2 Spike (S) protein. Blocking titer is shown as $IC_{50}$.
**(E-F)** Correlation analysis of RBD antibodies between neutralization and ACE2 blocking activities. Spearman correlation analysis were performed for (**E**) ACE2 blocking IC50 vs. PV neutralization IC50, as well as (**F**) for ACE2 blocking IC50 vs. SARS-CoV-2 neutralization titers (indicated by the lowest concentration that shows no CPE). Purified RBD antibodies in Table S1 and S2 that have pseudovirus neutralization data (n=59) or SARS-CoV-2 micro-neutralization assay data (n=80) were used in this analysis. P-value and r were indicated for each figures.
**(G-I)** Neutralization activity of NTD antibodies. (**G**) Proportion of SARS-CoV-2 NTD antibodies (n=41) that exhibited detectable neutralization in the microneutralization assay. (**H**) Neutralization $IC_{50}$ and $IC_{80}$ of NTD neutralizing antibodies against pseudotyped SARS-CoV-2. (**I**) Microneutralization titer, PRNT $IC_{50}$ and $IC_{80}$ of NTD neutralizing antibodies against replication-competent SARS-CoV-2. Antibodies with undetectable microneutralization titers are shown as gray symbols and neutralizing antibodies are represented by orange symbols. Horizontal bars represent the geometric means for each group of antibodies.

### Figure S4

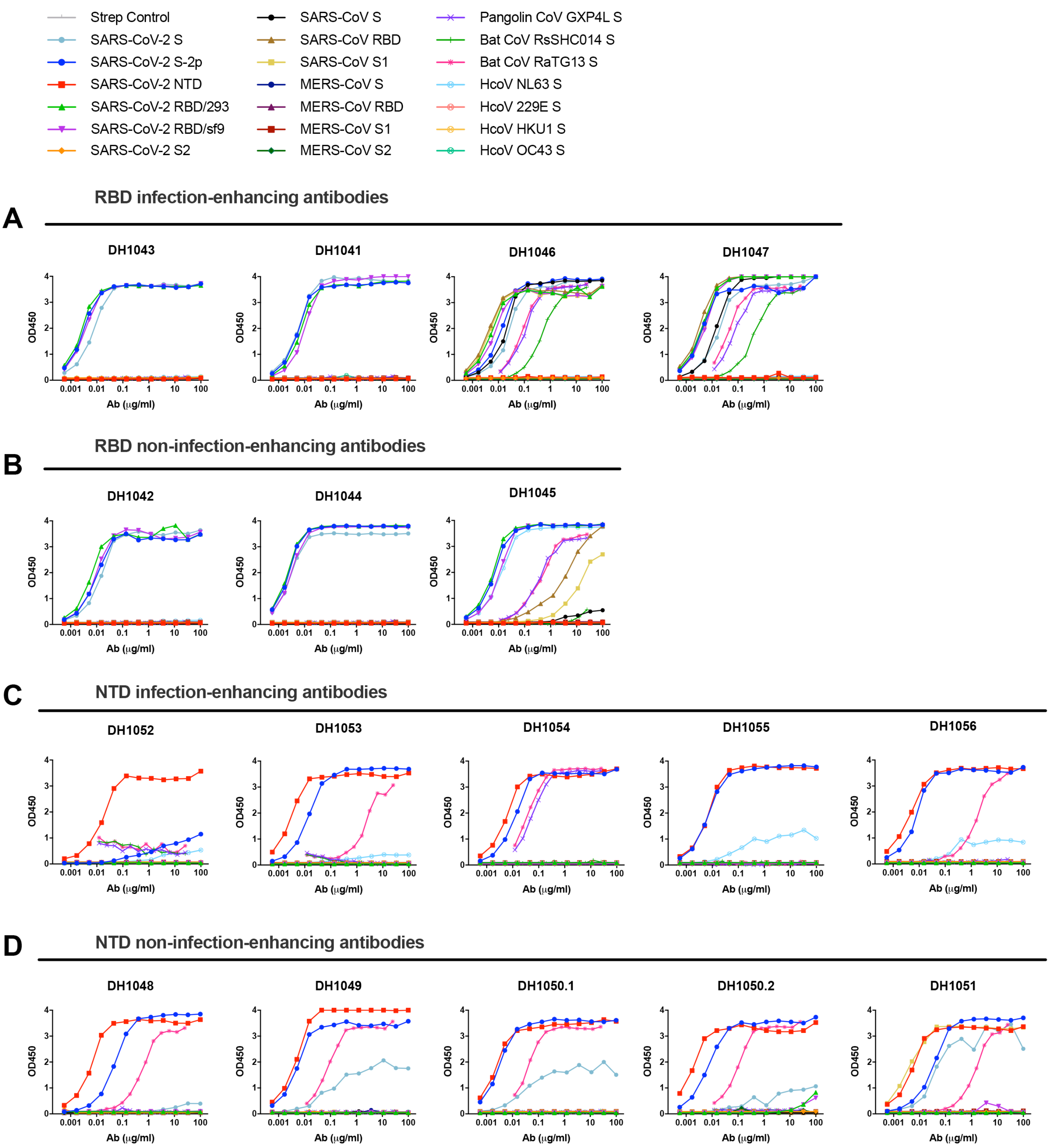

**Figure S4. ELISA binding curves of down-selected antibodies.** Different SARS-CoV-2 or other CoV viral antigens were coated on plates and detected with serial diluted (A) RBD infection-enhancing antibodies, (B) RBD non-infection-enhancing antibodies, (C) NTD infection-enhancing antibodies, and (D) NTD non-infection-enhancing antibodies.

### Figure S5

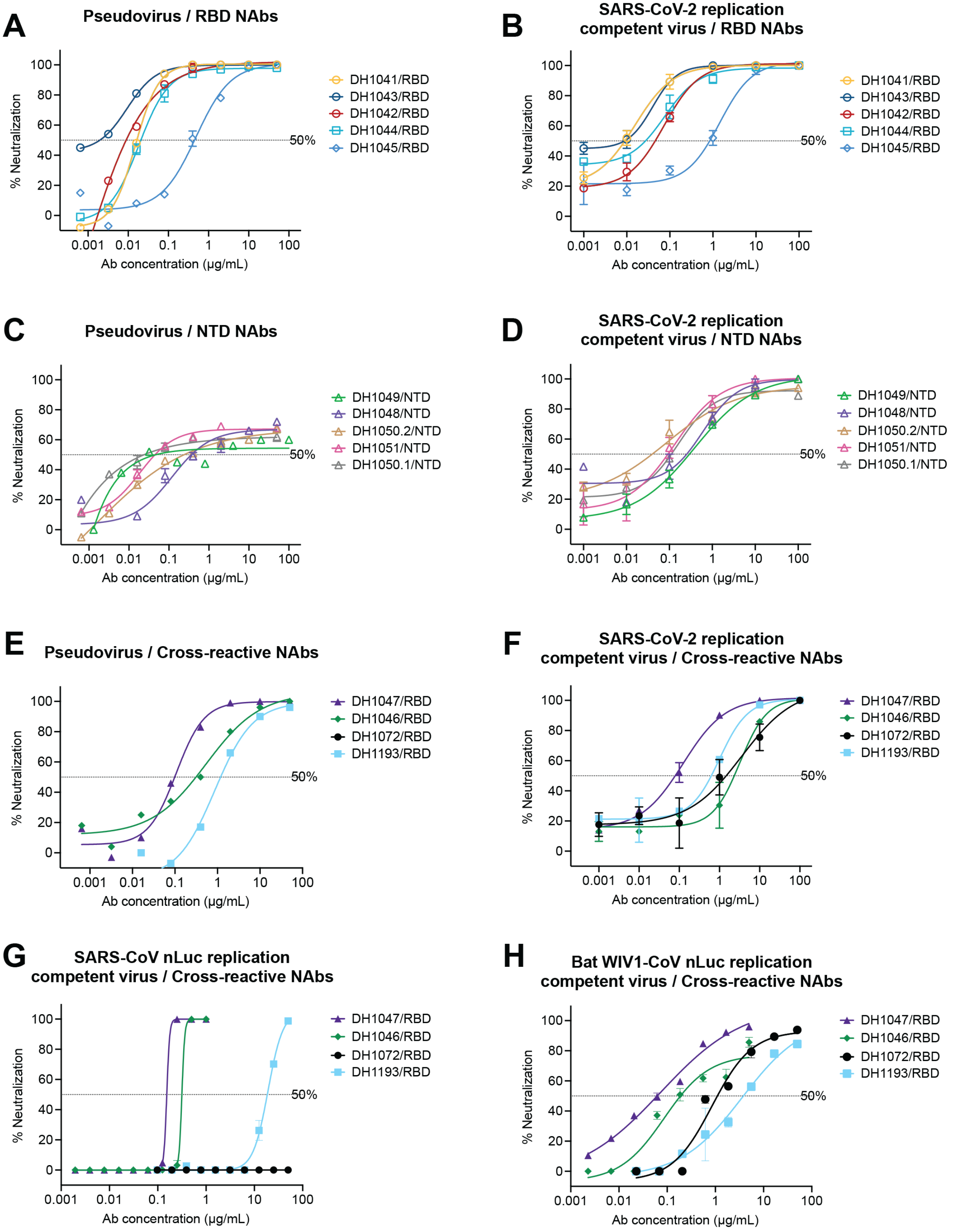

**Figure S5. Neutralization of SARS-CoV-2 and SARS-CoV-1 antibodies.**
**(A-B)** Neutralization curves for RBD antibodies against pseudotyped (A) and replication-competent (B) SARS-CoV-2.
**(C-D)** Neutralization curves for NTD antibodies against pseudotyped (C) and replication-competent (D) SARS-CoV-2.
**(E-H)** Neutralization curves for cross-neutralizing antibodies against pseudotyped (E) and replication-competent (F) SARS-CoV-2, SARS-CoV-1 nanoluciferase (nLuc) virus (G), and Bat WIV1-CoV nLuc virus (H).

### Figure S6

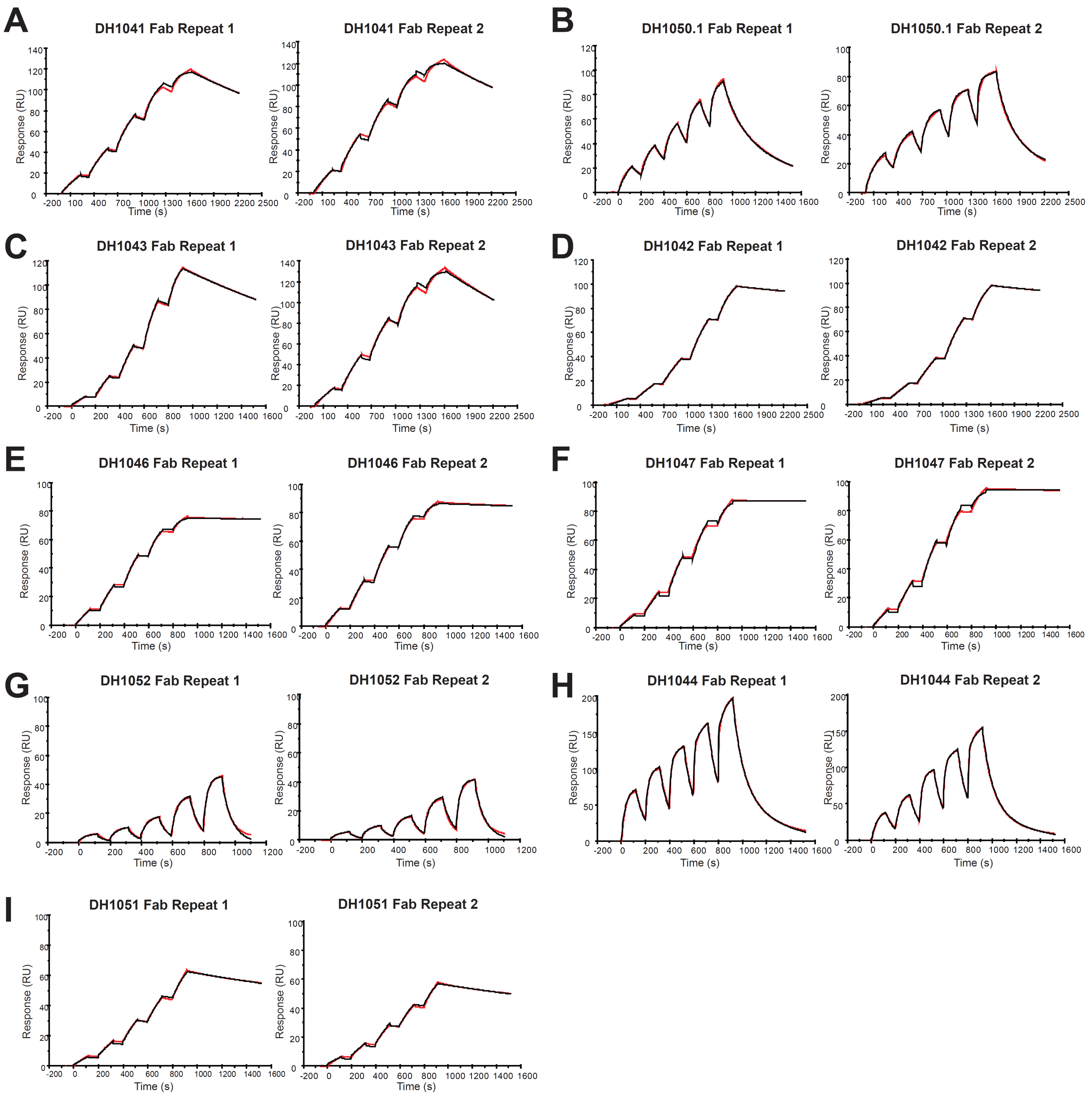

**Figure S6. Kinetics of RBD and NTD antibody Fabs on Spike-2P were measured by SPR.**

### Figure S7

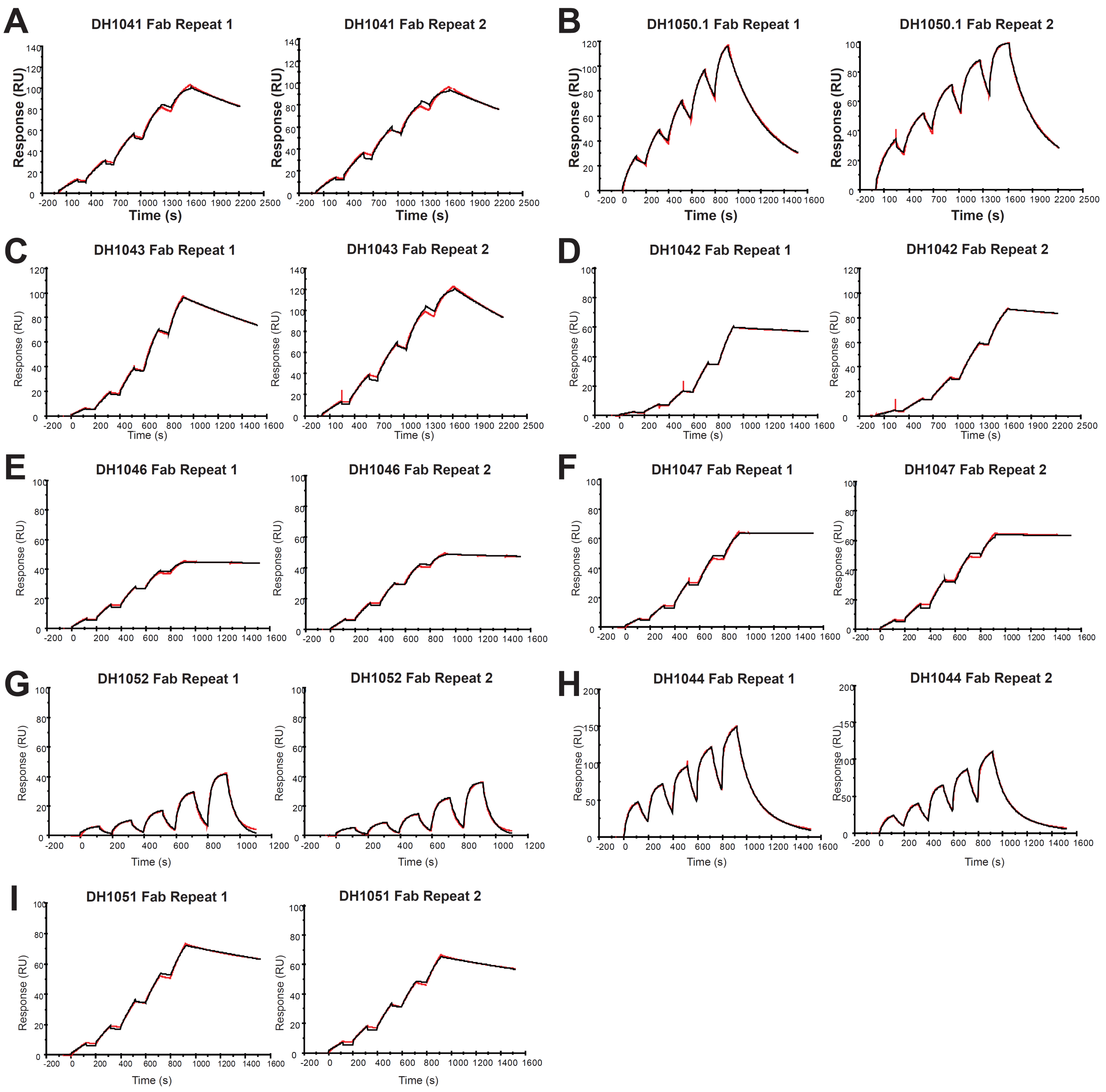

**Figure S7. Kinetics of RBD and NTD antibody Fabs on HexaPro were measured by SPR.**

### Figure S8

| Fab | Ligand | ka (1/Ms) | kd (1/s) | Kd (nM) |
|---|---|---|---|---|
| DH1041 | SARS-CoV-2 spike-2P | 1.27E+6 | 3.22E-4 | 0.3 |
| | SARS-CoV-2 spike-HexaPro | 9.31E+5 | 3.32E-4 | 0.4 |
| DH1042 | SARS-CoV-2 spike-2P | 1.61E+5 | 8.95E-5 | 0.6 |
| | SARS-CoV-2 spike-HexaPro | 9.63E+4 | 7.83E-5 | 0.8 |
| DH1043 | SARS-CoV-2 spike-2P | 2.65E+5 | 4.24E-4 | 1.6 |
| | SARS-CoV-2 spike-HexaPro | 2.21E+5 | 4.40E-4 | 2.0 |
| DH1044 | SARS-CoV-2 spike-2P | 1.45E+6 | 1.29E-2 | 9.0 |
| | SARS-CoV-2 spike-HexaPro | 1.27E+6 | 1.26E-2 | 9.9 |
| DH1046 | SARS-CoV-2 spike-2P | 9.78E+4 | 3.42E-5 | 0.3 |
| | SARS-CoV-2 spike-HexaPro | 8.54E+4 | 4.34E-5 | 0.5 |
| DH1047 | SARS-CoV-2 spike-2P | 1.57E+5 | <1.00E-5 | <0.1 |
| | SARS-CoV-2 spike-HexaPro | 2.87E+5 | 2.32E-5 | 0.08 |
| DH1050.1 | SARS-CoV-2 spike-2P | 2.70E+5 | 4.43E-3 | 16.4 |
| | SARS-CoV-2 spike-HexaPro | 4.54E+5 | 3.28E-3 | 7.2 |
| DH1051 | SARS-CoV-2 spike-2P | 1.12E+4 | 2.21E-4 | 19.8 |
| | SARS-CoV-2 spike-HexaPro | 1.16E+4 | 2.23E-4 | 19.2 |
| DH1052 | SARS-CoV-2 spike-2P | 5.36E+4 | 1.58E-2 | 294.0 |
| | SARS-CoV-2 spike-HexaPro | 6.06E+4 | 1.71E-2 | 281.8 |

**Figure S8. Affinity of RBD and NTD antibody Fabs.**

### Figure S9

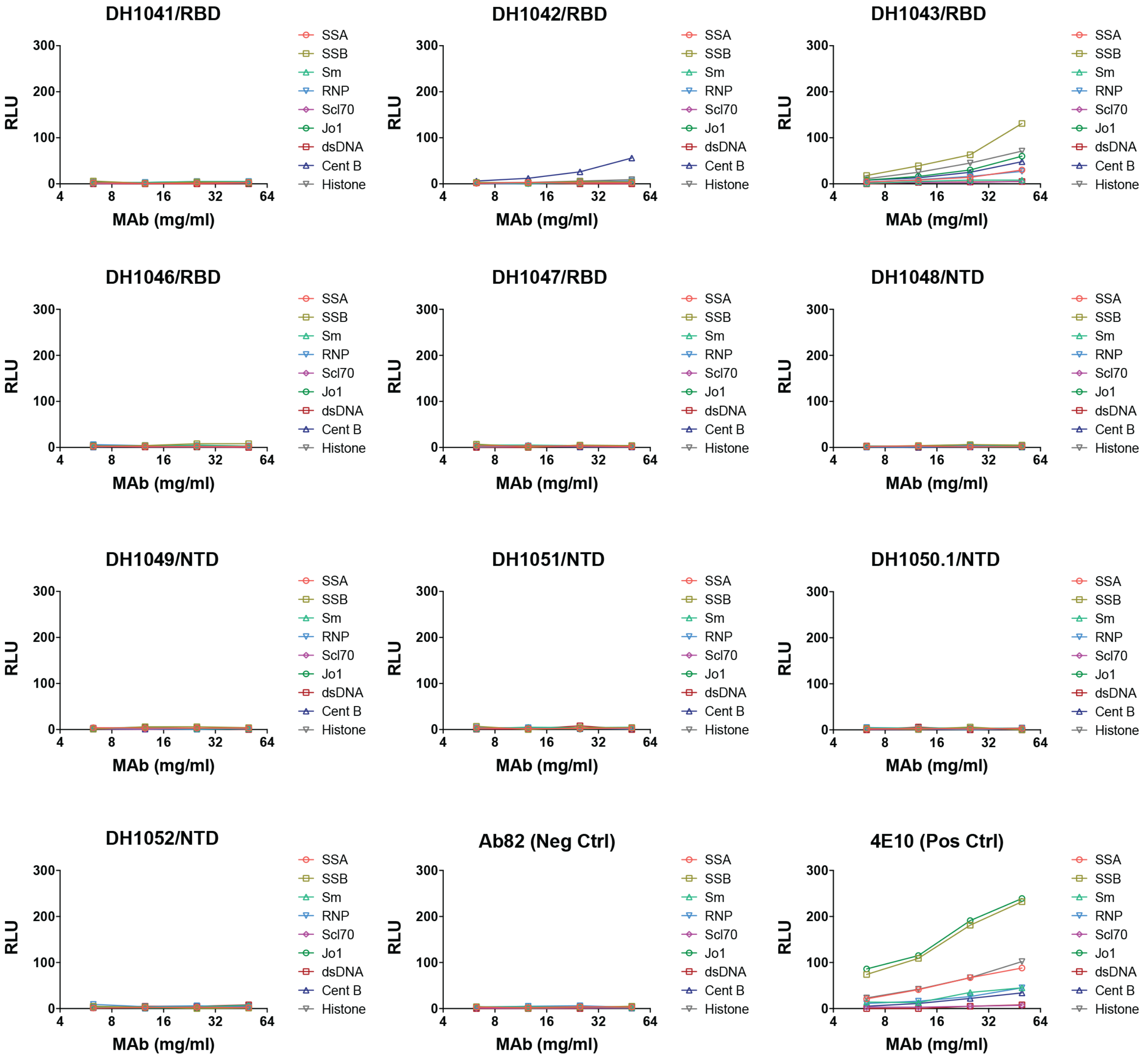

**Figure S9. Autoreactivity tests of SARS-CoV-2 RBD and NTD antibodies by AtheNA assay.** A panel of different antigens (SS/A, SS/B, Sm, RNP, Jo-1, Scl-70, dsDNA, Centromere B and Histone) were tested.

### Figure S10

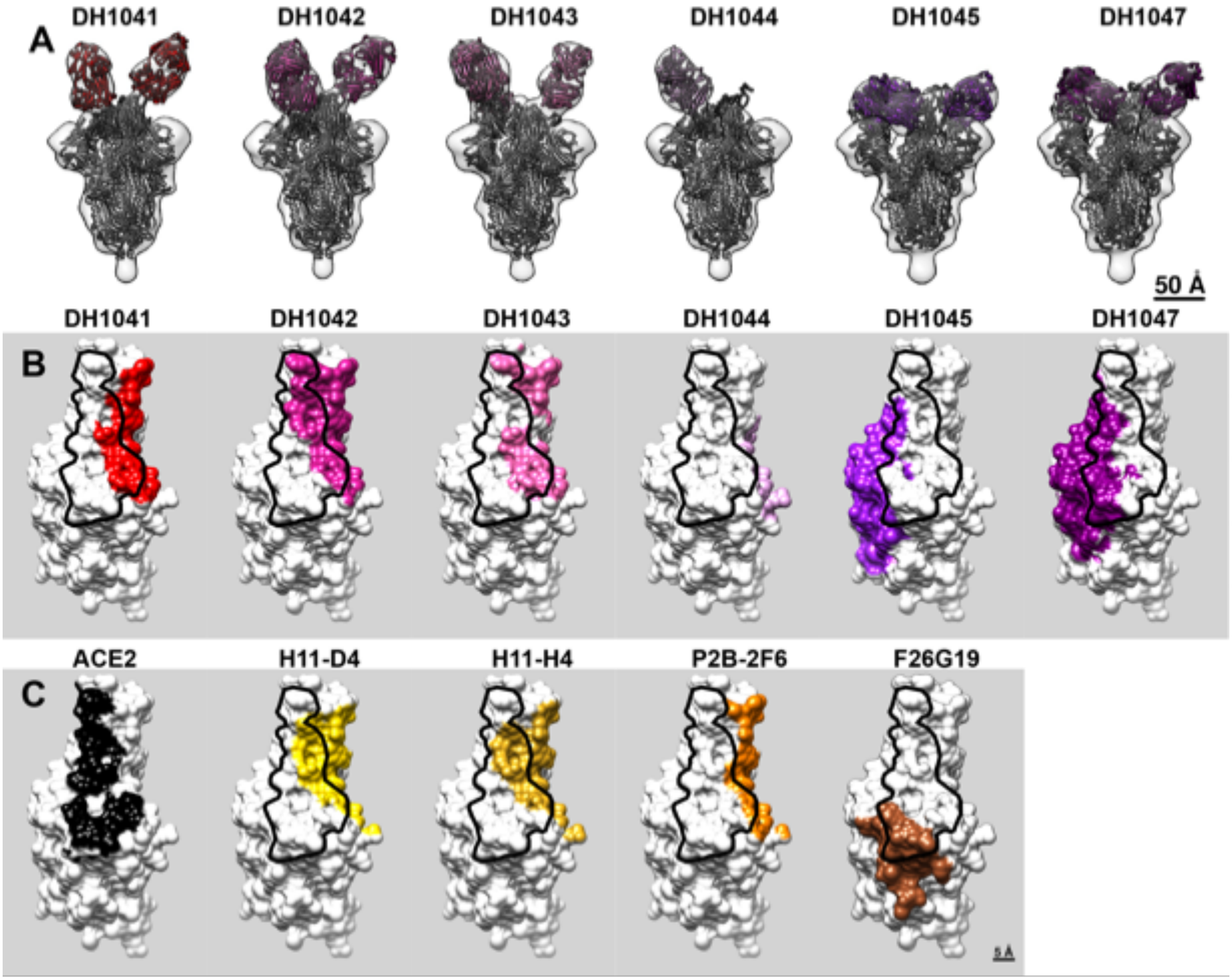

**Figure S10. Comparison of RBD epitopes from NSEM.**
**(A)** A spike model (PDB 6ZGE) and corresponding Fab homology models were manually docked and rigidly fit into each negative stain density map.
**(B)** The RBD of each model is enlarged and shown as a white surface, with the putative epitope of each antibody colored. Black outline indicates the ACE2 binding footprint.
**(C)** Comparison to ACE2 footprint and epitopes of three published antibodies with similar epitopes. See main text for references.

### Figure S11

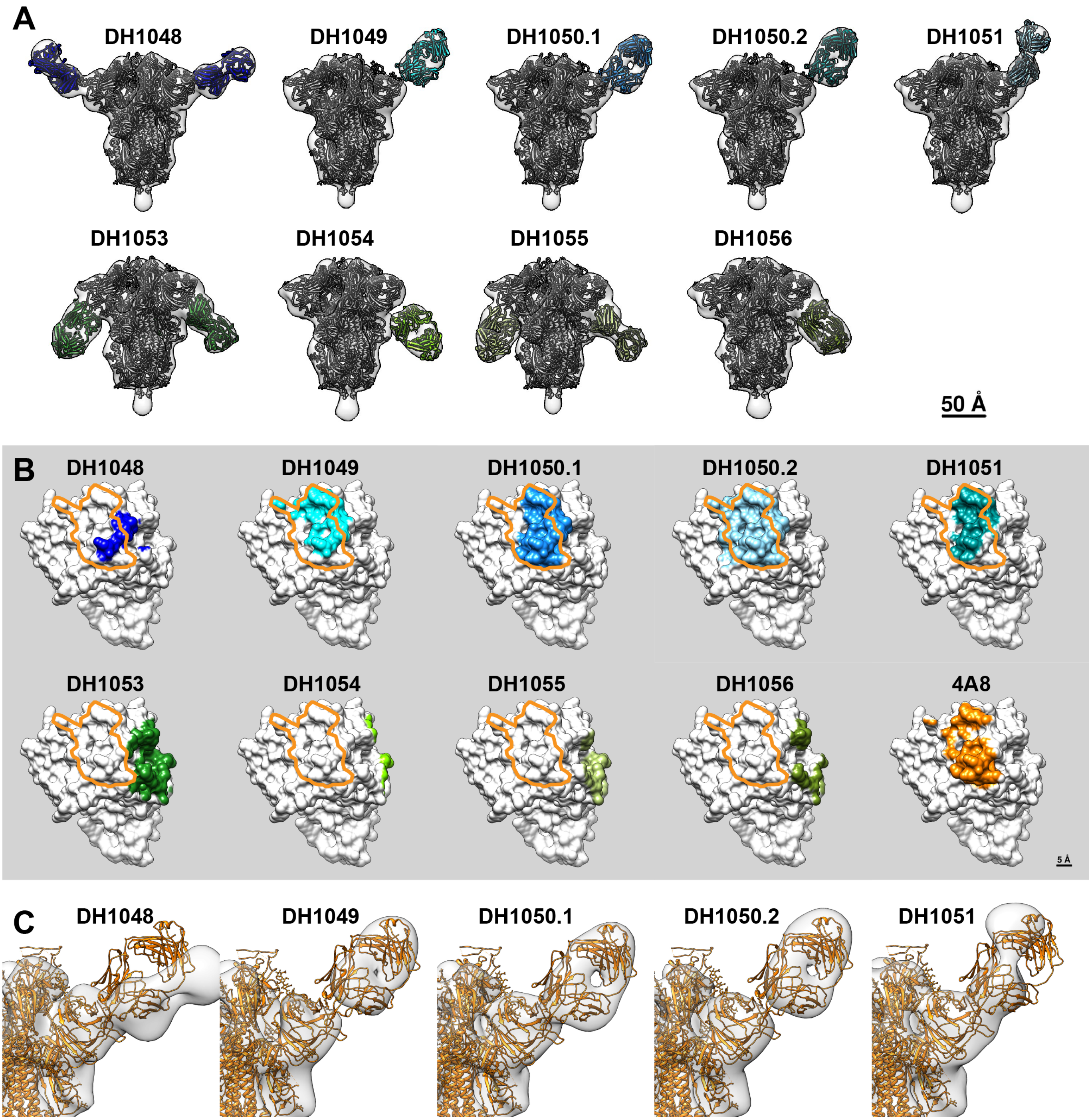

**Figure S11. Comparison of NTD epitopes from NSEM.**
**(A)** A spike model (PDB 6ZGE) and corresponding Fab homology models were manually docked and rigidly fit into each negative stain density map.
**(B)** The NTD of each model is enlarged and shown as a white surface, with the epitope of each antibody colored. Orange outline indicates the epitope of antibody 4A8, shown at bottom right. Outlines illustrate that the neutralizing antibodies DH1048-51 share the same epitope, whereas the infection-enhancing antibodies DH1053-56 bind a distinct epitope.
**(C)** The model of spike complex with Fab 4A8 (orange ribbons, PDG 7C2L) is rigidly fit into each of the NSEM maps (transparent surfaces). The close fit of 4A8 into DH1049, DH1050.1 and DH1050.2 indicate theses have the same approach angle as 4A8, whereas DH1048 and DH1051 have slightly different approaches.

### Figure S12

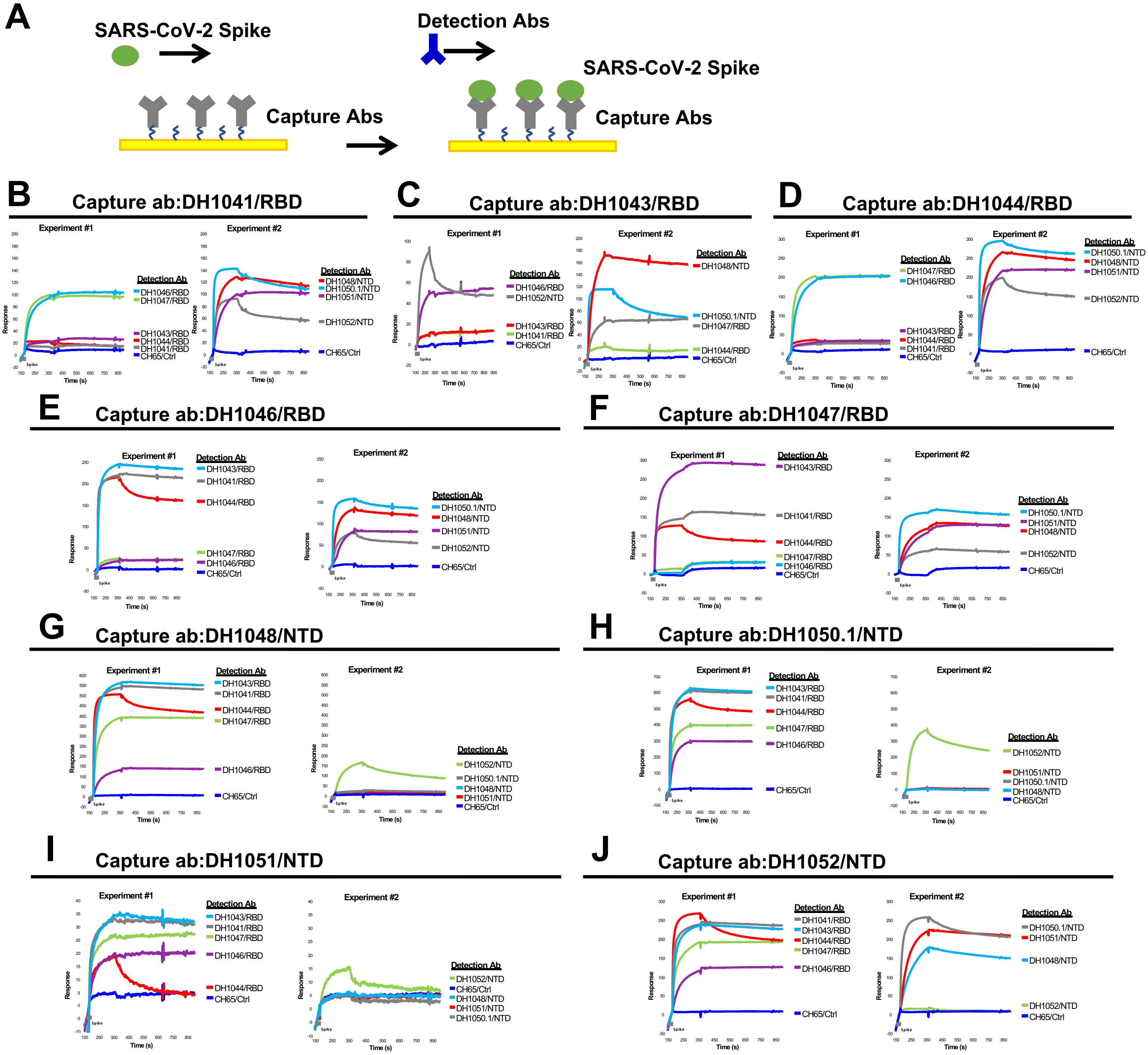

**Figure S12. Cross-blocking assay of RBD and NTD neutralizing antibodies (Related to Figure 3A).**

**(A)** Scheme of cross-blocking decection assay by SPR. Capture antibody was immoblized on sensor chip, followed by injection of SARS-CoV-2 Spike and then immediately injection of detection antibodies. Human antibody CH65 was used as negative control antibody in all SPR experiments.

**(B-D)** RBD antibodies DH1041 (B), DH1043 (C), or DH1044 (D) was immoblized as capture antibodies to test cross-blocking with different detection antibodies. Three independent experiments were shown.

**(E,F)** RBD cross-reactive antibodies DH1046 (E) or DH1046 (F) was immoblized as capture antibodies to test cross-blocking with different detection antibodies as indicated.

**(G-J)** NTD antibodies DH1048 (G), DH1050.1 (H), DH1051 (I), or NTD ADE antibody DH1052 (J) was immoblized as capture antibodies to test cross-blocking with different detection antibodies. Two or three independent experiments were shown.

### Figure S13

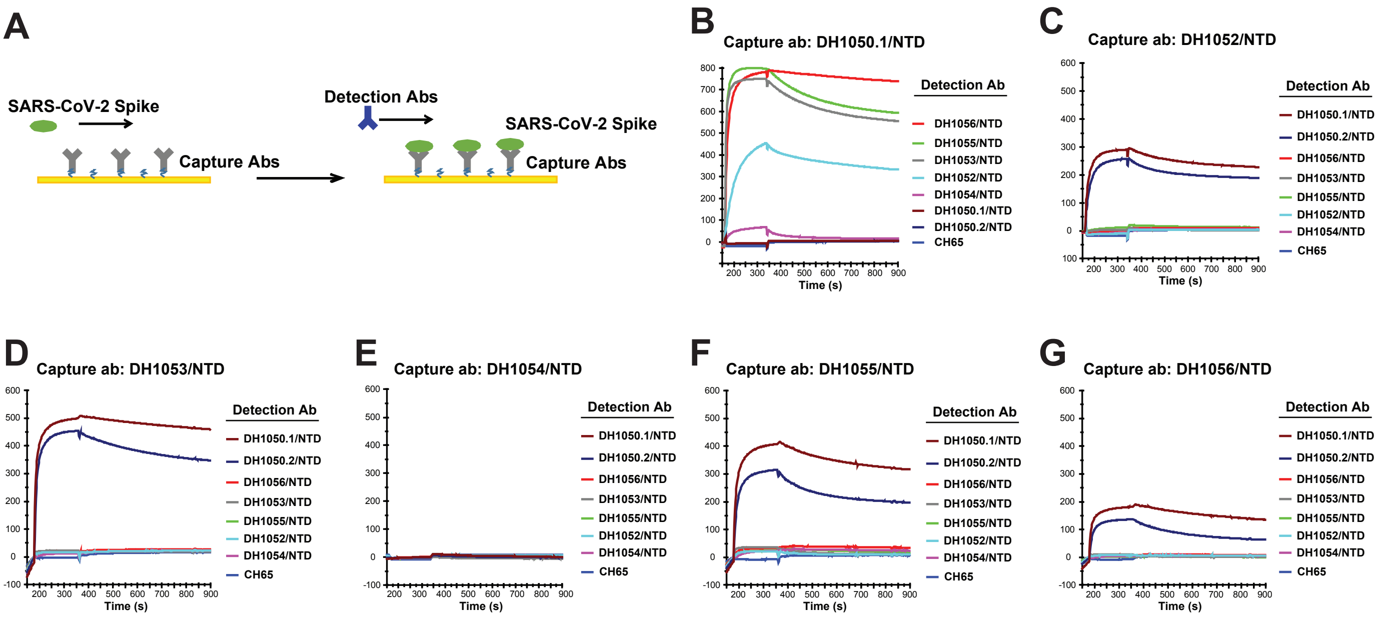

**Figure S13. Cross-blocking assay of RBD and NTD neutralizing antibodies (Related to Figure 3C).**

**(A)** Scheme of cross-blocking decection assay by SPR. Capture antibody was immoblized on sensor chip, followed by injection of SARS-CoV-2 Spike and then immediately injection of detection antibodies. Human antibody CH65 was used as negative control antibody in all SPR experiments.

**(B-G)** SPR cross-blocking assays. NTD NAb DH1050.1 (B), NTD infection-enhancing abs DH1052 (C), DH1053 (D), DH1054 (E), DH1055 (F) or DH1056 (G) was immoblized as capture antibodies to test cross-blocking with different detection antibodies.

### Figure S14

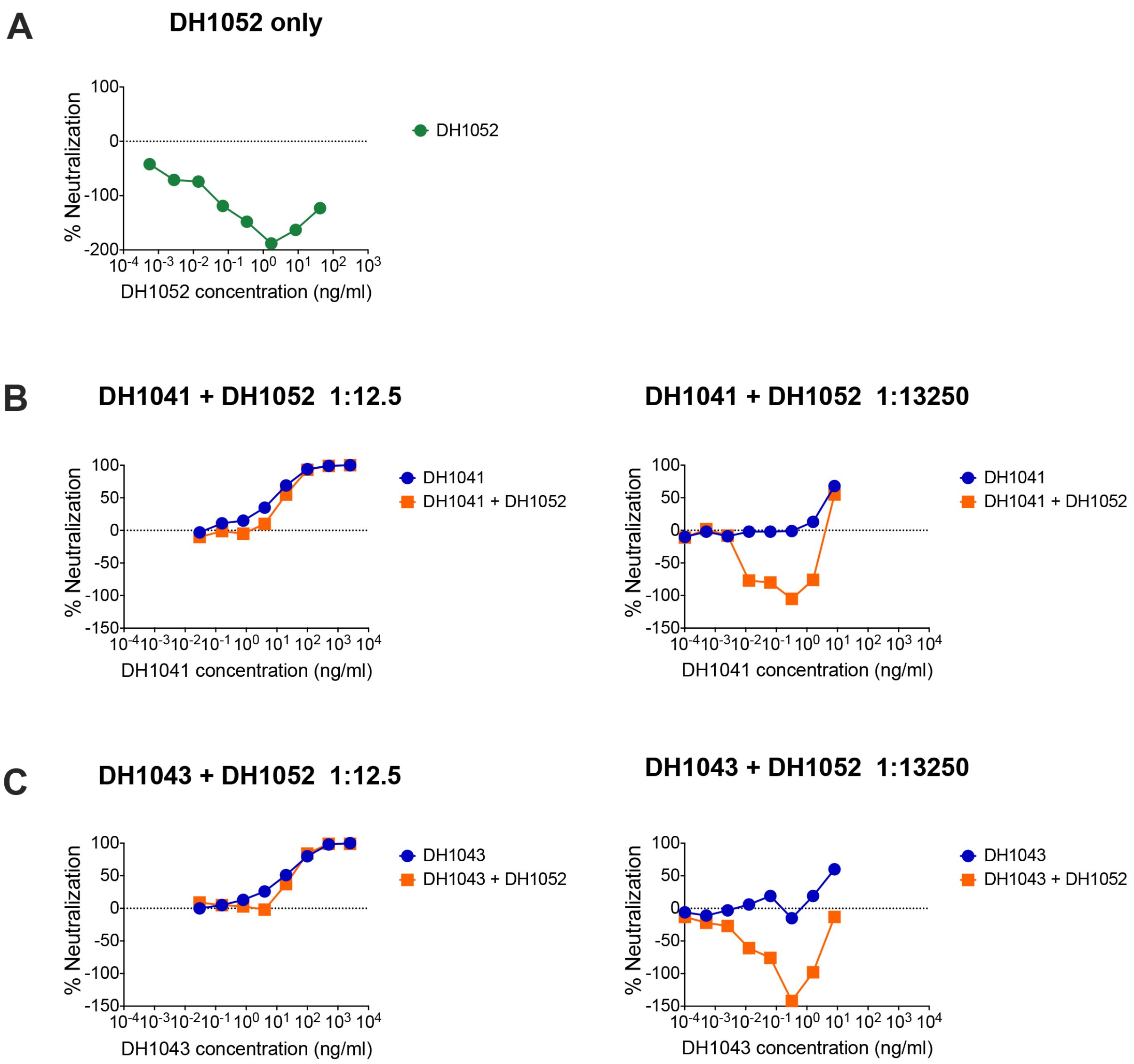

**Figure S14. Effect of combining infection-enhancing RBD and NTD antibodies on SARS-CoV-2 pseudovirus infection in ACE2-expressing cells (related to Figure 3G-H).** The infection-enhancing NTD antibody DH1052 was tested alone **(A)** or mixed with infection-enhancing RBD antibodies DH1041 **(B)** or DH1043 **(C)** in 1:12.5 ratio or 1:13250 ratio, respectively. The NTD:RBD antibody mixtures (orange), as well as RBD antibody alone (blue), were five-fold serially diluted and tested for neutralization against SARS-CoV-2 D614G pseudoviruse in 293T/ACE2 cells.

### Figure S15

| | 24MFK DH1041 | 30MFK DH1052 | 35MFK DH1047 | 113KJ DH1043 | 26MFK DH1050.1 |
|---|---|---|---|---|---|
| **Data Collection** | | | | | |
| Microscope | FEI Titan Krios | FEI Titan Krios | FEI Titan Krios | FEI Titan Krios | FEI Titan Krios |
| Voltage (kV) | 300 | 300 | 300 | 300 | 300 |
| Electron dose ($e^-/Å^2$) | 65.94 | 66.71 | 66.77 | 66.77 | 65.09 |
| Detector | Gatan K3 | Gatan K3 | Gatan K3 | Gatan K3 | Gatan K3 |
| Pixel Size (Å) | 1.058 | 1.058 | 1.058 | 1.058 | 1.058 |
| Defocus Range (μm) | ~0.75-2.50 | ~0.75-2.50 | ~0.75-2.50 | ~0.75-2.50 | ~0.75-2.50 |
| Magnification | 81000 | 81000 | 81000 | 81000 | 81000 |
| Micrographs collected | 7362 | 9375 | 3440 | 1900 | 2154 |
| **Reconstruction** | | | | | |
| Software | cryoSparc | cryoSparc | cryoSparc | cryoSparc | cryoSparc |
| Particles | 151,384 | 143,4115 | 127,401 | 133,814 | 426,025 |
| Symmetry | C1 | C1 | C1 | C1 | C1 |
| Box size (pix) | 350 | 350 | 350 | 350 | 350 |
| Resolution (Å) (FSC 0.143)* | 3.42 | 2.97 | 3.4 | 3.66 | 3.35 |
| **Refinement (Phenix)** | | | | | |
| Protein residues | 2994 | 4194 | 4251 | 4212 | 4296 |
| Chimera CC | 0.72 | 0.77 | 0.61 | 0.52 | 0.73 |
| R.m.s. deviations | | | | | |
| Bond lengths (Å) | 0.013 | 0.012 | 0.012 | 0.012 | 0.013 |
| Bond angles (°) | 2.229 | 1.956 | 1.945 | 1.918 | 2.006 |
| Validation | | | | | |
| Molprobity score | 1.71 | 1.16 | 1.41 | 1.40 | 1.53 |
| Clash score | 0.81 | 0.18 | 0.22 | 1.19 | 0.42 |
| Favored rotamers (%) | 97.06 | 98.43 | 97.89 | 99.53 | 97.48 |
| Ramachandran | | | | | |
| Favored regions (%) | 87.95 | 93.07 | 88.75 | 89.27 | 89.05 |
| Allowed regions (%) | 10.58 | 6.45 | 9.62 | 9.57 | 9.36 |
| Disallowed regions (%) | 1.47 | 0.49 | 1.64 | 1.16 | 1.59 |
| EMRinger score | 2.73 | 3.25 | 2.78 | 1.0 | 2.09 |

*Resolutions are reported according to the FSC 0.143 gold-standard criterion

**Figure S15. Cryo-EM data collection and refinement statistics.**

### Figure S16

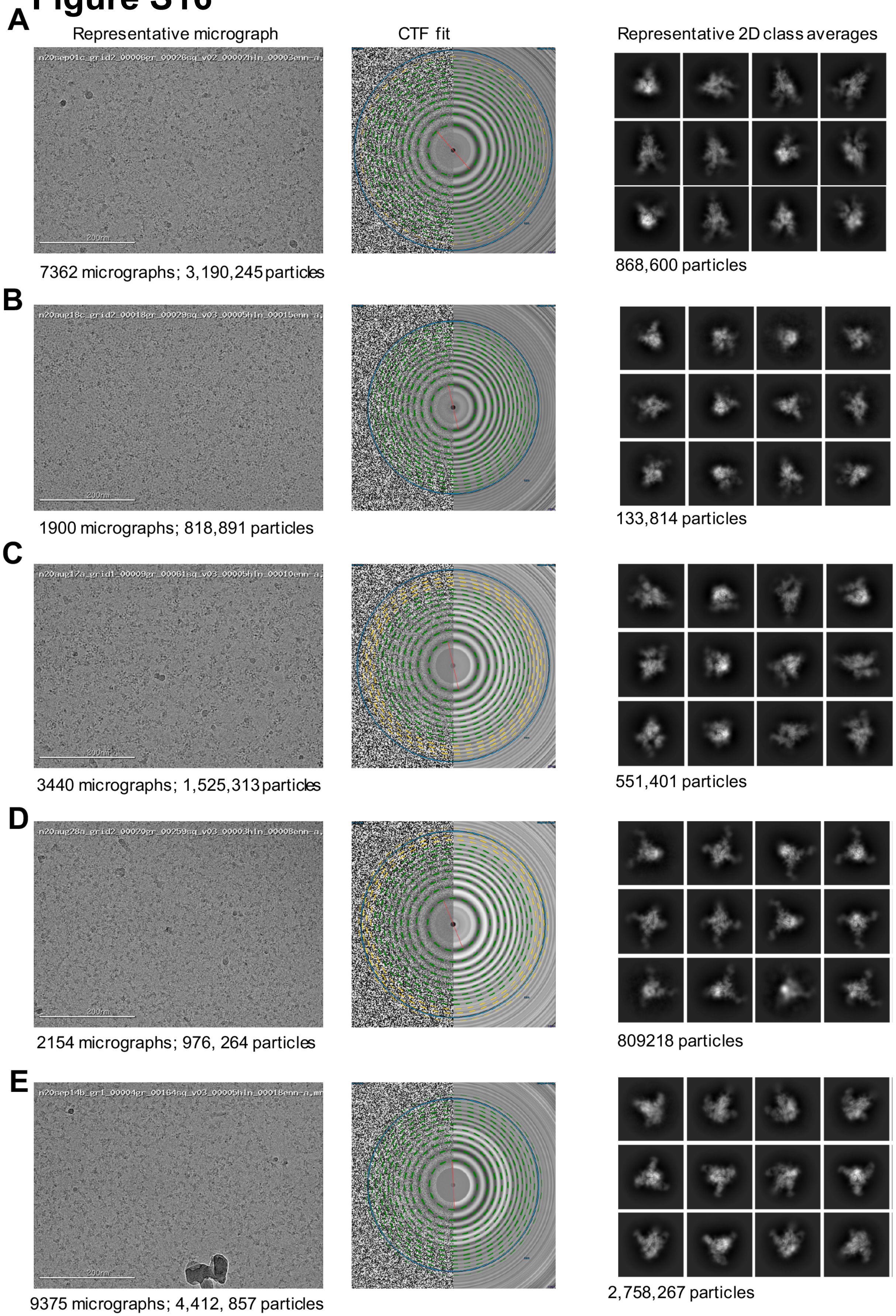

**Figure S16. Cryo-EM data processing details**. (left) Representative micrograph, (middle) CTF fit and (right) Representative 2D class averages for (A) DH1041-Spike-2P (S2P) complex, (B) DH1043-S2P complex, (C) DH1047-S2P complex, (D) DH1050.1-S2P complex, (E) DH1052-S2P complex.

### Figure S17

**A** DH1041 Fab bound to S-2P with 1 RBDs in “up” position

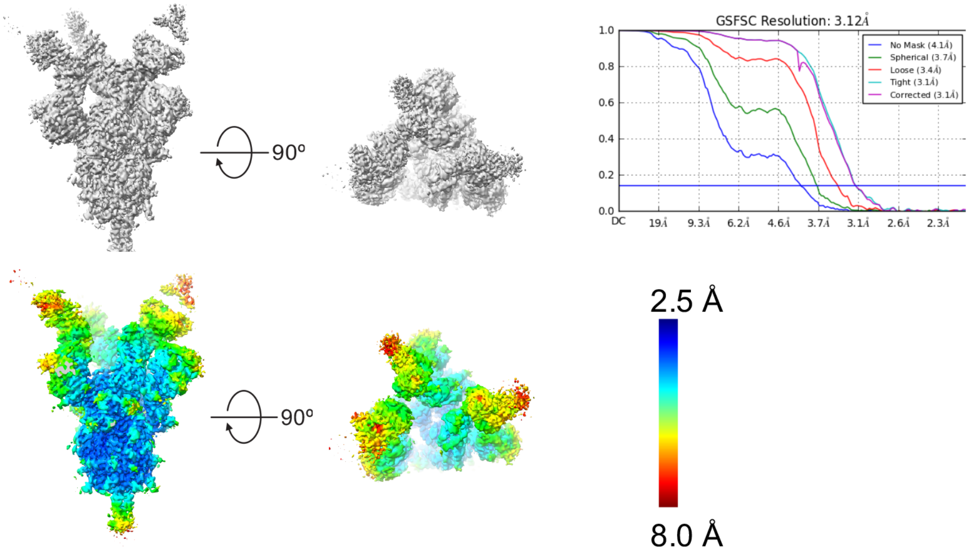

**B** DH1041 Fab bound to S-2P with 2 RBDs in “up” positions

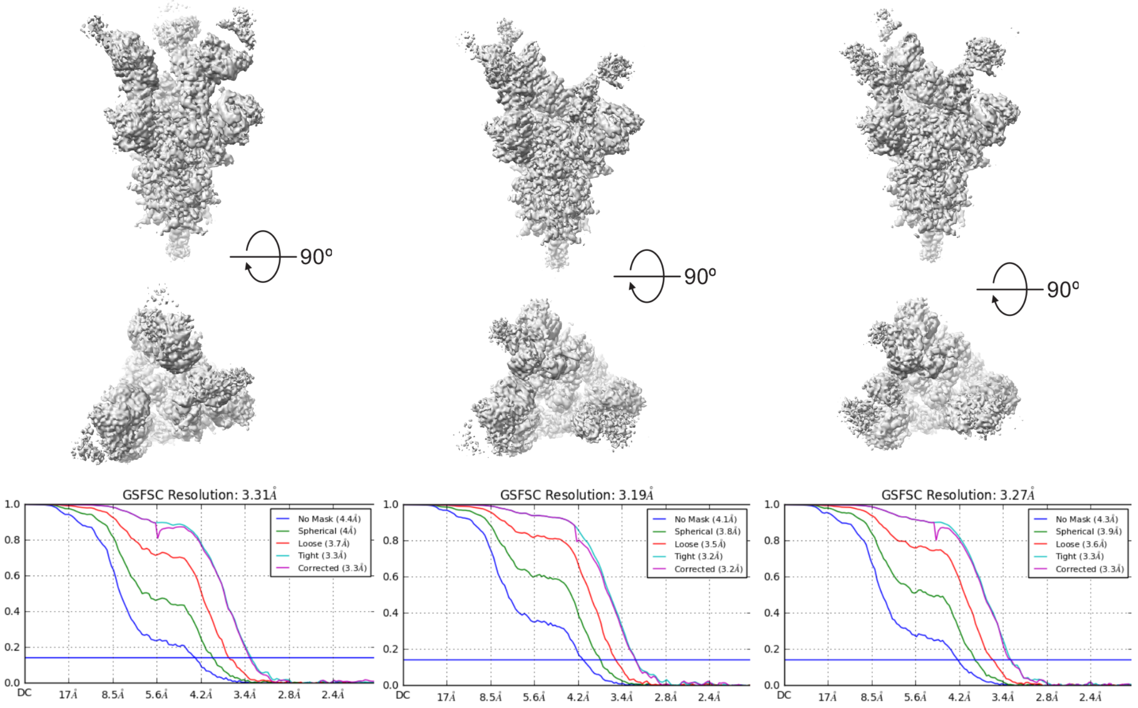

**C** DH1041 Fab bound to S-2P with 3 RBDs in “up” positions

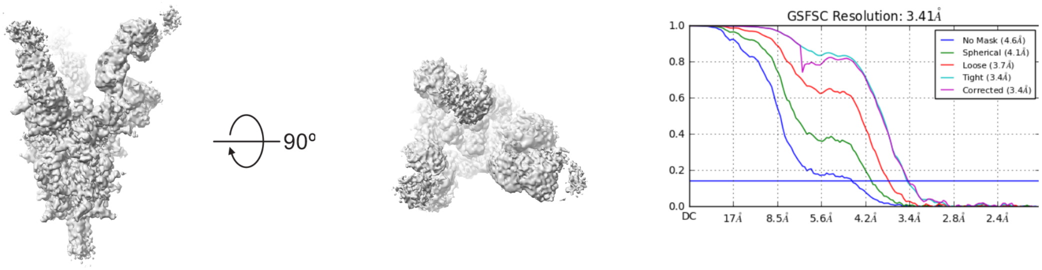

**Figure S17. Global and Local map resolutions for DH1041/S-2P complex.** (A) Cryo-EM reconstruction of DH1041 bound to 1-RBD-up 2P spike. (B) Cryo-EM reconstruction of DH1041 bound to 2-RBD-up 2P spike. (C) Cryo-EM reconstruction of DH1041 bound to 3-RBD-up 2P spike.

### Figure S18

**A** DH1043 Fab bound to S-2P with 1 RBD in “up” position

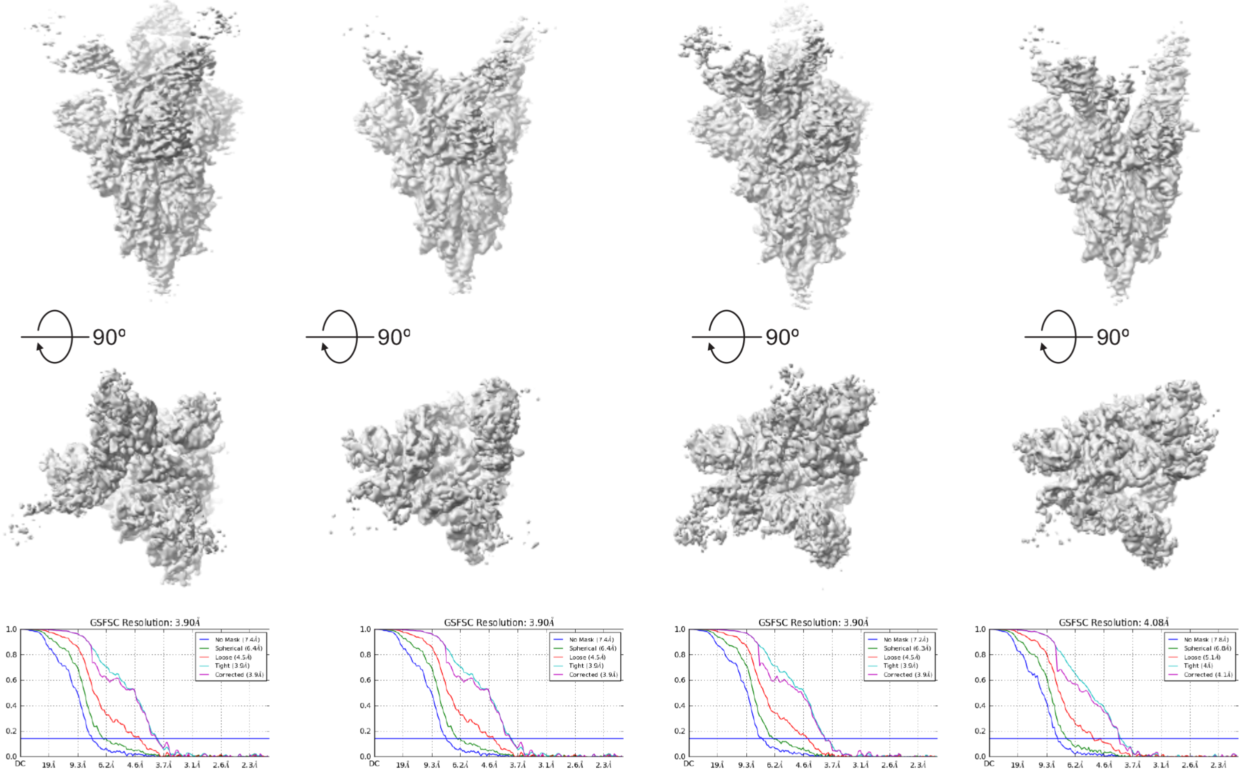

**B**

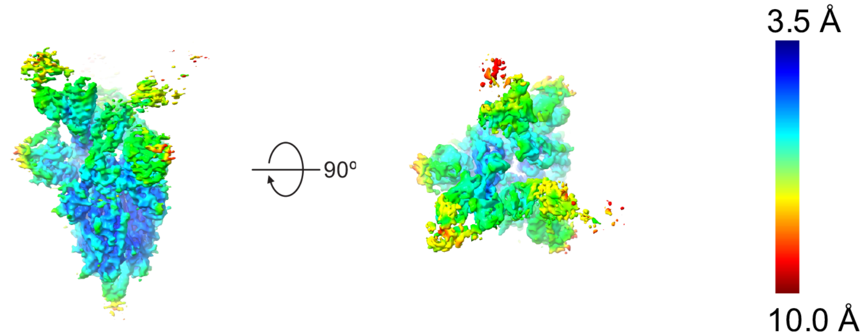

**C**

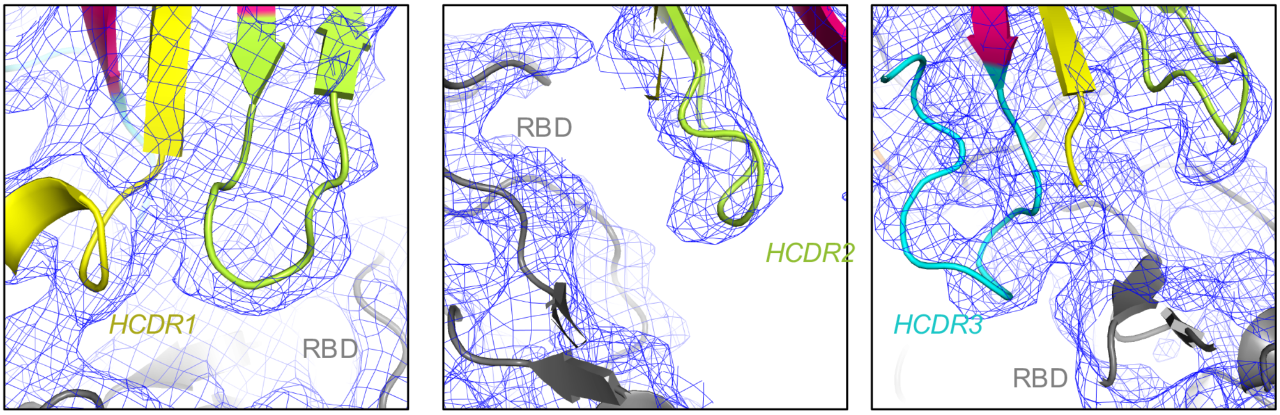

**Figure S18. Global and Local map resolutions for DH1043/S-2P complex.** (A) Cryo-EM reconstructions of DH1043 bound to 1-RBD-up 2P spike Top two rows show refined maps, bottom row shows the FSC curve for each corresponding map. (B) Refined cryo-EM map that was used for model building colored by local resolution.(C) Zoomed-in view of the DH1043 interface with RBD. The cryo-EM map is shown as a blue mesh with underlying fitted model shown in cartoon representation, with the DH1047 HCDR1 loop colored yellow, HCDR2 colored limon and HCDR3 cyan.

### Figure S19

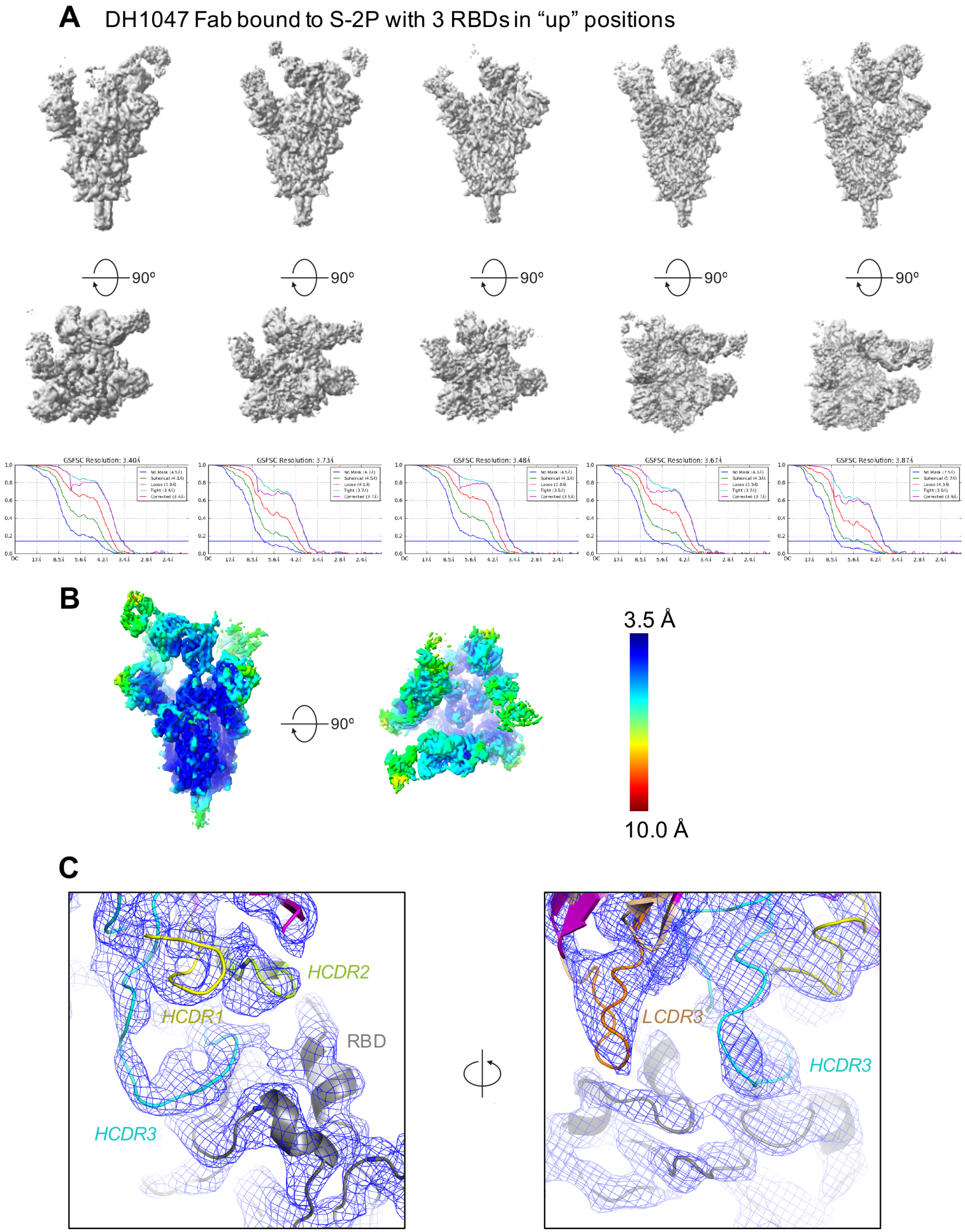
A
DH1047 Fab bound to S-2P with 3 RBDs in “up” positions
90º
GSFSC Resolution: 3.40Å
GSFSC Resolution: 3.73Å
GSFSC Resolution: 3.48Å
GSFSC Resolution: 3.67Å
GSFSC Resolution: 3.87Å
B
90º
3.5 Å
10.0 Å
C
HCDR2
HCDR1
RBD
HCDR3
LCDR3
HCDR3

**Figure S19. Global and Local map resolutions for DH1047/S-2P complex.** (A) Cryo-EM reconstructions of DH1047 bound to 3-RBD-up 2P spike. Top two rows show refined maps, bottom row shows the FSC curve for each corresponding map. (B) Refined cryo-EM map that was used for model building colored by local resolution. (C) Zoomed-in view of the DH1047 interface with RBD. The cryo-EM map is shown as a blue mesh with underlying fitted model shown in cartoon representation, with the DH1047 HCDR1 loop colored yellow,HCDR2 colored limon, HCDR3 cyan and LCDR1 orange.

### Figure S20

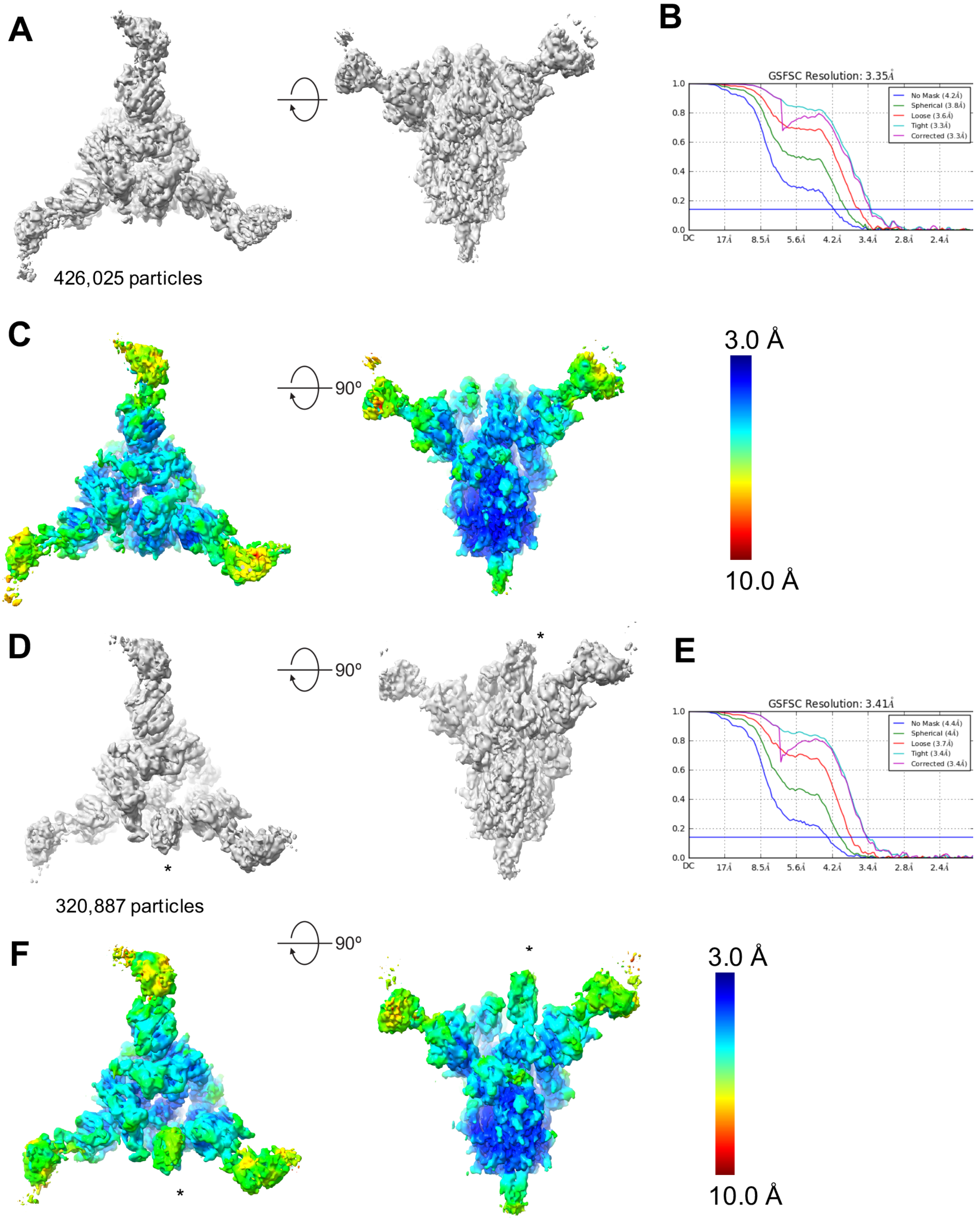

**Figure S20. Global and Local map resolutions for DH1050.1/S-2P complex.** (A) Cryo-EM reconstruction of DH1050.1 bound to 3-RBD-down 2P spike. (B) Fourier shell correlation curves. (C) Refined cryo-EM map colored by local resolution for the DH1050.1 bound to 3-RBD-down 2P spike. (D) Cryo-EM reconstruction of DH1050.1 bound to 1-RBD-up 2P spike. (E) Fourier shell correlation curves. (F) Refined cryo-EM map colored by local resolution for the DH1050.1 bound to 1-RBD-up 2P spike.

### Figure S21

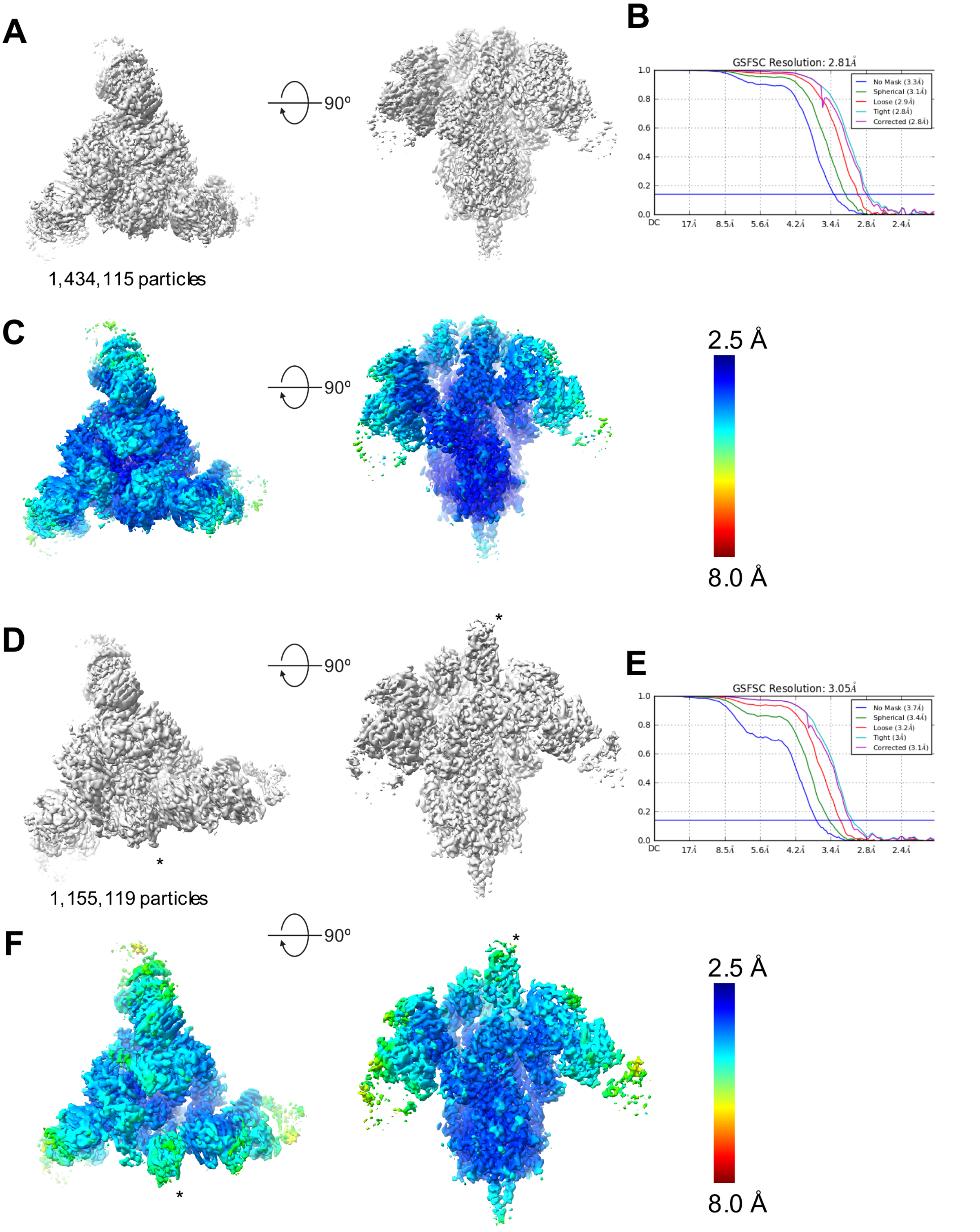

**Figure S21. Global and Local map resolutions for DH1052/S-2P complex.** (A) Cryo-EM reconstruction of DH1052 bound to 3-RBD-down stabilized Spike "2P" (S-2P). (B) Fourier shell correlation curves. (C) Refined cryo-EM map colored by local resolution for the DH1052 bound to 3-RBD-down S-2P. (D) Cryo-EM reconstruction of DH1052 bound to 1-RBD-up S-2P. (E) Fourier shell correlation curves. (F) Refined cryo-EM map colored by local resolution for the DH1052 bound to 1-RBD-up S-2P.

### Figure S22

| Group # | Animal # | Inflammation | | | | SARS-CoV-2 antigen expression | | | |
|---|---|---|---|---|---|---|---|---|---|
| | | Lc | Rm | Rc | Sum | Lc | Rm | Rc | Sum |
| Group1 DH1043 | 2025B | 0 | 1 | 0 | 1 | 0 | 0 | 0 | 0 |
| | BB174A | 1 | 0 | 1 | 2 | 0 | 0 | 0 | 0 |
| | CS054 | 1 | 1 | 1 | 3 | 0 | 0 | 0 | 0 |
| | AE969HA | 2 | 1 | 2 | 5 | 0 | 0 | 0 | 0 |
| | C0574 | 1 | 1 | 1 | 3 | 0 | 0 | 0 | 0 |
| Group2 DH1050.1 | T49PB | 0 | 0 | 1 | 1 | 0 | 0 | 0 | 0 |
| | VL50L | 1 | 0 | 0 | 1 | 0 | 0 | 0 | 0 |
| | VL75L | 0 | 0 | 0 | 0 | 0 | 0 | 0 | 0 |
| | P286JM | 0 | 0 | 1 | 1 | 0 | 0 | 0 | 0 |
| | P336IK | 0 | 0 | 0 | 0 | 0 | 0 | 1 | 1 |
| Group3 DH1046 | BC589A | 1 | 1 | 1 | 3 | 1 | 0 | 0 | 1 |
| | BB546A | 1 | 0 | 0 | 1 | 0 | 1 | 0 | 1 |
| | 2019B | 3 | 2 | 3 | 8 | 1 | 0 | 1 | 2 |
| | BB498A | 0 | 0 | 2 | 2 | 0 | 0 | 0 | 0 |
| | BF791E | 2 | 4 | 4 | 10 | 0 | 0 | 0 | 0 |
| Group4 CH65 Control | 2021C | 1 | 2 | 2 | 5 | 2 | 0 | 1 | 3 |
| | BB167A | 1 | 2 | 2 | 5 | 2 | 0 | 0 | 2 |
| | BB668AC | 3 | 2 | 1 | 6 | 1 | 3 | 3 | 7 |
| | BB785AE | 2 | 3 | 2 | 7 | 4 | 3 | 1 | 8 |
| | BB512A | 1 | 2 | 3 | 6 | 1 | 0 | 1 | 2 |
| Group5 DH1041 | BA507HA | 1 | 0 | 1 | 2 | 0 | 0 | 0 | 0 |
| | CB618B | 0 | 1 | 0 | 1 | 0 | 0 | 0 | 0 |
| | AP275E | 1 | 0 | 1 | 2 | 0 | 0 | 0 | 0 |
| | BB122A | 1 | 2 | 2 | 5 | 0 | 0 | 0 | 0 |
| | CDC063 | 2 | 1 | 1 | 4 | 0 | 0 | 0 | 0 |
| Group6 DH1052 | BB518A | 2 | 1 | 2 | 5 | 0 | 0 | 0 | 0 |
| | BB536A | 4 | 3 | 3 | 10 | 1 | 0 | 0 | 1 |
| | BC528A | 3 | 2 | 2 | 7 | 0 | 0 | 0 | 0 |
| | BC607D | 0 | 1 | 1 | 2 | 0 | 0 | 0 | 0 |
| | 2003AM | 1 | 2 | 1 | 4 | 0 | 0 | 0 | 0 |
| Group7 DH1047 | CZ605 | 1 | 1 | 2 | 4 | 0 | 0 | 0 | 0 |
| | DA014 | 2 | 1 | 0 | 3 | 0 | 0 | 0 | 0 |
| | SA10J | 0 | 1 | 0 | 1 | 0 | 0 | 0 | 0 |
| | SA56K | 0 | 0 | 0 | 0 | 0 | 0 | 0 | 0 |
| | SZ201 | 0 | 1 | 2 | 3 | 0 | 0 | 1 | 1 |

**Figure S22. Pathology scoring of the non-human primates that treated with RBD or NTD NAbs and infected with SARS-CoV-2 (Related to Figure 5H-I and 6F-G).** Lung histopathology. Sections of the left caudal (Lc), right middle (Rm), and right caudal (Rc) lung were evaluated histologically and scored using a scale of 1-4, assessing sections for the presence of inflammation by hematoxylin and eosin (H&E) staining, and for the presence of SARS-CoV-2 antigen by Immunohistochemistry (IHC) staining. The sums of scores for Lc, Rm, and Rc sections were shown in the table and shown as dot plots in Figure 5-6. **While individual animals from Groups 3 (2019B and BF791E) and Group 6 (BB536A) had cumulative lung pathology severity scores totaling 8-10, the characteristics of the inflammatory changes for BB536A were nuanced by the presence of perivascular and intralaveolar edema.*

### Figure S23

**A**

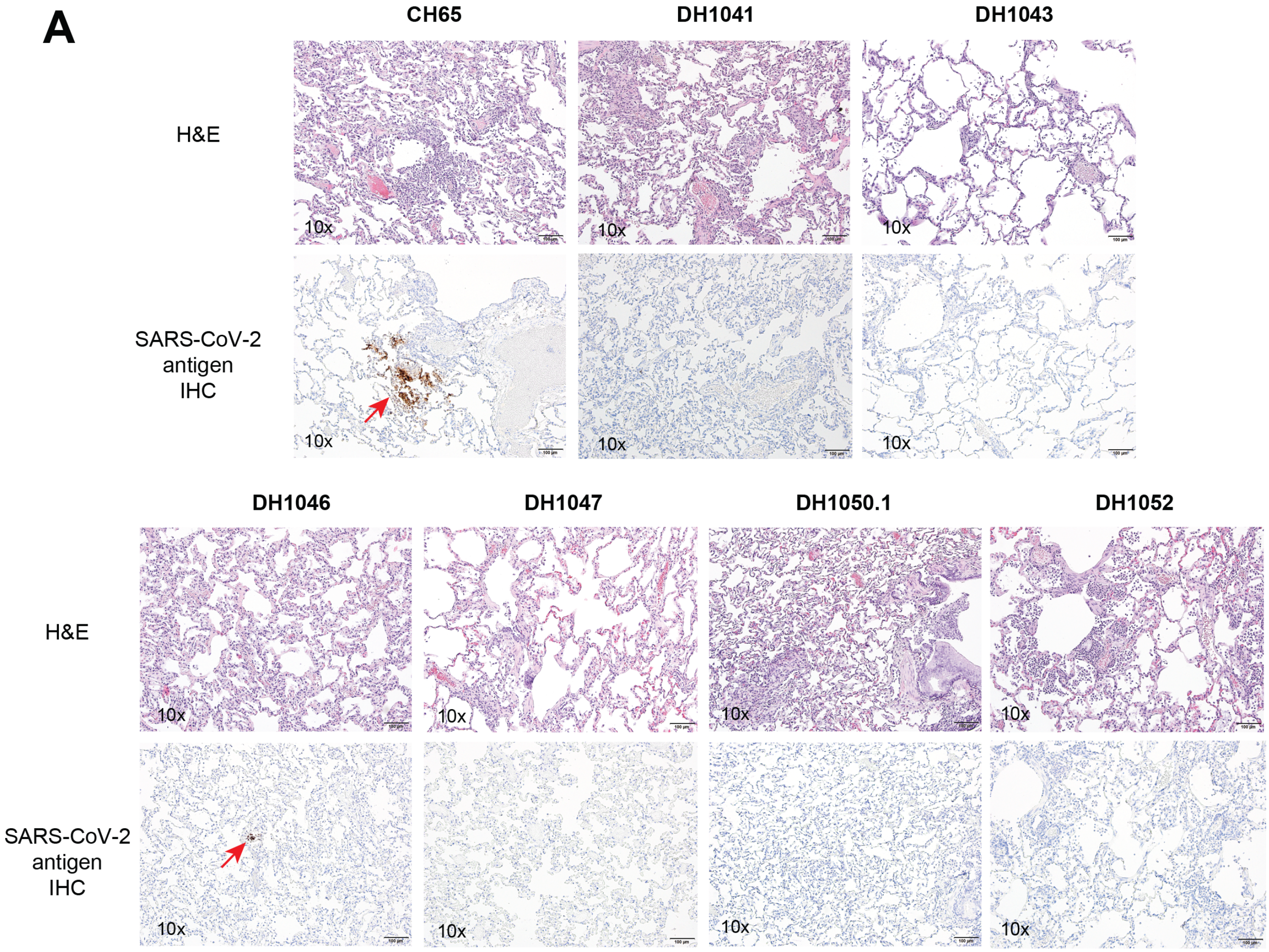

**B**

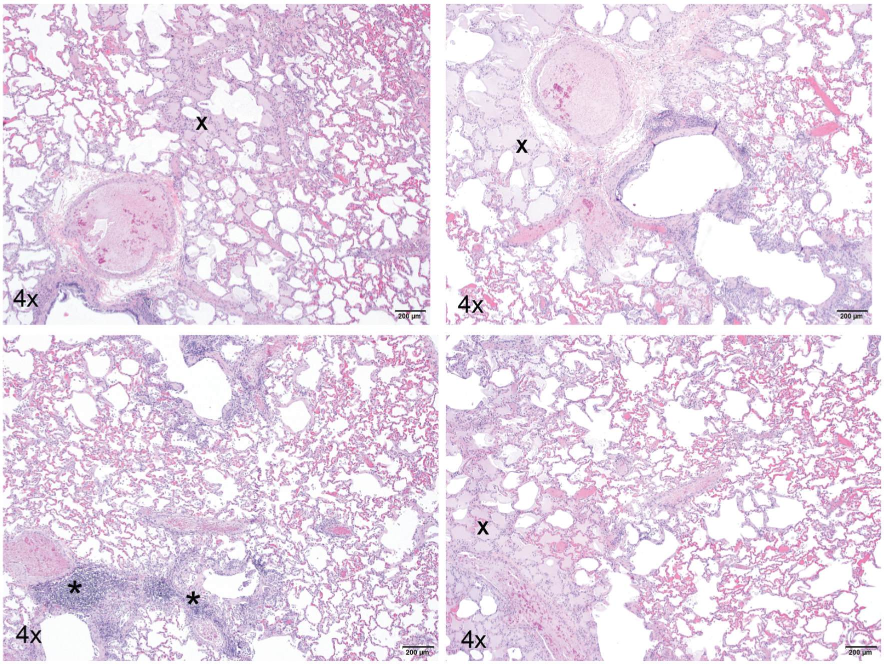

**Figure S23. Lung histopathology of antibody-treated and SARS-CoV-2 challenged cynomolgus macaques.**
**(A)** Representative images of hematoxylin and eosin (H&E) staining and SARS-CoV-2 antigen immunohistochemistry (IHC) staining from each group. All images were taken at 10x magnification. The images in this presentation are representative of the average severity of pathologic processes observed and recorded during microscopic evaluation. Red arrows indicate SARS-CoV-2 infection foci.
**(B)** Following microscopic evaluation of DH1052, 1 animal (BB536A) out of 5 animals in this group exhibited histologic features that was substantially more severe than the rest of the cohort and may suggest some degree of antibody-mediated disease enhancement. The features were characterized by prominent perivascular mononuclear inflammation (*) and a substantial amount of perivascular and alveolar edema (fluid; X). These findings suggest a vaso-centric process with some degree of altered vascular permeability. The remaining 4 animals in DH1052 group had inflammatory changes that ranged from minimal to moderate severity and more infiltrates were mixed and predominantly polymorphonuclear with lesser mononuclear cell involvement and present in the alveolar spaces.

### Figure S24

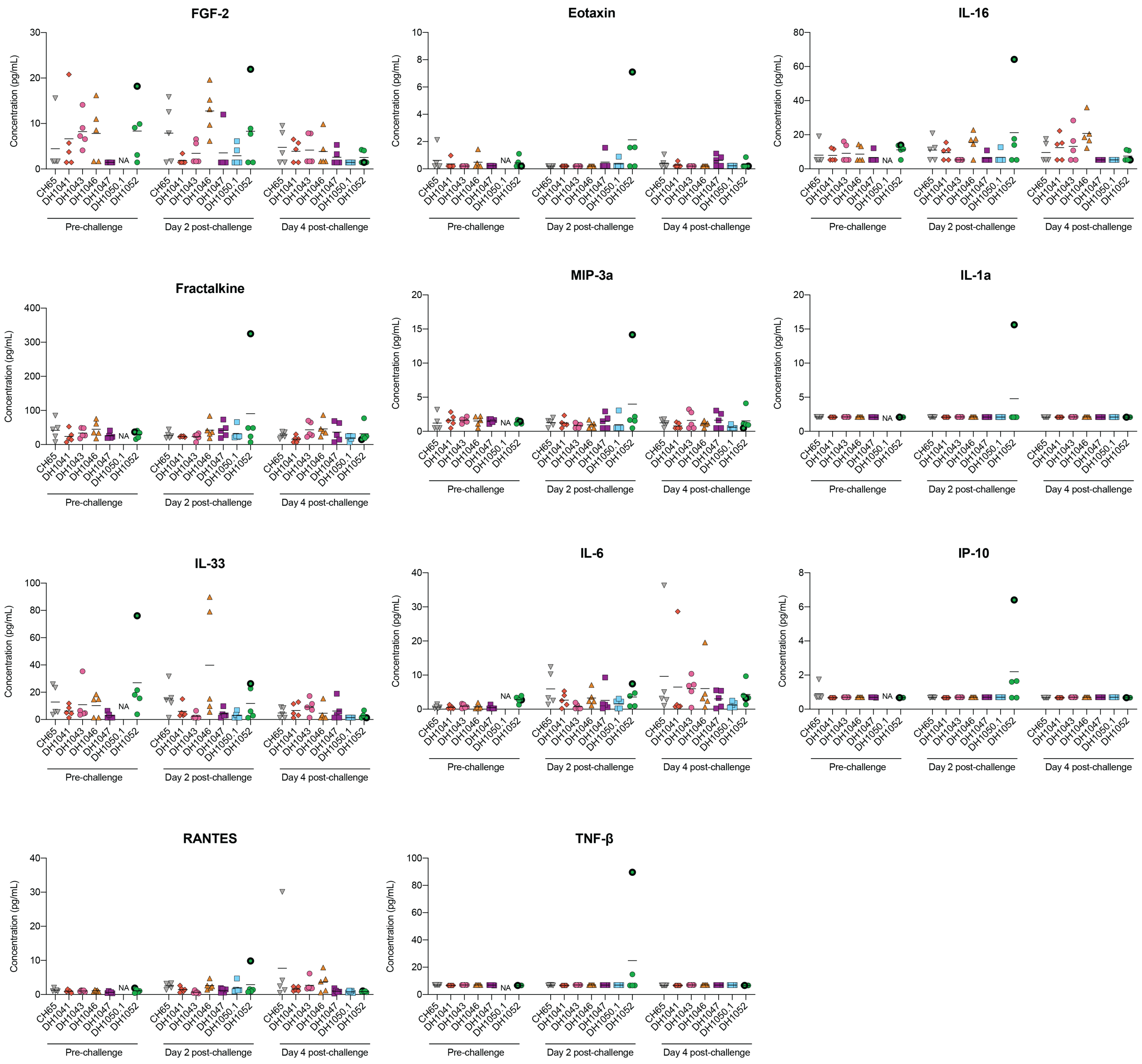

**Figure S24. Luminex profiling of cynomolgus macaques BAL samples for inflammatory cytokines (FGF-2, Eotaxin, IL-16, Fractalkine, MIP-3a, IL-1a, IL-33, IL-6, IP-10, RANTES, TNF-β).** BAL samples collected on Day -5 (pre-challenge), Day 2 and Day 4 post-challenge were concentrated (x7) and measured using a 25-analyte multiplex bead array by Luminex assay. The one animal (BB536A) in DH1052 group that exhibited substantially more severe histologic features than the rest of the cohort was labeled in black bold border.

### Figure S25

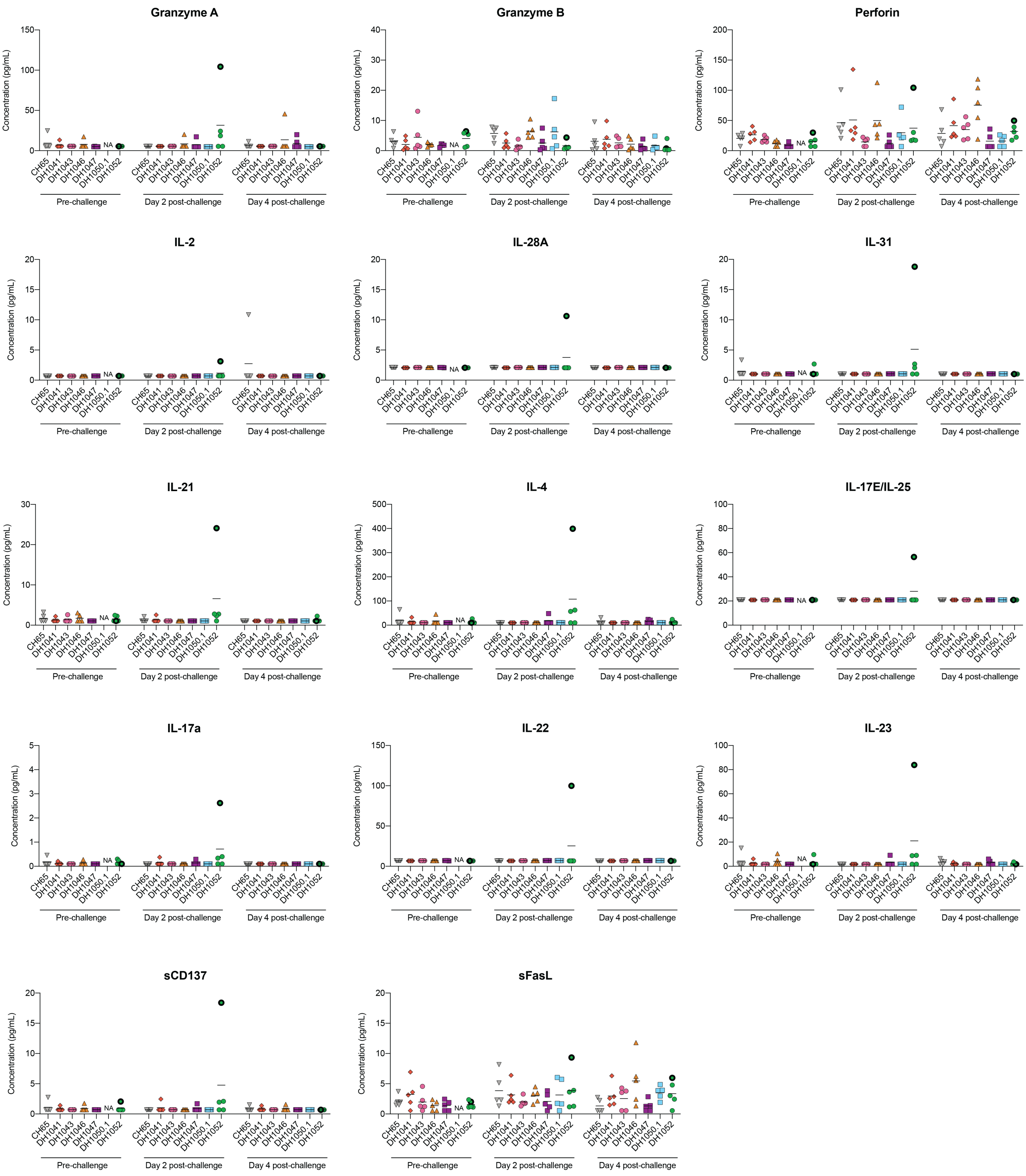

**Figure S25. Luminex profiling of cynomolgus macaques BAL samples for Th1, Th2, Th17 and other cytokines (Granzyme A, Granzyme B, Perforin, IL-2, IL-28a, IL-31, IL-21, IL-4, IL-17E/IL-25, IL17a, IL-22, IL-23, sCD137 and sFasL).** BAL samples collected on Day -5 (pre-challenge), Day 2 and Day 4 post-challenge were concentrated (x7) and measured using a 25-analyte multiplex bead array by Luminex assay. The one animal (BB536A) in DH1052 group that exhibited substantially more severe histologic features than the rest of the cohort was labeled in black bold border.

### Figure S26

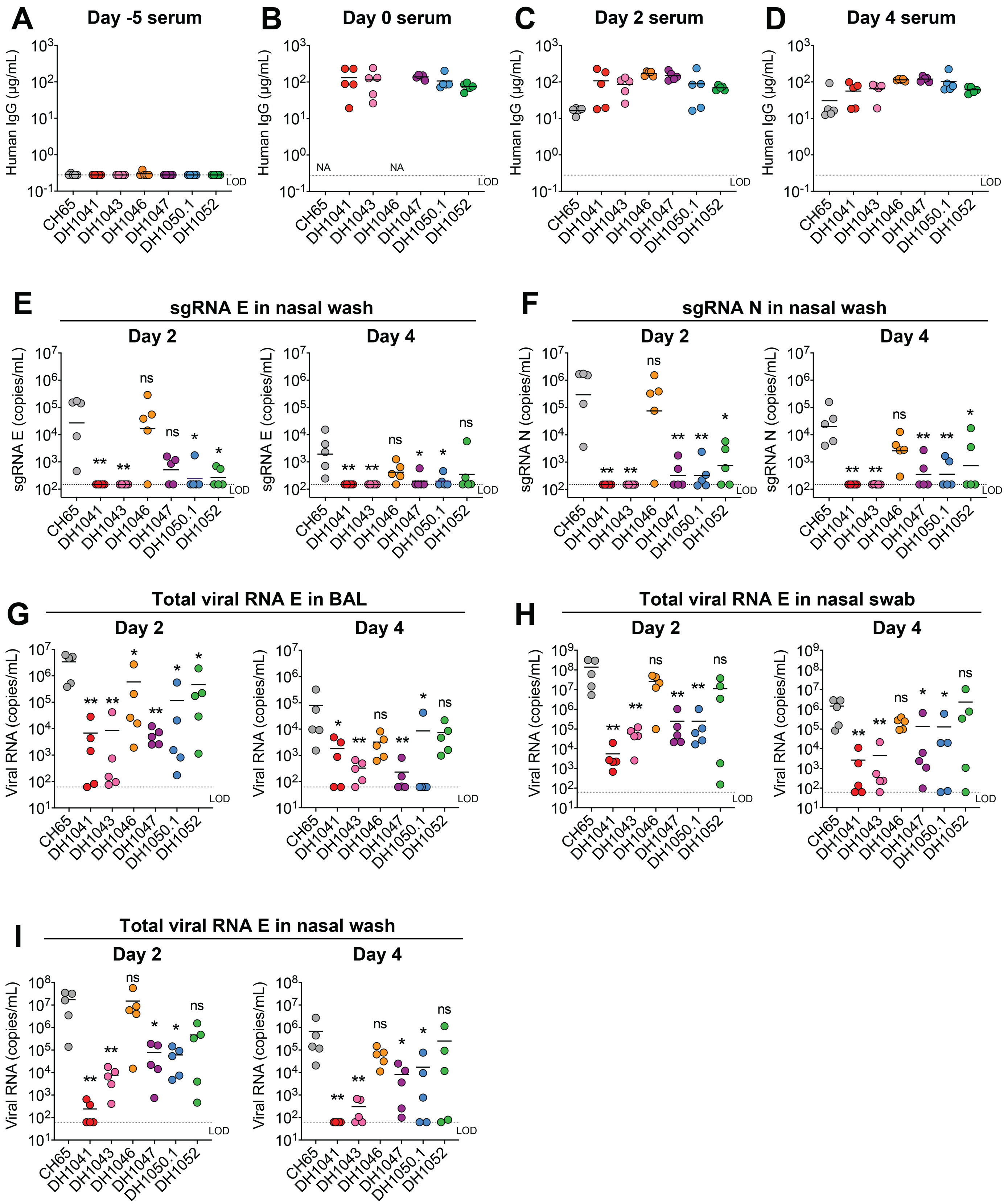
A
Day -5 serum
B
Day 0 serum
C
Day 2 serum
D
Day 4 serum
Human IgG (µg/mL)
NA
LOD
CH65
DH1041
DH1043
DH1046
DH1047
DH1050.1
DH1052
E
sgRNA E in nasal wash
Day 2
Day 4
sgRNA E (copies/mL)
F
sgRNA N in nasal wash
sgRNA N (copies/mL)
G
Total viral RNA E in BAL
H
Total viral RNA E in nasal swab
I
Total viral RNA E in nasal wash
Viral RNA (copies/mL)
ns
*
**

**Figure S26. SARS-CoV-2 total viral RNA and subgenomic RNA (sgRNA) in non-human primates that treated with RBD or NTD NAbs and infected with SARS-CoV-2 (Related to Figure 5 and 6).** Cynomolgus macaques (n=5 per group) were infused with RBD or NTD neutralizing antibodies 3 days before $10^5$ PFU of SARS-CoV-2 challenge. An irrelevant human antibody CH65 was used as a negative control. Viral load including viral RNA and subgenomic RNA (sgRNA) were measured on the indicated pre-challenge and post-challenge timepoints. Serum human IgG were quantified by Luminex assay. Statistical significance in all the panels were determined using Wilcoxon rank sum exact test (ns, no significance, *P<0.05, **P<0.001, ***P<0.0001). NA, samples not available. LOD, limit of detection.

**(A-D)** Serum human IgG concentrations at Day -5 (A), Day 0 (B), Day 2 (C), and Day 4 (D) post-challenge.

**(E-F)** SARS-CoV-2 (E) envelope gene (E gene) sgRNA and (F) nucleocapsid gene (N gene) sgRNA in nasal wash samples on Day 2 and Day 4 post challenge.

**(G)** SARS-CoV-2 E gene total (genomic + subgenomic) viral RNA in BAL samples on Day 2 and Day 4 post challenge.

**(H)** SARS-CoV-2 E gene total (genomic + subgenomic) viral RNA in nasal swab samples on Day 2 and Day 4 post challenge.

**(I)** SARS-CoV-2 E gene total (genomic + subgenomic) viral RNA in nasal wash samples on Day 2 and Day 4 post challenge.

### Figure S27

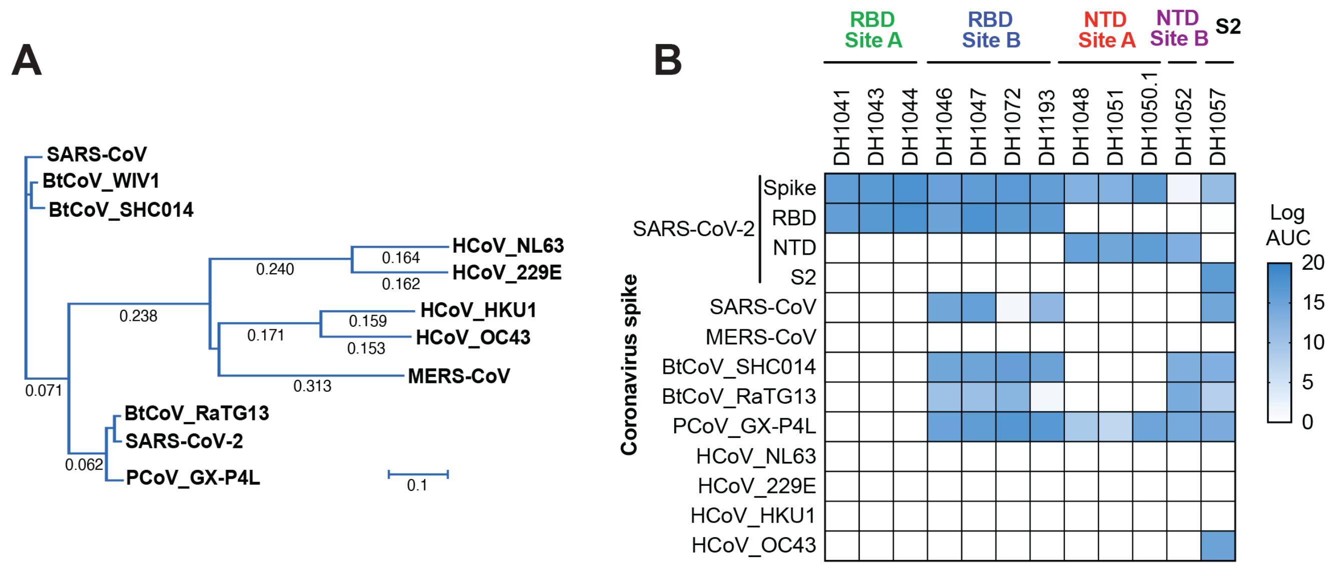

C

| Antibody | Epitope | Cross-reactivity | | | | | Cross-neutralization | | | |
|---|---|---|---|---|---|---|---|---|---|---|
| | | SARS | GXP4L | SHC014 | RaTG13 | OC43 | SARS-CoV-1 $IC_{50}$ | SARS-CoV-1 $IC_{80}$ | Bat CoV WIV-1 $IC_{50}$ | Bat CoV WIV-1 $IC_{80}$ |
| DH1046 | RBD | | | | | | 0.316 | 0.347 | 0.098 | 2.202 |
| DH1047 | RBD | | | | | | 0.154 | 0.169 | 0.073 | 1.120 |
| DH1072 | RBD | | | | | | >50 | >50 | 0.610 | 5.553 |
| DH1193 | RBD | | | | | | 19.160 | 31.400 | 3.605 | 28.054 |
| DH1057 | S2 | | | | | | >50 | >50 | 9.618 | 47.777 |

Binding
Non-reactive
Reactive
Neutralization titer (μg/mL)
<0.01
0.01-0.1
0.1-1
1-10
10-50
>50

**Figure S27. Cross-neutralizing RBD antibodies against SARS-CoV-1 and SARS-related bat WIV1-CoV.**
**(A)** Maximum likelihood tree of Spike amino acid sequences for SARS-related group 2B and group 2C coronaviruses.
**(B)** Monoclonal RBD, NTD and S2 antibody ELISA binding titer for soluble S protein ectodomains from human and animal SARS-related coronaviruses, human circulating coronaviruses and MERS. Titers are log area-under-the-curve (AUC).
**(C)** Summary of cross-neutralizing antibodies against SARS-CoV and/or bat WIV1-CoV.

**Table S1. High-throughput ELISA binding screen for the antibodies recovered from SARS-CoV-2 and SARS-CoV-1 donors.**

**Available online as excel sheet format.**

### Table S2. COVID-19 RBD neutralizing antibodies (n=44)

| DH# | Binding Specificity | Cross Reactivity | ACE-2 Blocking | | Neutralization | | | | | |
|---|---|---|---|---|---|---|---|---|---|---|
| | | | | | Pseudovirus | | | SARS-CoV-2 virus | | |
| | | | IC50 (µg/ml) | IC80 (µg/ml) | IC50 (µg/ml) | IC80 (µg/ml) | MPI | MN titer(µg/ml) | IC50 (µg/ml) | IC80 (µg/ml) |
| DH1043 | RBD | No | 0.043 | 0.172 | 0.0015 | 0.020 | 100 | <0.098 | 0.034 | 0.099 |
| DH1125 | RBD | No | <0.2 | 0.300 | 0.009 | 0.056 | 100 | 0.39 | 0.073 | 0.182 |
| DH1042 | RBD | No | 0.059 | 0.195 | 0.011 | 0.053 | 100 | 0.28 | 0.071 | 0.269 |
| DH1179 | RBD | No | <0.2 | 0.393 | 0.012 | 0.110 | 100 | 0.55 | 0.120 | 0.417 |
| DH1184 | RBD | No | <0.2 | 0.488 | 0.013 | 0.052 | 100 | 0.55 | 0.064 | 0.225 |
| DH1041 | RBD | No | 0.036 | 0.111 | 0.017 | 0.049 | 100 | <0.098 | 0.015 | 0.063 |
| DH1186 | RBD | No | <0.2 | 0.464 | 0.020 | 0.070 | 100 | 0.55 | 0.081 | 0.305 |
| DH1044 | RBD | No | >50 | >50 | 0.021 | 0.080 | 98 | 0.55 | 0.076 | 0.273 |
| DH1213 | RBD | No | 0.271 | 2.171 | 0.023 | 0.108 | 99 | 8.84 | 0.033 | 0.109 |
| DH1126 | RBD | No | <0.2 | 0.859 | 0.023 | 0.110 | 100 | 1.56 | 0.178 | 0.675 |
| DH1161.1 | RBD | No | 0.635 | 4.578 | 0.024 | 0.320 | 98 | 0.78 | 0.110 | 0.958 |
| DH1173 | RBD | No | 0.312 | 1.471 | 0.027 | 0.165 | 98 | 0.39 | 0.138 | 0.579 |
| DH1143 | RBD | No | 0.444 | 5.450 | 0.063 | 0.810 | 100 | 2.21 | 0.272 | 1.078 |
| DH1154 | RBD | No | 0.239 | 8.510 | 0.077 | 0.540 | 99 | 0.78 | 0.132 | 0.966 |
| DH1047 | RBD | SARS-CoV-1 | 0.069 | 0.315 | 0.090 | 0.360 | 100 | 1.10 | 0.124 | 0.666 |
| DH1151 | RBD | No | 22.647 | >50 | 0.100 | 0.400 | 100 | 1.10 | 0.367 | 1.587 |
| DH1197 | RBD | No | 0.356 | 1.371 | 0.170 | 1.000 | 100 | 4.42 | 0.987 | 2.330 |
| DH1160 | RBD | No | 4.159 | 34.271 | 0.180 | 1.690 | 94 | 3.13 | 0.335 | 4.749 |
| DH1140 | RBD | No | 1.671 | 49.444 | 0.250 | 1.750 | 96 | 12.50 | 3.167 | 12.220 |
| DH1210 | RBD | No | >50 | >50 | 0.280 | 2.290 | 99 | 8.84 | 5.334 | 23.660 |
| DH1150 | RBD | No | >50 | >50 | 0.320 | 3.720 | 99 | 17.68 | 8.800 | 10.800 |
| DH1045 | RBD | SARS-CoV-1 | 0.226 | 24.353 | 0.380 | 2.260 | 100 | 6.25 | 1.437 | 4.827 |
| DH1046 | RBD | SARS-CoV-1 | 0.170 | 1.274 | 0.396 | 2.030 | 100 | 12.50 | 1.086 | 9.383 |
| DH1180 | RBD | No | >50 | >50 | 0.420 | 1.930 | 100 | 35.36 | 8.969 | 11.490 |
| DH1183 | RBD | No | >50 | >50 | 0.550 | 7.560 | 96 | >100 | ND | ND |
| DH1128 | RBD | No | >50 | >50 | 0.703 | 4.685 | 95 | 50.00 | ND | ND |
| DH1129 | RBD | No | 15.226 | >50 | 0.970 | 6.410 | 95 | 17.68 | ND | ND |
| DH1155 | RBD | No | 16.309 | >50 | 1.060 | 6.030 | 99 | 25.00 | 7.904 | 13.670 |
| DH1193 | RBD | SARS-CoV-1 | >50 | >50 | 1.170 | 5.020 | 96 | 17.68 | 0.999 | 3.101 |
| DH1111 | RBD | No | >50 | >50 | 1.2000 | 10.3000 | 98 | 12.50 | 3.686 | 12.630 |
| DH1060 | RBD | No | >50 | >50 | 1.708 | 16.674 | 86 | 8.84 | 1.985 | 16.590 |
| DH1161.2 | RBD | No | >50 | >50 | 2.790 | 19.490 | 90 | 12.50 | 12.540 | 57.340 |
| DH1064 | RBD | SARS-CoV-1 | >50 | >50 | 4.360 | 23.800 | 92 | 70.71 | ND | ND |
| DH1073 | RBD | SARS-CoV-1 | >50 | >50 | 6.790 | 43.130 | 82 | 25.00 | 3.218 | 14.510 |
| DH1130 | RBD | ND | >50 | >50 | 7.614 | 36.512 | 86 | 35.40 | 8.113 | 22.800 |
| DH1063 | RBD | No | >50 | >50 | 9.199 | 44.588 | 82 | >100 | ND | ND |
| DH1082 | RBD | ND | >50 | >50 | 10.410 | >50 | 71 | 50.00 | 12.790 | 55.170 |
| DH1072 | RBD | SARS-CoV-1 | >50 | >50 | 11.7000 | 40.2000 | 85 | 17.68 | 3.540 | 28.660 |
| DH1159.2 | RBD | No | >50 | >50 | 11.900 | >50 | 77 | 100.00 | ND | ND |
| DH1088 | RBD | SARS-CoV-1 | >50 | >50 | 14.330 | 46.580 | 82 | 100.00 | ND | ND |
| DH1196 | RBD | HKU1 | >50 | >50 | 21.261 | >50 | 57 | 25.00 | 66.450 | >100 |
| DH1117 | RBD | SARS-CoV-1 | >50 | >50 | 22.560 | >50 | 65 | 100.00 | ND | ND |
| DH1156 | RBD | No | >50 | >50 | 26.520 | >50 | 58 | >100 | ND | ND |
| DH1214 | RBD | SARS-CoV-1 | >50 | >50 | >50 | >50 | 11.000 | 0.39 | >100 | >100 |

#### Table S3. COVID-19 RBD non-neutralizing antibodies (n=37)

| DH# | Binding Specificity | Cross Reactivity | ACE-2 Blocking | | Neutralization | | | | | |
|---|---|---|---|---|---|---|---|---|---|---|
| | | | | | Pseudovirus | | | SARS-CoV-2 virus | | |
| | | | IC50 (μg/ml) | IC80 (μg/ml) | IC50 (μg/ml) | IC80 (μg/ml) | MPI | MN titer(μg/ml) | IC50 (μg/ml) | IC80 (μg/ml) |
| DH1192 | RBD | No | >50 | >50 | >50 | >50 | 20 | >100 | ND | ND |
| DH1062 | RBD | No | >50 | >50 | ND | ND | ND | >100 | ND | ND |
| DH1091 | RBD | No | ND | ND | >50 | >50 | 1 | >100 | ND | ND |
| DH1102 | RBD | No | ND | ND | >50 | >50 | 20 | >100 | ND | ND |
| DH1139 | RBD | No | ND | ND | >50 | >50 | 26 | >100 | ND | ND |
| DH1169 | RBD | No | >50 | >50 | ND | ND | ND | 100.00 | ND | ND |
| DH1172 | RBD | SARS-CoV-1 | >50 | >50 | ND | ND | ND | 100.00 | ND | ND |
| DH1152 | RBD | SARS-CoV-1 | >50 | >50 | ND | ND | ND | >100 | ND | ND |
| DH1096 | RBD | No | >50 | >50 | ND | ND | ND | 50.00 | ND | ND |
| DH1104.2 | RBD | No | >50 | >50 | ND | ND | ND | >100 | ND | ND |
| DH1105.1 | RBD | No | >50 | >50 | ND | ND | ND | >100 | ND | ND |
| DH1109 | RBD | SARS-CoV-1 | >50 | >50 | ND | ND | ND | >100 | ND | ND |
| DH1127 | RBD | SARS-CoV-1 | >50 | >50 | ND | ND | ND | 100.00 | ND | ND |
| DH1185 | RBD | No | >50 | >50 | ND | ND | ND | >100 | ND | ND |
| DH1191 | RBD | SARS-CoV-1 | >50 | >50 | ND | ND | ND | >100 | ND | ND |
| DH1166 | RBD | SARS-CoV-1 | >50 | >50 | ND | ND | ND | 50.00 | ND | ND |
| DH1105.2 | RBD | No | >50 | >50 | ND | ND | ND | >100 | ND | ND |
| DH1203 | RBD | SARS-CoV-1 | >50 | >50 | ND | ND | ND | 50.00 | ND | ND |
| DH1170 | RBD | No | >50 | >50 | ND | ND | ND | >100 | ND | ND |
| DH1201 | RBD | No | >50 | >50 | ND | ND | ND | >100 | ND | ND |
| DH1209 | RBD | No | >50 | >50 | ND | ND | ND | >100 | ND | ND |
| DH1208 | RBD | SARS-CoV-1 | >50 | >50 | ND | ND | ND | >100 | ND | ND |
| DH1085 | RBD | SARS-CoV-1 | >50 | >50 | >50 | >50 | 47 | 141.42 | ND | ND |
| DH1075 | RBD | No | >50 | >50 | >50 | >50 | 14 | >100 | ND | ND |
| DH1080 | RBD | SARS-CoV-1 | >50 | >50 | >50 | >50 | 30 | >100 | ND | ND |
| DH1074 | RBD | No | >50 | >50 | >50 | >50 | 30 | >100 | ND | ND |
| DH1107 | RBD | No | >50 | >50 | >50 | >50 | 18 | >100 | ND | ND |
| DH1120 | RBD | SARS-CoV-1 | >50 | >50 | >50 | >50 | 0 | >100 | ND | ND |
| DH1092 | RBD | No | >50 | >50 | >50 | >50 | -1 | >100 | ND | ND |
| DH1112 | RBD | SARS-CoV-1 | >50 | >50 | >50 | >50 | -3 | >100 | ND | ND |
| DH1115 | RBD | SARS-CoV-1 | >50 | >50 | >50 | >50 | -4 | >100 | ND | ND |
| DH1095 | RBD | SARS-CoV-1 | 9.999 | >50 | >50 | >50 | -24 | >100 | ND | ND |
| DH1113 | RBD | SARS-CoV-1 | >50 | >50 | >50 | >50 | -7 | >100 | ND | ND |
| DH1116 | RBD | SARS-CoV-1 | >50 | >50 | >50 | >50 | 5 | >100 | ND | ND |
| DH1098 | RBD | SARS-CoV-1 | >50 | >50 | >50 | >50 | 6 | >100 | ND | ND |
| DH1101 | RBD | SARS-CoV-1 | >50 | >50 | >50 | >50 | 42 | >100 | ND | ND |

### Table S4. COVID-19 NTD neutralizing antibodies (n=10)

| DH# | Specificity | Cross Reactivity | ACE-2 Blocking | | Neutralization | | | | | |
|---|---|---|---|---|---|---|---|---|---|---|
| | | | | | Pseudovirus | | | SARS-CoV-2 virus | | |
| | | | IC50 (µg/ml) | IC80 (µg/ml) | IC50 (µg/ml) | IC80 (µg/ml) | MPI | MN titer(µg/ml) | IC50 (µg/ml) | IC80 (µg/ml) |
| DH1049 | NTD | No | >50 | >50 | 0.038 | >50 | 57 | 0.39 | 0.385 | 3.539 |
| DH1050.1 | NTD | No | >50 | >50 | 0.039 | >50 | 62 | 0.78 | 0.161 | 0.614 |
| DH1051 | NTD | No | >50 | >50 | 0.049 | >50 | 68 | 0.78 | 0.134 | 0.737 |
| DH1167 | NTD | No | >50 | >50 | 0.114 | >50 | 67 | 17.68 | 0.160 | 0.614 |
| DH1190 | NTD | No | >50 | >50 | 0.180 | >50 | 68 | 1.56 | 0.193 | 1.598 |
| DH1050.2 | NTD | No | >50 | >50 | 0.280 | >50 | 67 | 0.78 | 0.087 | 1.187 |
| DH1138 | NTD | No | >50 | >50 | 0.281 | >50 | 65 | 1.56 | 0.251 | 0.916 |
| DH1048 | NTD | No | >50 | >50 | 0.520 | >50 | 72 | 0.39 | 0.608 | 2.232 |
| DH1205 | NTD | No | >50 | >50 | 6.600 | >50 | 57 | 1.56 | 0.169 | 0.556 |
| DH1171 | NTD | SARS-CoV-1 | >50 | >50 | 46.750 | >50 | 51 | >100 | ND | ND |

### Table S5. COVID-19 NTD non-neutralizing antibodies (n=31)

| DH# | Binding Specificity | Cross Reactivity | ACE-2 Blocking | | Neutralization | | | | | |
|---|---|---|---|---|---|---|---|---|---|---|
| | | | | | Pseudovirus | | | SARS-CoV-2 virus | | |
| | | | IC50 (µg/ml) | IC80 (µg/ml) | IC50 (µg/ml) | IC80 (µg/ml) | MPI | MN titer(µg/ml) | IC50 (µg/ml) | IC80 (µg/ml) |
| DH1198 | NTD | No | >50 | >50 | >50 | >50 | 27 | >100 | ND | ND |
| DH1070 | NTD | No | >50 | >50 | >50 | >50 | -4 | >100 | ND | ND |
| DH1204 | NTD | No | >50 | >50 | ND | ND | ND | >100 | ND | ND |
| DH1162 | NTD | No | >50 | >50 | ND | ND | ND | >100 | ND | ND |
| DH1199 | NTD | No | >50 | >50 | ND | ND | ND | >100 | ND | ND |
| DH1090 | NTD | No | >50 | >50 | ND | ND | ND | >100 | ND | ND |
| DH1118 | NTD | No | >50 | >50 | ND | ND | ND | >100 | ND | ND |
| DH1119 | NTD | No | >50 | >50 | ND | ND | ND | >100 | ND | ND |
| DH1200 | NTD | No | >50 | >50 | ND | ND | ND | >100 | ND | ND |
| DH1144 | NTD | No | >50 | >50 | ND | ND | ND | >100 | ND | ND |
| DH1153 | NTD | No | >50 | >50 | >50 | >50 | 43 | >100 | ND | ND |
| DH1206 | NTD | No | >50 | >50 | >50 | >50 | 34 | >100 | ND | ND |
| DH1052 | NTD | No | >50 | >50 | >50 | >50 | -148 | >100 | ND | ND |
| DH1053 | NTD | No | >50 | >50 | >50 | >50 | -99 | >100 | ND | ND |
| DH1054 | NTD | No | >50 | >50 | >50 | >50 | -63 | >100 | ND | ND |
| DH1055 | NTD | No | >50 | >50 | >50 | >50 | -106 | >100 | ND | ND |
| DH1056 | NTD | No | >50 | >50 | >50 | >50 | -56 | >100 | ND | ND |
| DH1065 | NTD | SARS-CoV-1 | >50 | >50 | >32.5 | >32.5 | 24 | 50.00 | ND | ND |
| DH1069 | NTD | SARS-CoV-1 | >50 | >50 | >50 | >50 | 13 | >100 | ND | ND |
| DH1110 | NTD | SARS-CoV-1 | >50 | >50 | >50 | >50 | 6 | >100 | ND | ND |
| DH1106 | NTD | SARS-CoV-1 | >50 | >50 | >50 | >50 | 22 | >100 | ND | ND |
| DH1093 | NTD | SARS-CoV-1 | >50 | >50 | >50 | >50 | 17 | >100 | ND | ND |
| DH1114 | NTD | SARS-CoV-1 | >50 | >50 | >50 | >50 | 27 | >100 | ND | ND |
| DH1059 | NTD | SARS-CoV-1 | >50 | >50 | >50 | >50 | 6 | >100 | ND | ND |
| DH1081 | NTD | SARS-CoV-1 | >50 | >50 | >50 | >50 | 13 | >100 | ND | ND |
| DH1061.1 | NTD | SARS-CoV-1 | >50 | >50 | >50 | >50 | 41 | >100 | ND | ND |
| DH1066 | NTD | SARS-CoV-1 | >50 | >50 | >50 | >50 | 29 | >100 | ND | ND |
| DH1067 | NTD | SARS-CoV-1 | >50 | >50 | >50 | >50 | 30 | >100 | ND | ND |
| DH1068 | NTD | SARS-CoV-1 | >50 | >50 | >50 | >50 | 33 | >100 | ND | ND |
| DH1086 | NTD | SARS-CoV-1 | >50 | >50 | >50 | >50 | 29 | >100 | ND | ND |
| DH1071 | NTD | SARS-CoV-1 | >50 | >50 | >50 | >50 | 15 | >100 | ND | ND |

### Table S6. COVID-19 S2 antibodies (n=65)

| DH# | Binding Specificity | Cross Reactivity | ACE-2 Blocking | | Neutralization | | | | | |
|---|---|---|---|---|---|---|---|---|---|---|
| | | | | | Pseudovirus | | | SARS-CoV-2 virus | | |
| | | | IC50 (µg/ml) | IC80 (µg/ml) | IC50 (µg/ml) | IC80 (µg/ml) | MPI | MN titer(µg/ml) | IC50 (µg/ml) | IC80 (µg/ml) |
| DH1057 | S2 | SARS-CoV-1, OC43 | >50 | >50 | >50 | >50 | 46 | 17.68 | 10.890 | 44.100 |
| DH1189.3 | S2 | SARS-CoV-1, OC43 | >50 | >50 | ND | ND | ND | >100 | ND | ND |
| DH1175 | S2 | No | ND | ND | >50 | >50 | -1 | >100 | ND | ND |
| DH1103 | S2 | No | >50 | >50 | >50 | >50 | -12 | >100 | ND | ND |
| DH1121 | S2 | No | >50 | >50 | >50 | >50 | -12 | >100 | ND | ND |
| DH1122.2 | S2 | SARS-CoV-1, MERS-CoV, 229E, NL63, HKU1, OC43 | >50 | >50 | >50 | >50 | 0 | >100 | ND | ND |
| DH1123 | S2 | SARS-CoV-1, HKU1, OC43 | >50 | >50 | >26 | >26 | 5 | >100 | ND | ND |
| DH1078 | S2 | SARS-CoV-1, HKU1, OC43 | >50 | >50 | >50 | >50 | 15 | >100 | ND | ND |
| DH1137.3 | S2 | SARS-CoV-1, OC43 | >50 | >50 | >50 | >50 | -16 | >100 | ND | ND |
| DH1077 | S2 | No | >50 | >50 | >50 | >50 | -2 | >100 | ND | ND |
| DH1104.1 | S2 | SARS-CoV-1 | ND | ND | >20 | >20 | 3 | >100 | ND | ND |
| DH1137.1 | S2 | OC43 | ND | ND | >50 | >50 | 23 | >100 | ND | ND |
| DH1099 | S2 | SARS-CoV-1 | ND | ND | >50 | >50 | 21 | >100 | ND | ND |
| DH1132 | S2 | No | >50 | >50 | >50 | >50 | -6 | >100 | ND | ND |
| DH1189.1 | S2 | SARS-CoV-1, OC43 | >50 | >50 | >50 | >50 | 11 | >100 | ND | ND |
| DH1089 | S2 | No | >50 | >50 | >50 | >50 | 6 | >100 | ND | ND |
| DH1135 | S2 | No | >50 | >50 | >50 | >50 | 5 | >100 | ND | ND |
| DH1174 | S2 | No | >50 | >50 | >50 | >50 | -21 | >100 | ND | ND |
| DH1058 | S2 | SARS-CoV-1, MERS-CoV, 229E, NL63, HKU1, OC43 | >50 | >50 | >43.5 | >43.5 | -13 | >100 | ND | ND |
| DH1137.2 | S2 | OC43 | >50 | >50 | >50 | >50 | 43 | >100 | ND | ND |
| DH1124 | S2 | SARS-CoV-1, HKU1, OC43 | >50 | >50 | >50 | >50 | 27 | >100 | ND | ND |
| DH1122.1 | S2 | SARS-CoV-1, MERS-CoV, 229E, NL63, HKU1, OC43 | >50 | >50 | >50 | >50 | 24 | >100 | ND | ND |
| DH1195 | S2 | SARS-CoV-1 | >50 | >50 | >50 | >50 | 5 | >100 | ND | ND |
| DH1142.1 | S2 | No | >50 | >50 | ND | ND | ND | >100 | ND | ND |
| DH1189.2 | S2 | SARS-CoV-1, MERS-CoV, OC43 | >50 | >50 | ND | ND | ND | >100 | ND | ND |
| DH1188 | S2 | No | ND | ND | ND | ND | ND | >100 | ND | ND |
| DH1211 | S2 | SARS-CoV-1 | ND | ND | ND | ND | ND | >100 | ND | ND |
| DH1136 | S2 | No | ND | ND | ND | ND | ND | >100 | ND | ND |
| DH1083 | S2 | No | ND | ND | ND | ND | ND | >100 | ND | ND |
| DH1094 | S2 | No | ND | ND | ND | ND | ND | >100 | ND | ND |
| DH1100 | S2 | No | ND | ND | ND | ND | ND | >100 | ND | ND |
| DH1131 | S2 | SARS-CoV-1 | ND | ND | ND | ND | ND | >100 | ND | ND |
| DH1133 | S2 | SARS-CoV-1 | ND | ND | ND | ND | ND | >100 | ND | ND |
| DH1134 | S2 | No | ND | ND | ND | ND | ND | >100 | ND | ND |
| DH1165 | S2 | SARS-CoV-1 | ND | ND | ND | ND | ND | >100 | ND | ND |
| DH1141 | S2 | No | ND | ND | ND | ND | ND | >100 | ND | ND |
| DH1157 | S2 | SARS-CoV-1, MERS-CoV, 229E, NL63, HKU1, OC43 | ND | ND | ND | ND | ND | >100 | ND | ND |
| DH1168 | S2 | SARS-CoV-1 | ND | ND | ND | ND | ND | >100 | ND | ND |
| DH1097 | S2 | SARS-CoV-1 | ND | ND | ND | ND | ND | >100 | ND | ND |
| DH1159.2 | S2 | SARS-CoV-1, MERS-CoV, 229E, NL63, HKU1, OC43 | ND | ND | ND | ND | ND | >100 | ND | ND |
| DH1159.1 | S2 | SARS-CoV-1, MERS-CoV, 229E, NL63, HKU1, OC43 | ND | ND | ND | ND | ND | >100 | ND | ND |
| DH1163 | S2 | SARS-CoV-1, MERS-CoV | ND | ND | ND | ND | ND | >100 | ND | ND |
| DH1187 | S2 | SARS-CoV-1, MERS-CoV, 229E, NL63, HKU1, OC43 | ND | ND | ND | ND | ND | >100 | ND | ND |
| DH1194 | S2 | SARS-CoV-1 | ND | ND | ND | ND | ND | >100 | ND | ND |
| DH1164 | S2 | SARS-CoV-1 | ND | ND | ND | ND | ND | >100 | ND | ND |
| DH1145 | S2 | No | ND | ND | ND | ND | ND | >100 | ND | ND |
| DH1146 | S2 | No | ND | ND | ND | ND | ND | >100 | ND | ND |
| DH1212 | S2 | SARS-CoV-1 | ND | ND | ND | ND | ND | >100 | ND | ND |
| DH1147 | S2 | No | ND | ND | ND | ND | ND | >100 | ND | ND |
| DH1148 | S2 | No | ND | ND | ND | ND | ND | >100 | ND | ND |
| DH1149 | S2 | SARS-CoV-1 | ND | ND | ND | ND | ND | >100 | ND | ND |
| DH1176 | S2 | SARS-CoV-1 | ND | ND | ND | ND | ND | >100 | ND | ND |
| DH1177 | S2 | No | ND | ND | ND | ND | ND | >100 | ND | ND |
| DH1178 | S2 | No | ND | ND | ND | ND | ND | >100 | ND | ND |
| DH1181 | S2 | No | ND | ND | ND | ND | ND | >100 | ND | ND |
| DH1182 | S2 | No | ND | ND | ND | ND | ND | >100 | ND | ND |
| DH1202 | S2 | SARS-CoV-1 | ND | ND | ND | ND | ND | >100 | ND | ND |
| DH1207 | S2 | SARS-CoV-1 | ND | ND | ND | ND | ND | >100 | ND | ND |
| DH1142.2 | S2 | SARS-CoV-1 | ND | ND | ND | ND | ND | >100 | ND | ND |
| DH1142.3 | S2 | No | ND | ND | ND | ND | ND | >100 | ND | ND |
| DH1142.4 | S2 | No | ND | ND | ND | ND | ND | >100 | ND | ND |
| DH1079 | S2 | No | >50 | >50 | >50 | >50 | -9 | >100 | ND | ND |
| DH1084 | S2 | No | >50 | >50 | >50 | >50 | 29 | >100 | ND | ND |
| DH1108 | S2 | No | >50 | >50 | >50 | >50 | 1 | >100 | ND | ND |
| DH1087 | S2 | No | >50 | >50 | >50 | >50 | 11 | >100 | ND | ND |

### Table S7. Cross-reactive COVID-19 antibodies (n=80)

| DH# | Binding Specificity | Cross Reactivity | ACE-2 Blocking | | Neutralization | | | | | | | | | |
|---|---|---|---|---|---|---|---|---|---|---|---|---|---|---|
| | | | | | Pseudovirus | | | SARS-CoV-2 virus | | | SARS-CoV-1 virus | | Bat CoV WIV-1 virus | |
| | | | IC50 (µg/ml) | IC80 (µg/ml) | IC50 (µg/ml) | IC80 (µg/ml) | MPI | MN titer(µg/ml) | IC50 (µg/ml) | IC80 (µg/ml) | IC50 (µg/ml) | IC80 (µg/ml) | IC50 (µg/ml) | IC80 (µg/ml) |
| DH1049 | NTD | BCoV RaTG13 | >50 | >50 | 0.038 | >50 | 57 | 0.39 | 0.385 | 3.539 | ND | ND | ND | ND |
| DH1050.1 | NTD | BCoV RaTG13 | >50 | >50 | 0.039 | >50 | 62 | 0.78 | 0.161 | 0.614 | ND | ND | ND | ND |
| DH1051 | NTD | BCoV RaTG13 | >50 | >50 | 0.049 | >50 | 68 | 0.78 | 0.134 | 0.737 | ND | ND | ND | ND |
| DH1050.2 | NTD | BCoV RaTG13 | >50 | >50 | 0.280 | >50 | 67 | 0.78 | 0.087 | 1.187 | ND | ND | ND | ND |
| DH1048 | NTD | BCoV RaTG13 | >50 | >50 | 0.520 | >50 | 72 | 0.39 | 0.608 | 2.232 | ND | ND | ND | ND |
| DH1056 | NTD | BCoV RaTG13 | >50 | >50 | >50 | >50 | -56 | >100 | ND | ND | ND | ND | ND | ND |
| DH1054 | NTD | PCoV GXP4L, BCoV RaTG13 | >50 | >50 | >50 | >50 | -63 | >100 | ND | ND | ND | ND | ND | ND |
| DH1052 | NTD | PCoV GXP4L, BCoV RsSHC014, BCoV RaTG13 | >50 | >50 | >50 | >50 | -148 | >100 | ND | ND | ND | ND | ND | ND |
| DH1053 | NTD | PCoV GXP4L, BCoV RsSHC014, BCoV RaTG13 | >50 | >50 | >50 | >50 | -99 | >100 | ND | ND | ND | ND | ND | ND |
| DH1055 | NTD | PCoV GXP4L, BCoV RsSHC014, BCoV RaTG13 | >50 | >50 | >50 | >50 | -106 | >100 | ND | ND | ND | ND | ND | ND |
| DH1065 | NTD | SARS-CoV-1 | >50 | >50 | >32.5 | >32.5 | 24 | 50.00 | ND | ND | ND | ND | ND | ND |
| DH1069 | NTD | SARS-CoV-1 | >50 | >50 | >50 | >50 | 13 | >100 | ND | ND | ND | ND | ND | ND |
| DH1110 | NTD | SARS-CoV-1 | >50 | >50 | >50 | >50 | 6 | >100 | ND | ND | ND | ND | ND | ND |
| DH1106 | NTD | SARS-CoV-1 | >50 | >50 | >50 | >50 | 22 | >100 | ND | ND | ND | ND | ND | ND |
| DH1093 | NTD | SARS-CoV-1 | >50 | >50 | >50 | >50 | 17 | >100 | ND | ND | ND | ND | ND | ND |
| DH1114 | NTD | SARS-CoV-1 | >50 | >50 | >50 | >50 | 27 | >100 | ND | ND | ND | ND | ND | ND |
| DH1059 | NTD | SARS-CoV-1 | >50 | >50 | >50 | >50 | 6 | >100 | ND | ND | ND | ND | ND | ND |
| DH1081 | NTD | SARS-CoV-1 | >50 | >50 | >50 | >50 | 13 | >100 | ND | ND | ND | ND | ND | ND |
| DH1061 | NTD | SARS-CoV-1 | >50 | >50 | >50 | >50 | 41 | >100 | ND | ND | ND | ND | ND | ND |
| DH1066 | NTD | SARS-CoV-1 | >50 | >50 | >50 | >50 | 29 | >100 | ND | ND | ND | ND | ND | ND |
| DH1067 | NTD | SARS-CoV-1 | >50 | >50 | >50 | >50 | 30 | >100 | ND | ND | ND | ND | ND | ND |
| DH1068 | NTD | SARS-CoV-1 | >50 | >50 | >50 | >50 | 33 | >100 | ND | ND | ND | ND | ND | ND |
| DH1086 | NTD | SARS-CoV-1 | >50 | >50 | >50 | >50 | 29 | >100 | ND | ND | ND | ND | ND | ND |
| DH1071 | NTD | SARS-CoV-1 | >50 | >50 | >50 | >50 | 15 | >100 | ND | ND | ND | ND | ND | ND |
| DH1171 | NTD | SARS-CoV-1, BCoV RaTG13 | >50 | >50 | 46.750 | >50 | 51 | >100 | ND | ND | ND | ND | ND | ND |
| DH1196 | RBD | HKU1 | >50 | >50 | 21.261 | >50 | 57 | 25.00 | 66.450 | >100 | ND | ND | ND | ND |
| DH1172 | RBD | SARS-CoV-1 | >50 | >50 | ND | ND | ND | 100.00 | ND | ND | ND | ND | ND | ND |
| DH1214 | RBD | SARS-CoV-1 | >50 | >50 | >50 | >50 | 11 | 0.39 | >100 | >100 | ND | ND | ND | ND |
| DH1152 | RBD | SARS-CoV-1 | >50 | >50 | ND | ND | ND | >100 | ND | ND | ND | ND | ND | ND |
| DH1109 | RBD | SARS-CoV-1 | >50 | >50 | ND | ND | ND | >100 | ND | ND | ND | ND | ND | ND |
| DH1127 | RBD | SARS-CoV-1 | >50 | >50 | ND | ND | ND | 100.00 | ND | ND | ND | ND | ND | ND |
| DH1191 | RBD | SARS-CoV-1 | >50 | >50 | ND | ND | ND | >100 | ND | ND | ND | ND | ND | ND |
| DH1166 | RBD | SARS-CoV-1 | >50 | >50 | ND | ND | ND | 50.00 | ND | ND | ND | ND | ND | ND |
| DH1203 | RBD | SARS-CoV-1 | >50 | >50 | ND | ND | ND | 50.00 | ND | ND | ND | ND | ND | ND |
| DH1208 | RBD | SARS-CoV-1 | >50 | >50 | ND | ND | ND | >100 | ND | ND | ND | ND | ND | ND |
| DH1085 | RBD | SARS-CoV-1 | >50 | >50 | >50 | >50 | 47 | 141.42 | ND | ND | ND | ND | ND | ND |
| DH1088 | RBD | SARS-CoV-1 | >50 | >50 | 14.330 | 46.580 | 82 | 100.00 | ND | ND | ND | ND | ND | ND |
| DH1080 | RBD | SARS-CoV-1 | >50 | >50 | >50 | >50 | 30 | >100 | ND | ND | ND | ND | ND | ND |
| DH1064 | RBD | SARS-CoV-1 | >50 | >50 | 4.360 | 23.800 | 92 | 70.71 | ND | ND | ND | ND | ND | ND |
| DH1073 | RBD | SARS-CoV-1 | >50 | >50 | 6.790 | 43.130 | 82 | 25.00 | 3.218 | 14.510 | ND | ND | ND | ND |
| DH1120 | RBD | SARS-CoV-1 | >50 | >50 | >50 | >50 | 0 | >100 | ND | ND | ND | ND | ND | ND |
| DH1112 | RBD | SARS-CoV-1 | >50 | >50 | >50 | >50 | -3 | >100 | ND | ND | ND | ND | ND | ND |
| DH1117 | RBD | SARS-CoV-1 | >50 | >50 | 22.560 | >50 | 65 | 100.00 | ND | ND | ND | ND | ND | ND |
| DH1115 | RBD | SARS-CoV-1 | >50 | >50 | >50 | >50 | -4 | >100 | ND | ND | ND | ND | ND | ND |
| DH1095 | RBD | SARS-CoV-1 | 9.999 | >50 | >50 | >50 | -24 | >100 | ND | ND | ND | ND | ND | ND |
| DH1113 | RBD | SARS-CoV-1 | >50 | >50 | >50 | >50 | -7 | >100 | ND | ND | ND | ND | ND | ND |
| DH1116 | RBD | SARS-CoV-1 | >50 | >50 | >50 | >50 | 5 | >100 | ND | ND | ND | ND | ND | ND |
| DH1098 | RBD | SARS-CoV-1 | >50 | >50 | >50 | >50 | 6 | >100 | ND | ND | ND | ND | ND | ND |
| DH1101 | RBD | SARS-CoV-1 | >50 | >50 | >50 | >50 | 42 | >100 | ND | ND | ND | ND | ND | ND |
| DH1072 | RBD | SARS-CoV-1, PCoV GXP4L, BCoV RsSHC014, BCoV RaTG13 | >50 | >50 | 11.700 | 40.200 | 85 | 17.68 | 3.540 | 28.660 | >50 | >50 | 0.610 | 5.553 |
| DH1045 | RBD | SARS-CoV-1, PCoV GXP4L, BCoV RsSHC014, BCoV RaTG13 | 0.226 | 24.353 | 0.380 | 2.260 | 100 | 6.25 | 1.437 | 4.827 | >50 | >50 | >50 | >50 |
| DH1193 | RBD | SARS-CoV-1, PCoV GXP4L, BCoV RsSHC014, BCoV RaTG13 | >50 | >50 | 1.170 | 5.020 | 96 | 17.68 | 0.999 | 3.101 | 19.160 | 31.400 | 3.605 | 28.054 |
| DH1046 | RBD | SARS-CoV-1, PCoV GXP4L, BCoV RsSHC014, BCoV RaTG13 | 0.201 | 6.511 | 0.396 | 2.030 | 100 | 12.50 | 1.086 | 9.383 | 0.316 | 0.347 | 0.098 | 2.202 |
| DH1047 | RBD | SARS-CoV-1, PCoV GXP4L, BCoV RsSHC014, BCoV RaTG13 | 0.078 | 0.567 | 0.090 | 0.360 | 100 | 1.10 | 0.124 | 0.666 | 0.154 | 0.169 | 0.073 | 1.120 |
| DH1076 | S1 | SARS-CoV-1 specific | >50 | >50 | >50 | >50 | 15 | >100 | ND | ND | ND | ND | ND | ND |
| DH1137.1 | S2 | OC43 | ND | ND | >50 | >50 | 23 | >100 | ND | ND | ND | ND | ND | ND |
| DH1137.2 | S2 | OC43 | >50 | >50 | >50 | >50 | 43 | >100 | ND | ND | ND | ND | ND | ND |
| DH1104.1 | S2 | SARS-CoV-1 | ND | ND | >20 | >20 | 3 | >100 | ND | ND | ND | ND | ND | ND |
| DH1099 | S2 | SARS-CoV-1 | ND | ND | >50 | >50 | 21 | >100 | ND | ND | ND | ND | ND | ND |
| DH1195 | S2 | SARS-CoV-1 | >50 | >50 | >50 | >50 | 5 | >100 | ND | ND | ND | ND | ND | ND |
| DH1211 | S2 | SARS-CoV-1 | ND | ND | ND | ND | ND | >100 | ND | ND | ND | ND | ND | ND |
| DH1131 | S2 | SARS-CoV-1 | ND | ND | ND | ND | ND | >100 | ND | ND | ND | ND | ND | ND |
| DH1133 | S2 | SARS-CoV-1 | ND | ND | ND | ND | ND | >100 | ND | ND | ND | ND | ND | ND |
| DH1165 | S2 | SARS-CoV-1 | ND | ND | ND | ND | ND | >100 | ND | ND | ND | ND | ND | ND |
| DH1168 | S2 | SARS-CoV-1 | ND | ND | ND | ND | ND | >100 | ND | ND | ND | ND | ND | ND |
| DH1097 | S2 | SARS-CoV-1 | ND | ND | ND | ND | ND | >100 | ND | ND | ND | ND | ND | ND |
| DH1194 | S2 | SARS-CoV-1 | ND | ND | ND | ND | ND | >100 | ND | ND | ND | ND | ND | ND |
| DH1164 | S2 | SARS-CoV-1 | ND | ND | ND | ND | ND | >100 | ND | ND | ND | ND | ND | ND |
| DH1212 | S2 | SARS-CoV-1 | ND | ND | ND | ND | ND | >100 | ND | ND | ND | ND | ND | ND |
| DH1149 | S2 | SARS-CoV-1 | ND | ND | ND | ND | ND | >100 | ND | ND | ND | ND | ND | ND |
| DH1176 | S2 | SARS-CoV-1 | ND | ND | ND | ND | ND | >100 | ND | ND | ND | ND | ND | ND |
| DH1202 | S2 | SARS-CoV-1 | ND | ND | ND | ND | ND | >100 | ND | ND | ND | ND | ND | ND |
| DH1207 | S2 | SARS-CoV-1 | ND | ND | ND | ND | ND | >100 | ND | ND | ND | ND | ND | ND |
| DH1142.2 | S2 | SARS-CoV-1 | ND | ND | ND | ND | ND | >100 | ND | ND | ND | ND | ND | ND |
| DH1123 | S2 | SARS-CoV-1, HKU1, OC43 | >50 | >50 | >26 | >26 | 5 | >100 | ND | ND | ND | ND | ND | ND |
| DH1078 | S2 | SARS-CoV-1, HKU1, OC43 | >50 | >50 | >50 | >50 | 15 | >100 | ND | ND | ND | ND | ND | ND |
| DH1124 | S2 | SARS-CoV-1, HKU1, OC43 | >50 | >50 | >50 | >50 | 27 | >100 | ND | ND | ND | ND | ND | ND |
| DH1163 | S2 | SARS-CoV-1, MERS-CoV | ND | ND | ND | ND | ND | >100 | ND | ND | ND | ND | ND | ND |
| DH1121 | S2 | SARS-CoV-1, MERS-CoV, 229E, NL63, HKU1, OC43 | >50 | >50 | >50 | >50 | 0 | >100 | ND | ND | ND | ND | ND | ND |
| DH1122.1 | S2 | SARS-CoV-1, MERS-CoV, 229E, NL63, HKU1, OC43 | >50 | >50 | >50 | >50 | 24 | >100 | ND | ND | ND | ND | ND | ND |
| DH1157 | S2 | SARS-CoV-1, MERS-CoV, 229E, NL63, HKU1, OC43 | ND | ND | ND | ND | ND | >100 | ND | ND | ND | ND | ND | ND |
| DH1158.1 | S2 | SARS-CoV-1, MERS-CoV, 229E, NL63, HKU1, OC43 | ND | ND | ND | ND | ND | >100 | ND | ND | ND | ND | ND | ND |
| DH1158.2 | S2 | SARS-CoV-1, MERS-CoV, 229E, NL63, HKU1, OC43 | ND | ND | ND | ND | ND | >100 | ND | ND | ND | ND | ND | ND |
| DH1187 | S2 | SARS-CoV-1, MERS-CoV, 229E, NL63, HKU1, OC43 | ND | ND | ND | ND | ND | >100 | ND | ND | ND | ND | ND | ND |
| DH1058 | S2 | SARS-CoV-1, MERS-CoV, 229E, NL63, HKU1, OC43, PCoV GXP4L, BCoV RsSHC014, BCoV RaTG13 | >50 | >50 | >43.5 | >43.5 | -13 | >100 | ND | ND | ND | ND | ND | ND |
| DH1189.2 | S2 | SARS-CoV-1, MERS-CoV, OC43 | >50 | >50 | ND | ND | ND | >100 | ND | ND | ND | ND | ND | ND |
| DH1189.3 | S2 | SARS-CoV-1, OC43 | >50 | >50 | ND | ND | ND | >100 | ND | ND | ND | ND | ND | ND |
| DH1136 | S2 | SARS-CoV-1, OC43 | >50 | >50 | >50 | >50 | -16 | >100 | ND | ND | ND | ND | ND | ND |
| DH1189.1 | S2 | SARS-CoV-1, OC43 | >50 | >50 | >50 | >50 | 11 | >100 | ND | ND | ND | ND | ND | ND |
| DH1057 | S2 | SARS-CoV-1, OC43, PCoV GXP4L, BCoV RsSHC014, BCoV RaTG13 | >50 | >50 | >50 | >50 | 46 | 17.68 | 10.890 | 44.100 | >50 | >50 | 9.618 | 47.777 |

**Table S8. Immunogenetic analysis of select neutralizing and non-neutralizing SARS-CoV-2 antibodies.**

**Available online as excel sheet format.**
